# Subcortical Volume Trajectories across the Lifespan: Data from 18,605 healthy individuals aged 3-90 years

**DOI:** 10.1101/2020.05.05.079475

**Authors:** Danai Dima, Efstathios Papachristou, Amirhossein Modabbernia, Gaelle E Doucet, Ingrid Agartz, Moji Aghajani, Theophilus N Akudjedu, Anton Albajes-Eizagirre, Dag Alnæs, Kathryn I Alpert, Micael Andersson, Nancy Andreasen, Ole A Andreassen, Philip Asherson, Tobias Banaschewski, Nuria Bargallo, Sarah Baumeister, Ramona Baur-Streubel, Alessandro Bertolino, Aurora Bonvino, Dorret I Boomsma, Stefan Borgwardt, Josiane Bourque, Daniel Brandeis, Alan Breier, Henry Brodaty, Rachel M Brouwer, Jan K Buitelaar, Geraldo F Busatto, Randy L Buckner, Vincent Calhoun, Erick J Canales-Rodríguez, Dara M Cannon, Xavier Caseras, Francisco X Castellanos, Simon Cervenka, Tiffany M Chaim-Avancini, Christopher RK Ching, Vincent P Clark, Patricia Conrod, Annette Conzelmann, Benedicto Crespo-Facorro, Fabrice Crivello, Eveline AM Crone, Anders M Dale, Cristopher Davey, Eco JC de Geus, Lieuwe de Haan, Greig I de Zubicaray, Anouk den Braber, Erin W Dickie, Annabella Di Giorgio, Nhat Trung Doan, Erlend S Dørum, Stefan Ehrlich, Susanne Erk, Thomas Espeseth, Helena Fatouros-Bergman, Simon E Fisher, Jean-Paul Fouche, Barbara Franke, Thomas Frodl, Paola Fuentes-Claramonte, David C Glahn, Ian H Gotlib, Hans-Jörgen Grabe, Oliver Grimm, Nynke A Groenewold, Dominik Grotegerd, Oliver Gruber, Patricia Gruner, Rachel E Gur, Ruben C Gur, Ben J Harrison, Catharine A Hartman, Sean N Hatton, Andreas Heinz, Dirk J Heslenfeld, Derrek P Hibar, Ian B Hickie, Beng-Choon Ho, Pieter J Hoekstra, Sarah Hohmann, Avram J Holmes, Martine Hoogman, Norbert Hosten, Fleur M Howells, Hilleke E Hulshoff Pol, Chaim Huyser, Neda Jahanshad, Anthony James, Jiyang Jiang, Erik G Jönsson, John A Joska, Rene Kahn, Andrew Kalnin, Ryota Kanai, Sim Kang, Marieke Klein, Tatyana P Klushnik, Laura Koenders, Sanne Koops, Bernd Krämer, Jonna Kuntsi, Jim Lagopoulos, Luisa Lázaro, Irina Lebedeva, Won Hee Lee, Klaus-Peter Lesch, Christine Lochner, Marise WJ Machielsen, Sophie Maingault, Nicholas G Martin, Ignacio Martínez-Zalacaín, David Mataix-Cols, Bernard Mazoyer, Colm McDonald, Brenna C McDonald, Andrew M McIntosh, Katie L McMahon, Genevieve McPhilemy, José M Menchón, Sarah E Medland, Andreas Meyer-Lindenberg, Jilly Naaijen, Pablo Najt, Tomohiro Nakao, Jan E Nordvik, Lars Nyberg, Jaap Oosterlaan, Víctor Ortiz-García de la Foz, Yannis Paloyelis, Paul Pauli, Giulio Pergola, Edith Pomarol-Clotet, Maria J Portella, Steven G Potkin, Joaquim Radua, Andreas Reif, Joshua L Roffman, Pedro GP Rosa, Matthew D Sacchet, Perminder S Sachdev, Raymond Salvador, Pascual Sánchez-Juan, Salvador Sarró, Theodore D Satterthwaite, Andrew J Saykin, Mauricio H Serpa, Lianne Schmaal, Knut Schnell, Gunter Schumann, Jordan W Smoller, Iris Sommer, Carles Soriano-Mas, Dan J Stein, Lachlan T Strike, Suzanne C Swagerman, Christian K Tamnes, Henk S Temmingh, Sophia I Thomopoulos, Alexander S Tomyshev, Diana Tordesillas-Gutiérrez, Julian N Trollor, Jessica A Turner, Anne Uhlmann, Odille A van den Heuvel, Dennis van den Meer, Nic JA van der Wee, Neeltje EM van Haren, Dennis van ’t Ent, Theo GM van Erp, Ilya M Veer, Dick J Veltman, Henry Völzke, Henrik Walter, Esther Walton, Lei Wang, Yang Wang, Thomas H Wassink, Bernd Weber, Wei Wen, John D West, Lars T Westlye, Heather Whalley, Lara M Wierenga, Steven CR Williams, Katharina Wittfeld, Daniel H Wolf, Amanda Worker, Margaret J Wright, Kun Yang, Yulyia Yoncheva, Marcus V Zanetti, Georg C Ziegler, Paul M Thompson, Sophia Frangou

## Abstract

Age has a major effect on brain volume. However, the normative studies available are constrained by small sample sizes, restricted age coverage and significant methodological variability. These limitations introduce inconsistencies and may obscure or distort the lifespan trajectories of brain morphometry. In response, we capitalised on the resources of the Enhancing Neuroimaging Genetics through Meta-Analysis (ENIGMA) Consortium to examine the age-related morphometric trajectories of the ventricles, the basal ganglia (caudate, putamen, pallidum, and nucleus accumbens), the thalamus, hippocampus and amygdala using magnetic resonance imaging data obtained from 18,605 individuals aged 3-90 years. All subcortical structure volumes were at their maximum early in life; the volume of the basal ganglia showed a gradual monotonic decline thereafter while the volumes of the thalamus, amygdala and the hippocampus remained largely stable (with some degree of decline in thalamus) until the sixth decade of life followed by a steep decline thereafter. The lateral ventricles showed a trajectory of continuous enlargement throughout the lifespan. Significant age-related increase in inter-individual variability was found for the hippocampus and amygdala and the lateral ventricles. These results were robust to potential confounders and could be used to derive risk predictions for the early identification of diverse clinical phenotypes.

## Introduction

Over the last 20 years, studies using structural magnetic resonance imaging (MRI) have confirmed that brain morphometric measures changes with age. In general, whole brain, global and regional gray matter volumes increase during development and decrease with aging (Brain Development Cooperative Group, 2012; Driscoll et al. 2009; Fotenos et al. 2005; Good et al. 2001; Pfefferbaum et al. 2013; Pomponio et al., 2019; Raz et al. 2005; Raznahan et al. 2014; Resnick et al. 2003; Walhovd et al. 2011). However, most published studies are constrained by small sample sizes, restricted age coverage and methodological variability. These limitations introduce inconsistencies and may obscure or distort the lifespan trajectories of brain structures. To address these limitations, we formed the Lifespan Working group of the Enhancing Neuroimaging Genetics through Meta-Analysis (ENIGMA) Consortium (Thompson et al. 2014, 2017) to perform large-scale analyses of brain morphometric data extracted from MRI images using standardized protocols and unified quality control procedures, harmonized and validated across all participating sites.

Here we focus on ventricular, striatal (caudate, putamen, nucleus accumbens), pallidal, thalamic, hippocampal and amygdala volumes. Subcortical structures are crucial for normal cognitive and emotional adaptation (Grossberg, 2009). The striatum and pallidum (together referred to as basal ganglia) are best known for their role in action selection and movement coordination (Calabresi et al. 2014) but they are also involved in other aspects of cognition particularly memory, inhibitory control, reward and salience processing (Chudasama and Robbins 2006; Richard et al. 2013; Scimeca and Badre 2012; Tremblay et al. 2015). The role of the hippocampus has been most clearly defined in connection to declarative memory (Eichenbaum, 2004; Shohamy and Turk-Browne 2013) while the amygdala has been historically linked to affect processing (Kober et al. 2008). The thalamus is centrally located in the brain and acts as a key hub for the integration of motor and sensory information with higher-order functions (Sherman 2005; Zhang et al. 2010). The role of subcortical structures extends beyond normal cognition because changes in the volume of these regions have been reliably identified in developmental (Ecker et al. 2015; Krain and Castellanos 2006), psychiatric (Kempton et al. 2011; Hibar et al. 2016; Schmaal et al. 2016; van Erp et al. 2016) and degenerative disorders (Risacher et al. 2009).

Using data from 18,605 individuals aged 3-90 years from the ENIGMA Lifespan working group we delineated the age-related trajectories of subcortical volumes from early to late life in order to (a) identify periods of volume change or stability, (b) provide normative, age-adjusted centile curves of subcortical volumes and (c) quantify inter-individual variability in subcortical volumes which is considered a major source of inter-study differences in age-related trajectories derived from smaller samples (Dickie et al. 2013; Raz et al. 2010).

## Materials and Methods

### Study Samples

The study data comes from 88 samples and comprising 18,605 healthy participants, aged 3-90 years, with near equal representation of men and women (48% and 52%) (Table 1, Figure 1). At the time of scanning, participating individuals were screened to exclude the presence of mental disorders, cognitive impairment or significant medical morbidity. Details of the screening process and eligibility criteria for each research group are shown in Table S1).

**Figure 1.**
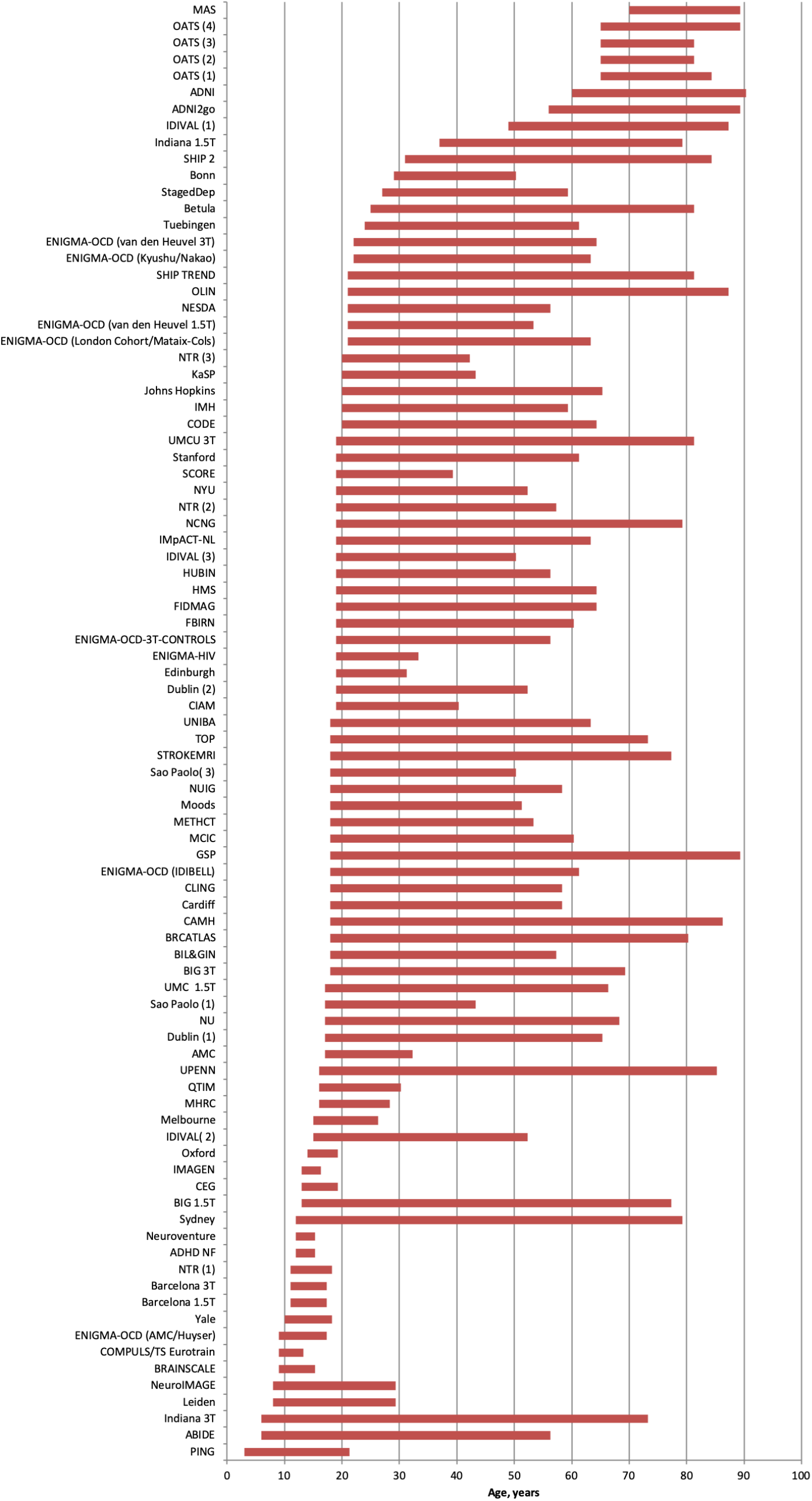
ENIGMA Lifespan Samples. Details of each sample are provided Table 1 and in the supplemental material. Abbreviations are provided in Table 1.

**Table 1.**
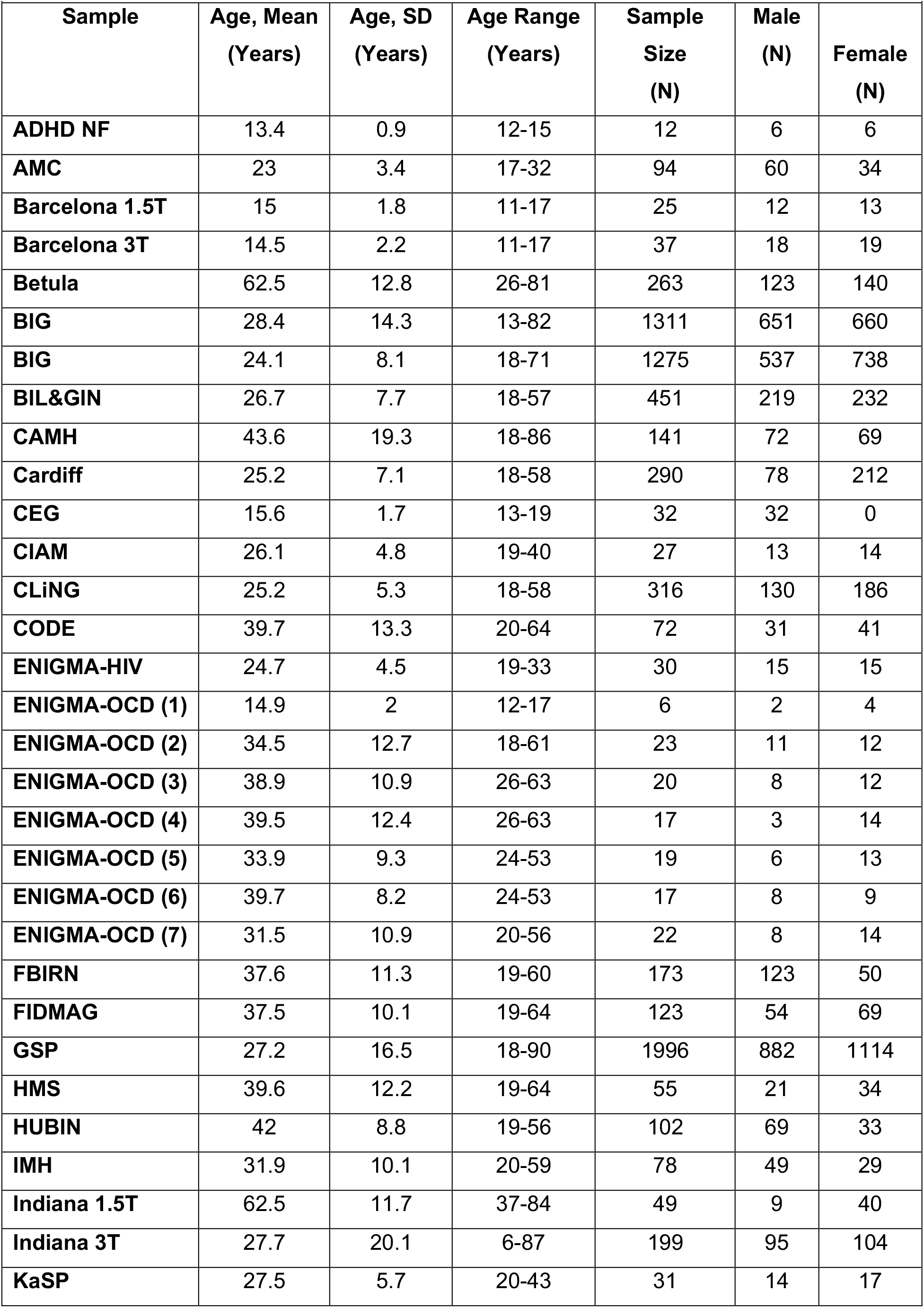
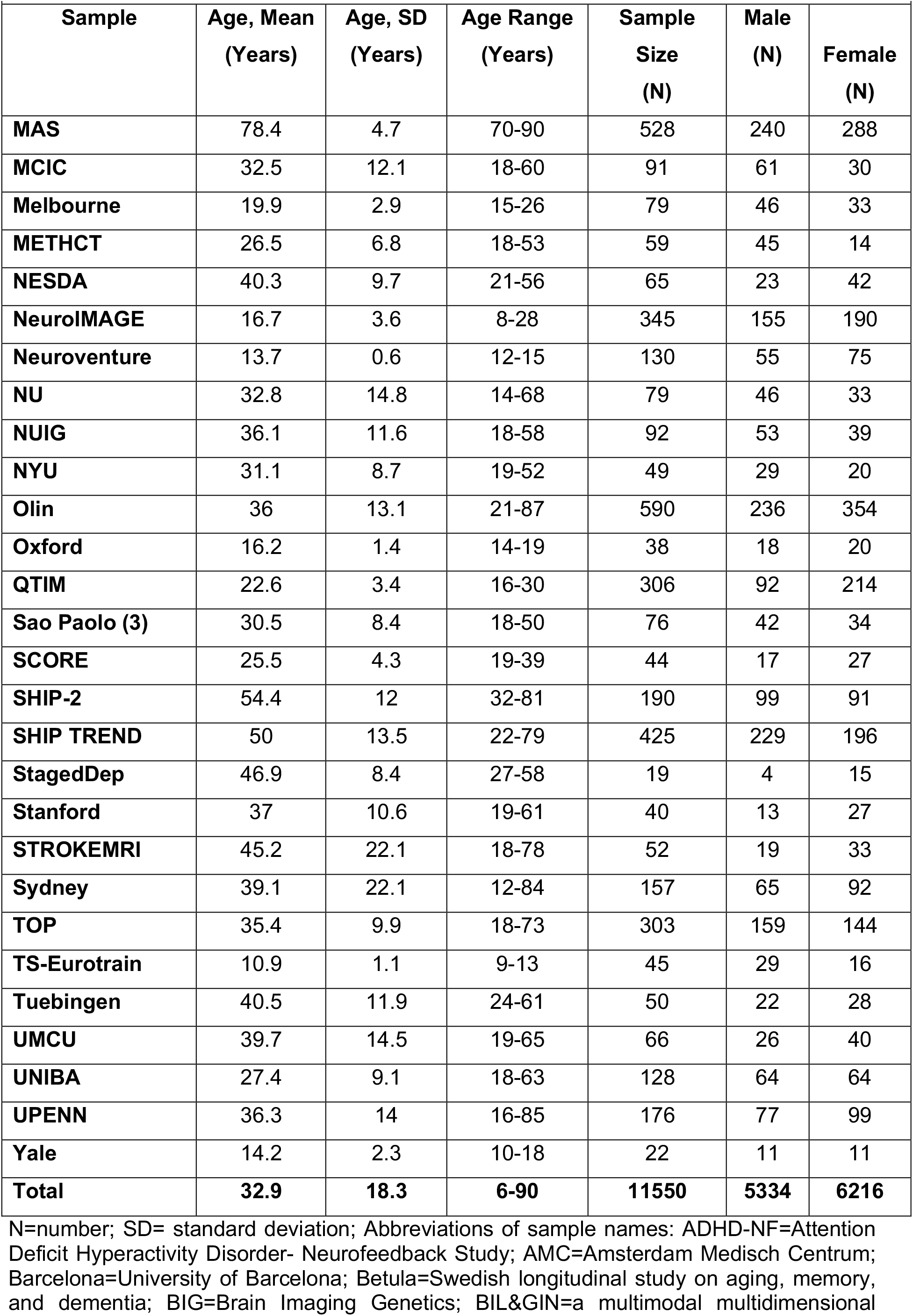
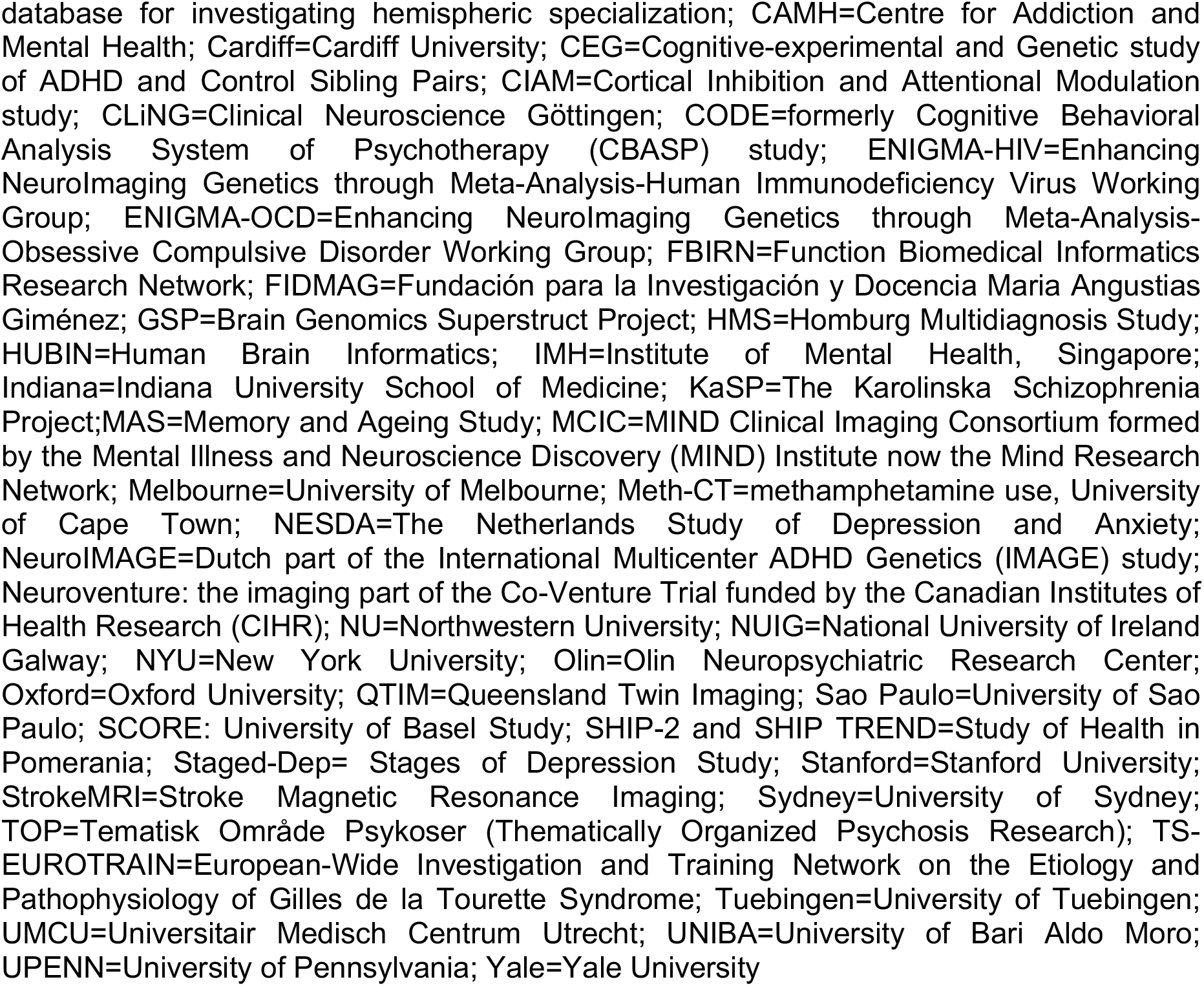
ENIGMA samples

### Neuroimaging

Detailed information on scanner vendor, magnet strength and acquisition parameters for each sample are presented in Table S1. For each sample, the intracranial volume (ICV) and the volume of the basal ganglia (caudate, putamen, pallidum, nucleus accumbens), thalamus, hippocampus, amygdala and lateral ventricles were extracted using FreeSurfer (http://surfer.nmr.mgh.harvard.edu) from high-resolution T1-weighted MRI brain scans (Fischl et al. 2002, 2012). Prior to data pooling, images were visually inspected at each site to exclude participants whose scans were improperly segmented. After merging the samples, outliers were identified and excluded using Mahalanobis distances. In each sample, the intracranial volume (Figure S1) was used to adjust the subcortical volumes via a formula based on the analysis of the covariance approach: ‘adjusted volume = raw volume – b x (ICV – mean ICV)’, where b is the slope of regression of a region of interest volume on ICV (Raz et al. 2005). The values of the subcortical volumes were then harmonized between sites using the ComBat method in R (Fortin, et al. 2017; 2018; Radua et al., 2019 this issue). Originally developed to adjust for batch effect in genetic studies, ComBat uses an empirical Bayes to adjust for inter-site variability in the data, while preserving variability related to the variables of interest.

### Fractional polynomial regression analyses

The effect of age on each ICV- and site-adjusted subcortical volume was modelled using high order fractional polynomial regression (Royston and Altman 1994; Sauerbrei et al. 2006) in each hemisphere. Because the effect of site (and thus scanner and Freesurfer version) were adjusted using ComBat, we only included sex as a covariate in the regression models. Fractional polynomial regression is currently considered the most advantageous modelling strategy for continuous variables (Moore et al. 2011) as it allows testing for a wider range of trajectory shapes than conventional lower-order polynomials (e.g., linear or quadratic) and for multiple turning points (Royston and Altman 1994; Royston et al. 1999). For each subcortical structure, the best model was obtained by comparing competing models of up to three power combinations. The powers used to identify the best fitting model were −2, −1, −0.5, 0.5, 1, 2, 3 and the natural logarithm (ln) function. The optimal model describing the association between age and each of the volumes was selected as the lowest degree model based on the partial F-test (if linear) or the likelihood-ratio test. To avoid overfitting at ages with more data points, we used the stricter 0.01 level of significance as the cut-off for each respective likelihood-ratio tests, rather than adding powers, until the 0.05 level was reached. For ease of interpretation we centred the volume of each structure so that the intercept of a fractional polynomial was represented as the effect at zero for sex. Fractional polynomial regression models were fitted using Stata/IC software v.13.1 (Stata Corp., College Station, TX). Standard errors were also adjusted for the effect of site in the FP regression.

We conducted two supplemental analyses: (a) we specified additional FP models separately for males and females and, (b) we calculated Pearson’s correlation coefficient between subcortical volumes and age in the early (6-29 years), middle (30-59 years), and late-life (60-90 years) age-group. The results of these analyses have been included in the supplemental material.

### Inter-individual variability

Inter-individual variability was assessed using two complimentary approaches. First, for each subcortical structure we compared the early (6-29 years), middle (30-59 years) and late-life (60-90 years) age-groups in terms of their mean inter-individual variability; these groups were defined following conventional notions regarding periods of development, midlife and aging. The variance of each structure in each age-group was calculated as

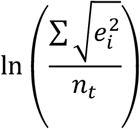

where e represents the residual variance of each individual (i) around the non-linear best fitting regression line, and n the number of observations in each age-group (t). The residuals (e_i_) were normally distributed suggesting good fit of the model without having over- or underfitted the data. Upon calculating the square root of the squared residuals we used the natural logarithm to account for the positive skewness of the new distribution. Then the mean inter-individual variability between early (6-29 years), middle (30-59 years) and late-life (60-90 years) age-groups was compared using between-groups omnibus tests for the residual variance around the identified best-fitting non-linear fractional polynomial model of each structure. The critical alpha value was set at 0.003 following Bonferroni correction for multiple comparisons.

The second approach entailed the quantification of the mean individual variability of each subcortical structure through a meta-analysis of the standard deviation of the adjusted volumes according to the method proposed by Senior et al. 2016.

### Centile Curves

Reference curves for each structure by sex and hemisphere were produced from ICV- and site-adjusted volumes as normalized growth centiles using the parametric Lambda (λ), Mu (μ), Sigma (σ) (LMS) method (Cole and Green, 1992) implemented using the Generalised Additive Models for Location, Scale and Shape (GAMLSS) in R (http://cran.r-project.org/web/packages/gamlss/index.html)(Rigby and Stasinopoulos, 2005; 2007). LMS allows for the estimation of the distribution at each covariate value after a suitable transformation and is summarized using three smoothing parameters, the Box-Cox power λ, the mean μ and the coefficient of variation σ. GAMLSS uses an iterative maximum (penalized) likelihood estimation method to estimate λ, μ and σ as well as distribution dependent smoothing parameters and provides optimal values for effective degrees of freedom (edf) for every parameter (Indrayan, 2014). This procedure minimizes the Generalized Akaike Information Criterion (GAIC) goodness of fit index; smaller GAIC values indicate better fit of the model to the data. GAMLSS is a flexible way to derive normalized centile curves as it allows each curve to have its own number of edf while overcoming biased estimates resulting from skewed data.

## Results

### Fractional polynomial regression analyses

The volume of the caudate, putamen, globus pallidus and nucleus accumbens peaked early during the first decade of life and showed a linear decline immediately thereafter (Figure 2, Figures S2-S4). The age-related trajectories of the thalamic, hippocampal and amygdala volumes followed a flattened, inverted U-curve (Figure 3, Figures S5-S6). Specifically, the volumes of these structures were largest during the first 2-3 decades of life, remained largely stable until the 6^th^ decade and declined gradually thereafter (Table S2). The volume of the lateral ventricles bilaterally increased steadily with age (Figure S7). The smallest proportion of variance explained by age and its FP derivatives was noted in the right amygdala (7%) and the largest in the lateral ventricles bilaterally (38%) (Table S2).

**Figure 2.**
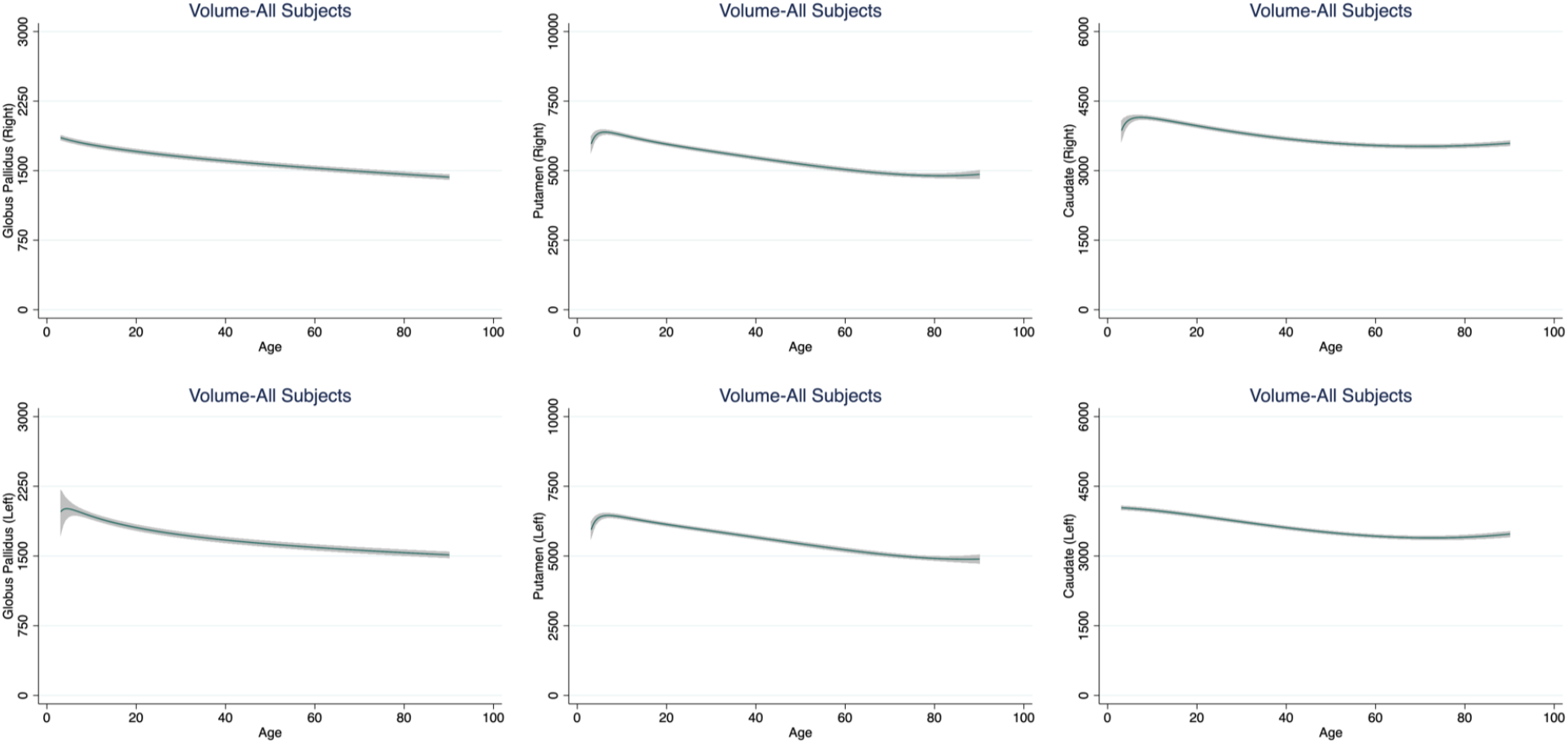
Fractional Polynomial Plots for the Volume of the Basal Ganglia. Fractional Polynomial plots of adjusted volumes (mm^3^) against age (years) with a fitted regression line (solid line) and 95% confidence intervals (shaded area).

**Figure 3.**
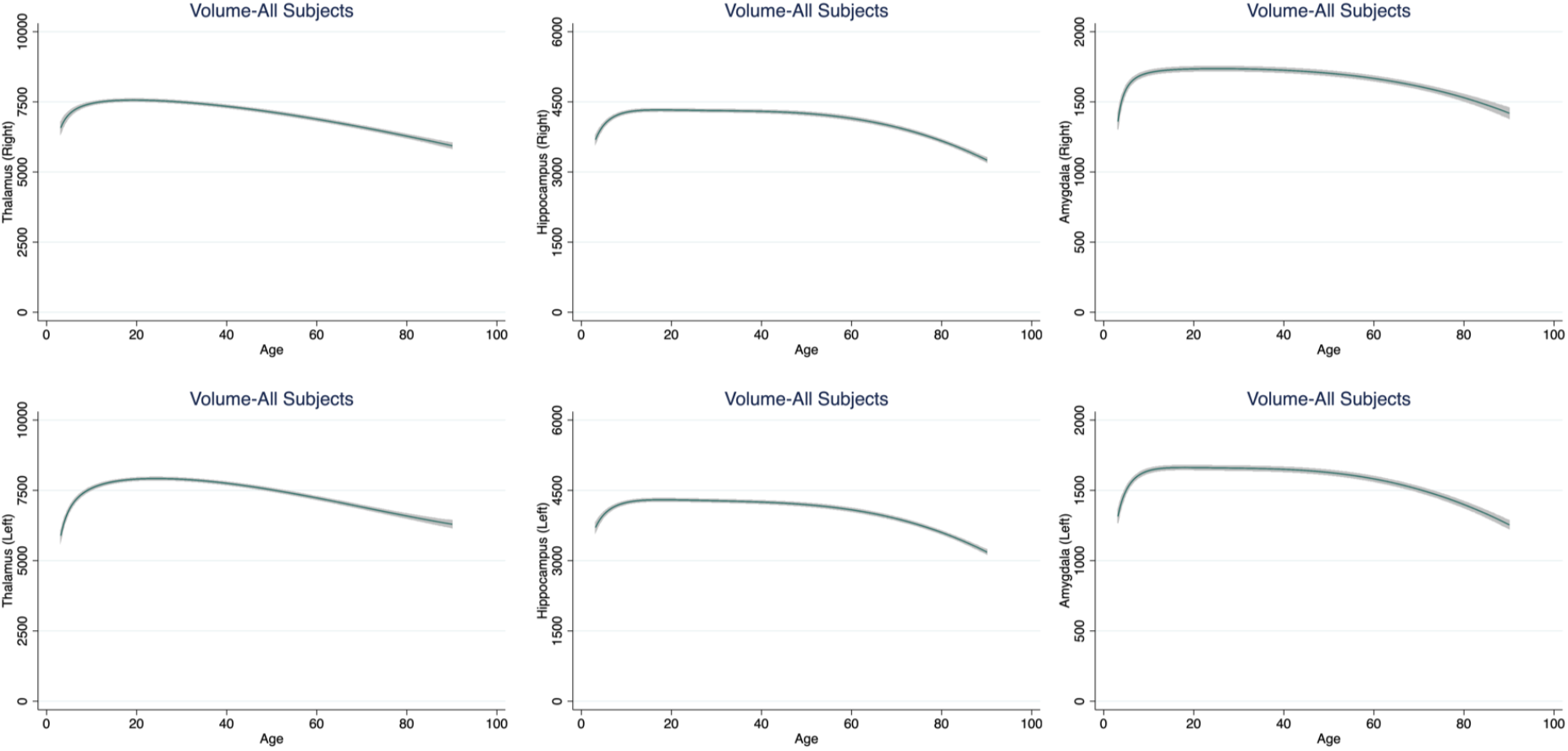
Fractional Polynomial Plots for the Volume of the Thalamus, Hippocampus and Amygdala. Fractional Polynomial plots of adjusted volumes (mm^3^) against age (years) with a fitted regression line (solid line) and 95% confidence intervals (shaded area).

Striatal volumes correlated negatively with age throughout the lifespan with the largest coefficients observed in the middle-life age-group (r=-0.39 to −0.20) and the lowest (|r|<0.05) in the late-life age-group, particularly in the caudate. The volumes of the thalamus, the hippocampus and the amygdala showed small positive correlations with age (r≈0.16) in the early-life age-group. In the middle-life age-group, the correlation between age and subcortical volumes became more negative (r=-0.30 to −0.27) for the thalamus but remained largely unchanged for the amygdala and the hippocampus. In the late-life age-group, the largest negative correlation coefficients between age and volume were observed for the hippocampus bilaterally (r=-0.44 to −0.39). The correlation between age and lateral ventricular volumes bilaterally increased throughout the lifespan from r=0.19 to 0.20 in early-life age-group to r= 0.40 to 0.45 in the late-life age-group (Table S3). No effect of sex was noted for any pattern of correlation between subcortical volumes and age in any age-group.

#### Inter-individual variability

For each structure, the mean inter-individual variability in volume in each age-group is shown in Table S5. Inter-individual variance was significantly higher for the hippocampus, thalamus amygdala and lateral ventricles bilaterally in the late-life age-group compared to both the early- and middle-life group. These findings were recapitulated when data were analysed using a meta-analytic approach (Figure 4 and Figure S8).

**Figure 4.**
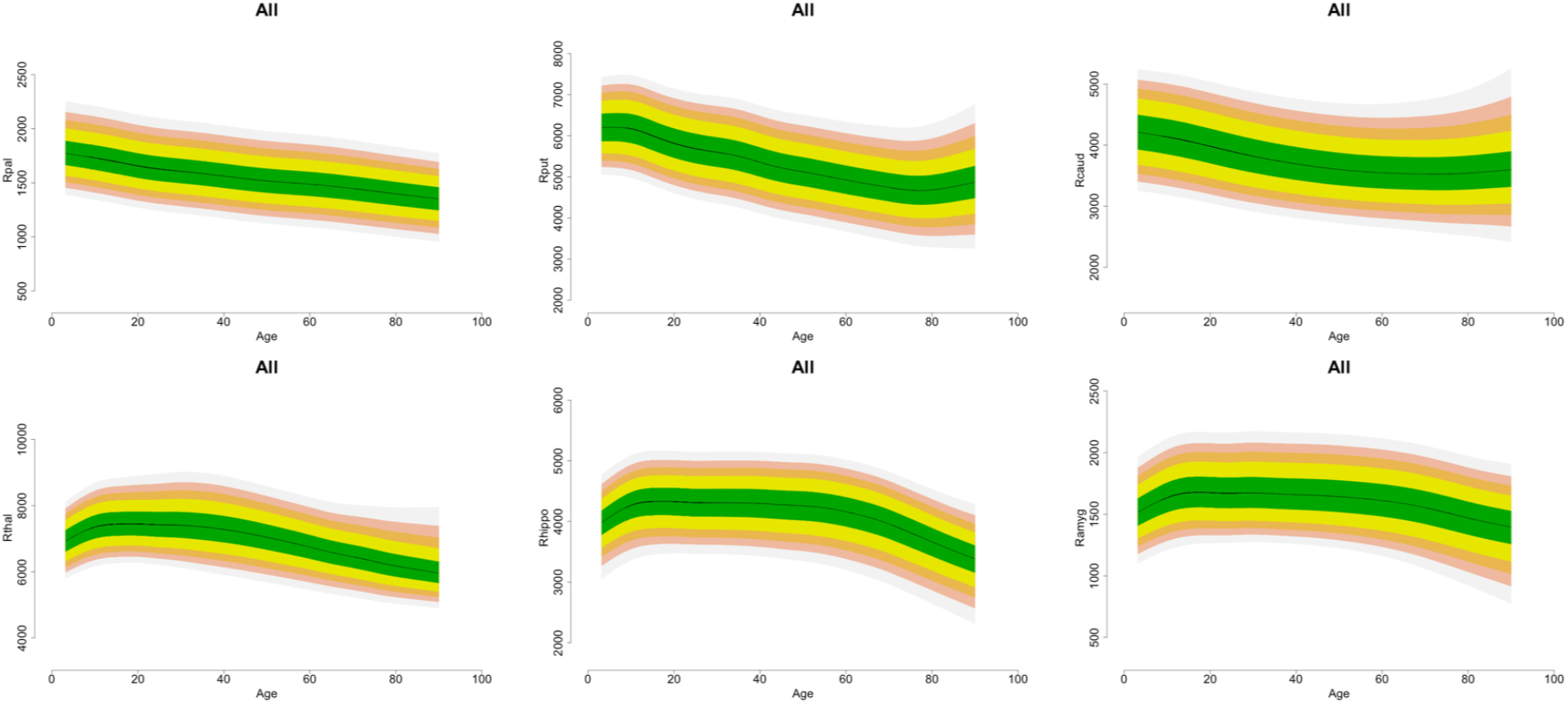
Mean Inter-individual Variability of Subcortical Volumes. Mean individual variability for each subcortical structure was estimated by means of a meta-analysis of the standard deviation of the adjusted volumes in each age-group.

#### Normative Centile Curves

Centile normative values for each subcortical structure stratified by sex and hemisphere are shown in Tables S6-S8.

## Discussion

We analysed subcortical volumes from 18,605 healthy individuals from multiple crosssectional cohorts to infer age-related trajectories between the ages of 3 to 90 years. Our lifespan perspective and our large sample size complement and enrich previous literature on age-related changes in subcortical volumes.

We found three distinct age-related trajectories. The volume of the lateral ventricles increased monotonically with age. Striatal and pallidal volumes peaked in childhood and declined thereafter. The volumes of the thalamus, hippocampuus and amygdala peaked later and showed a prolonged period of stability lasting till the 6^th^ decade of life, before they also started to decline. The trajectories defined here represent a close approximation to a normative reference dataset and are in line with findings from Pomponio et al (2019) who also used harmonised multi-site MRI data from 10,323 individuals aged 3-96 years. Similar findings were reported by Douaud and colleagues (2014) who analysed volumetric data from 484 healthy participants aged 8 to 85 years; they also noted the similarity in the age-related trajectories of the thalamus, hippocampus and the amygdala. Our results also underscore the acceleration in age-related decline from the 6^th^ decade of life onwards. This effect seemed relatively more pronounced for the hippocampus, compared to the other subcortical regions, as observed in other studies (Jernigan et al. 2001; Pomponio et al. 2019; Raz et al. 2010).

The trajectories of subcortical volumes are shaped by genetic and non-genetic exposures, biological or otherwise (Eyler et al. 2011; Somel et al. 2010; Wardlaw et al. 2011). Our findings of high age-related inter-individual variability in the volumes of the thalamus, hippocampus and amygdala suggest that these structures may be more susceptible to person-specific exposures, or late-acting genes, particularly from the 6th decade onwards.

In medicine, biological measures from each individual are typically categorised as normal or otherwise in reference to a population derived normative range. This approach is yet to be applied to neuroimaging data, despite the widespread use of structural MRI for clinical purposes and the obvious benefit of a reference range from the early identification of deviance (Dickie et al. 2013; Pomponio et al. 2019). Alzheimer’s disease provides an informative example as the degree of baseline reduction in medial temporal regions, and particularly the hippocampus, is one of the most significant predictors of conversion from mild cognitive impairment to Alzheimer’s disease (Risacher et al. 2009). The data presented here demonstrate the power of international collaborations within ENIGMA for analyzing very large-scale datasets that could eventually lead to normative range for brain volumes for well-defined reference populations. The unique strengths of this study are the availability of age-overlapping cross-sectional data from healthy individuals, lifespan coverage and the use of standardized protocols for volumetric data extraction across all samples. Study participants in each site were screened to ensure mental and physical wellbeing at the time of scanning using procedures considered as standard in designating study participants as healthy controls. Although health is not a permanent attribute, it is extremely unlikely given the size of our sample that our results could have been systematically biased by incipient disease.

A similar longitudinal design would be near infeasible in terms of recruitment and retention both of participants and investigators. Although multisite studies have to account for differences in scanner type and acquisition, lengthy longitudinal designs encounter similar issues due to inevitable changes in scanner type and strength and acquisition parameters over time. In this study, the use of age-overlapping samples from multiple different countries has the theoretical advantage of diminishing systematic biases reflecting cohort and period effects (Glenn, 2003; Keyes et al. 2010) that are likely to operate in single site studies.

In conclusion, we used existing data to derive age-related trajectories of regional subcortical volumes. The size and age-coverage of the analysis sample has the potential to disambiguate uncertainties regarding developmental and aging changes in subcortical volumes while the normative centile values could be further developed to derive clinically meaningful predictors of risk of adverse health outcomes.

## Acknowledgments

This study presents independent research funded by multiple agencies. The funding sources had no role in the study design, data collection, analysis, and interpretation of the data. The views expressed in the manuscript are those of the authors and do not necessarily represent those of any of the funding agencies. Dr. Dima received funding from the National Institute for Health Research (NIHR) Biomedical Research Centre at South London and Maudsley NHS Foundation Trust and King’s College London, the Psychiatry Research Trust and 2014 NARSAD Young Investigator Award. Dr. Frangou received support from the National Institutes of Health (R01 MH104284) the European Community’s Seventh Framework Programme (FP7/2007-2013) (grant agreement n°602450). FBIRN data collection and analysis was supported by the National Center for Research Resources at the National Institutes of Health (grant numbers: NIH 1 U24 RR021992 (Function Biomedical Informatics Research Network) and NIH 1 U24 RR025736-01 (Biomedical Informatics Research Network Coordinating Center; http://www.birncommunity.org). FBIRN data was processed by the UCI High Performance Computing cluster supported by the National Center for Research Resources and the National Center for Advancing Translational Sciences, National Institutes of Health, through Grant UL1 TR000153. Betula sample: Data collection was supported by a grant from Knut and Alice Wallenberg Foundation (KAW). Indiana sample: Brenna McDonald acknowledges the support in part by grants to BCM from Siemens Medical Solutions, from the members of the Partnership for Pediatric Epilepsy Research, which includes the American Epilepsy Society, the Epilepsy Foundation, the Epilepsy Therapy Project, Fight Against Childhood Epilepsy and Seizures (F.A.C.E.S.), and Parents Against Childhood Epilepsy (P.A.C.E.), from the Indiana State Department of Health Spinal Cord and Brain Injury Fund Research Grant Program, and by a Project Development Team within the ICTSI NIH/NCRR Grant Number RR025761. Andrew Saykin received support from U.S. National Institutes of Health grants R01 AG19771, P30 AG10133 and R01 CA101318. For the QTIM sample: We are grateful to the twins for their generosity of time and willingness to participate in our study. We also thank the many research assistants, radiographers, and other staff at QIMR Berghofer Medical Research Institute and the Centre for Advanced Imaging, University of Queensland. QTIM was funded by the Australian National Health and Medical Research Council (Project Grants No. 496682 and 1009064) and US National Institute of Child Health and Human Development (RO1HD050735). Lachlan Strike was supported by a University of Queensland PhD scholarship. The TOP study was supported by the European Community’s Seventh Framework Programme (FP7/2007-2013), grant agreement n°602450. The Southern and Eastern Norway Regional Health Authority supported Lars T. Westlye (grant no. 2014-097) and STROKEMRI (grant no. 2013-054). For the HUBIN sample: HUBIN was supported by the Swedish Research Council (K2007-62X-15077-04-1, K2008-62P-20597-01-3. K2010-62X-15078-07-2, K2012-61X-15078-09-3), the regional agreement on medical training and clinical research between Stockholm County Council, and the Karolinska Institutet, and the Knut and Alice Wallenberg Foundation. P.M.T., N.J., M.J.W., S.E.M., O.A.A., D.A.R., L.S., D.J.V., T.G.M. v.E., D.G., and D.P.H. were supported in part by a Consortium grant (U54 EB020403 to P.M.T.) from the NIH Institutes contributing to the Big Data to Knowledge (BD2K) Initiative.

## Data Availability Statement

The ENIGMA Lifespan Working Group welcomes expression of interest from researchers in the field who wish to use the ENIGMA samples. Data sharing is possible subsequent to consent for the principal investigators of the contributing datasets. Requests should be directed to the corresponding authors.

