## Supplementary Figures and Tables for "Subcortical Volume Trajectories across the Lifespan: Data from 18,605 healthy individuals aged 3-90 years"

**Online Supplement**

**Supplementary Figures**

**Figure S1.** Intracranial volume (ICV) by sex and age (mean, standard error)

**Figure S2.** Age-related trajectories in globus pallidus, putamen, and caudate in males

**Figure S3.** Age-related trajectories in globus pallidus, putamen, and caudate in females

**Figure S4.** Age-related trajectories in nucleus accumbens

**Figure S5.** Age-related trajectories in thalamus, hippocampus, and amygdala in males

**Figure S6.** Age-related trajectories in thalamus, hippocampus, and amygdala in females

**Figure S7.** Age-related trajectories in lateral ventricles

**Figure S8.** Meta-analysis of the Pooled Standard Deviation of the Volume of each Subcortical Structure Stratified by Sex

**Supplementary Tables**

**Table S1.** Screening Process and Eligibility Criteria, Scanner, Image Acquisition Parameters and Image Segmentation Software

**Table S2.** Age at Maximum Fitted Value for Subcortical Volumes

**Table S3.** Variance Explained by Age in Fractional Polynomial Model

**Table S4:** Pearson's Correlation Coefficient between Age and Subcortical Volumes

**Table S5.** Inter-individual Variations in Subcortical Volumes

**Table S6.** Centile Values for Subcortical Volumes- All participants

**Table S7.** Centile Values for Subcortical Volumes in males

**Table S8.** Centile Values for Subcortical Volumes in females

**Figure S1. Intracranial volume (ICV) by Sex and Age (mean, standard error)**

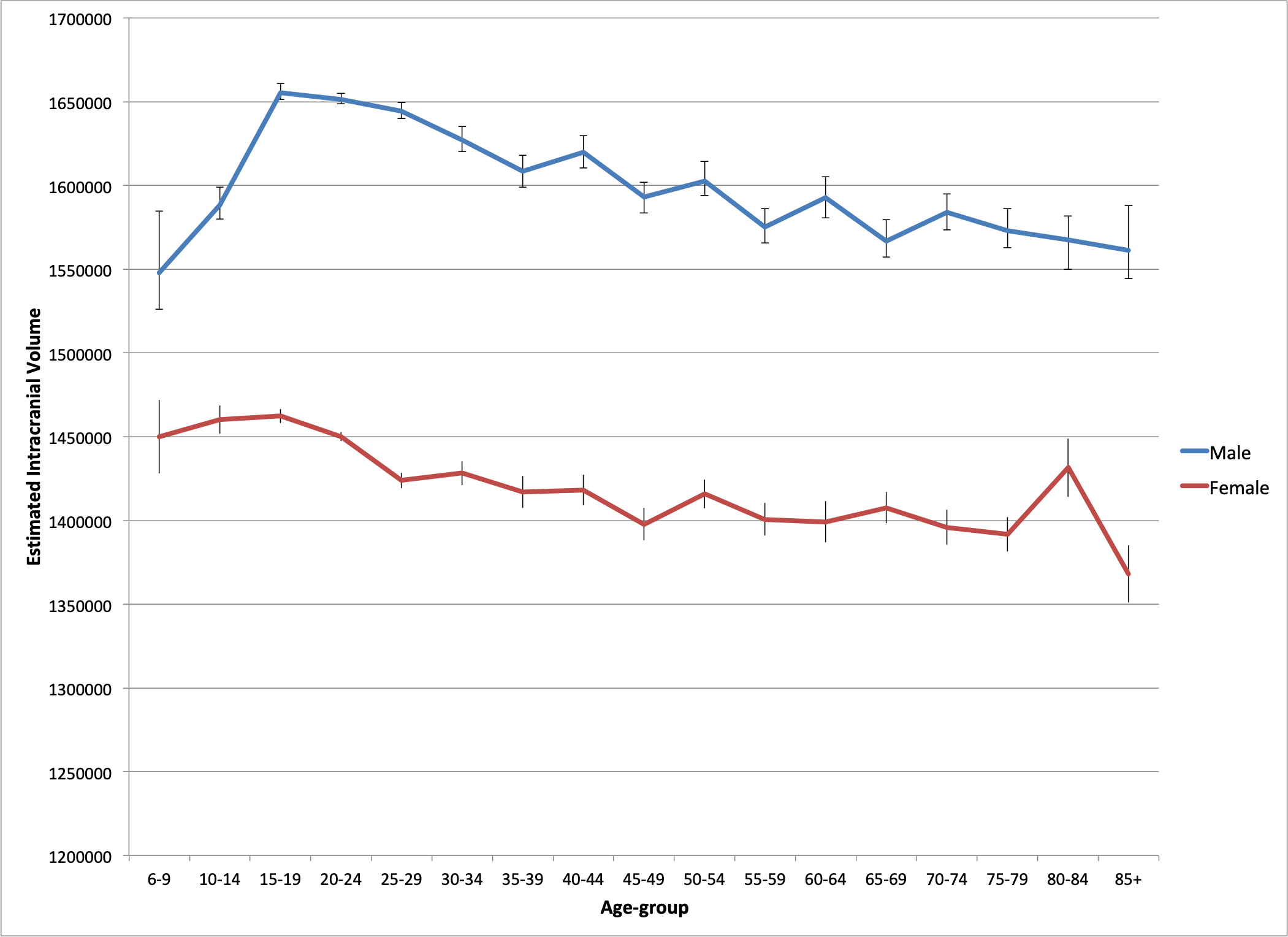

**Figure S2. Age-related trajectories in Globus Pallidus, Putamen and Caudate in Males**

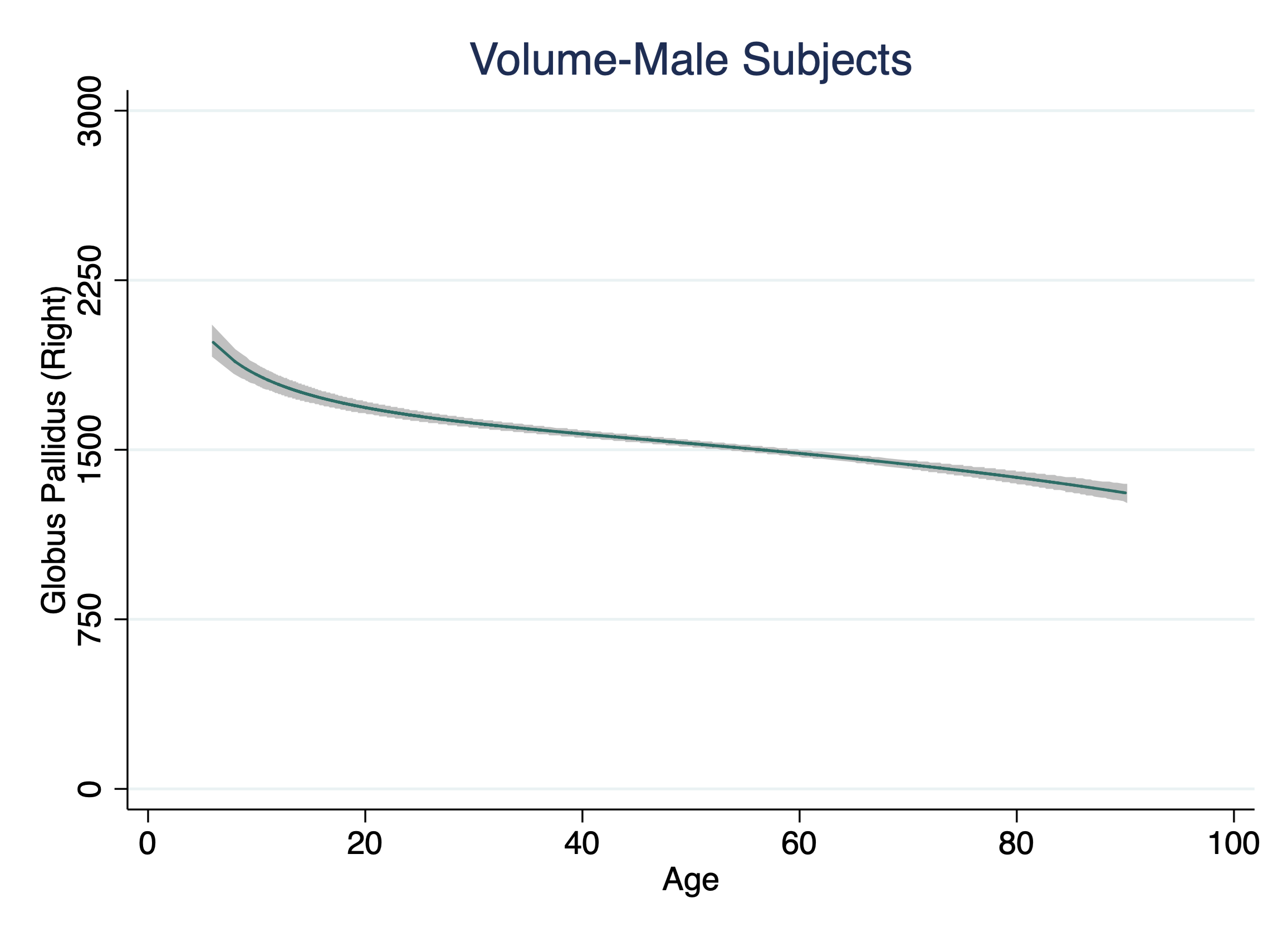

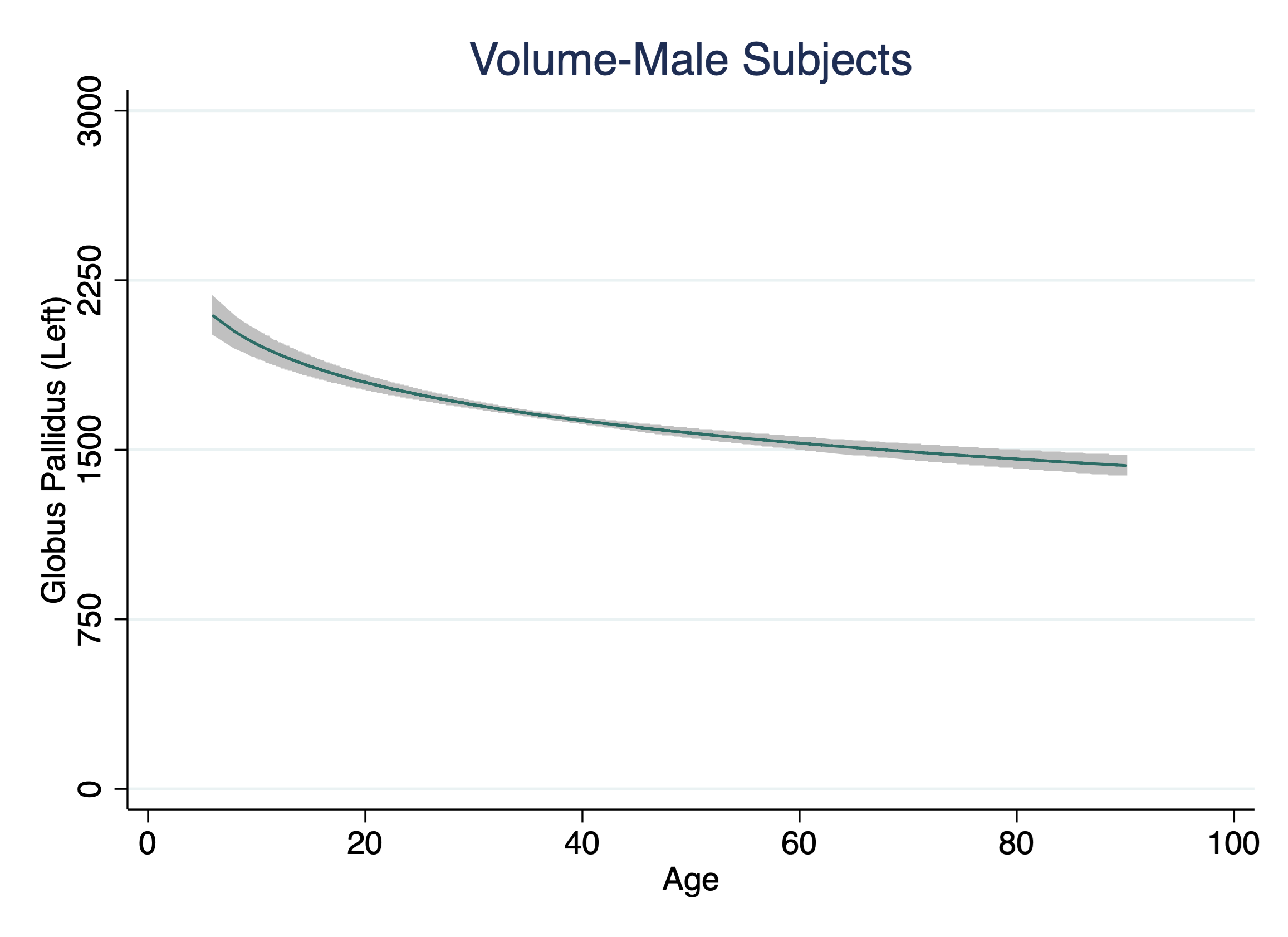

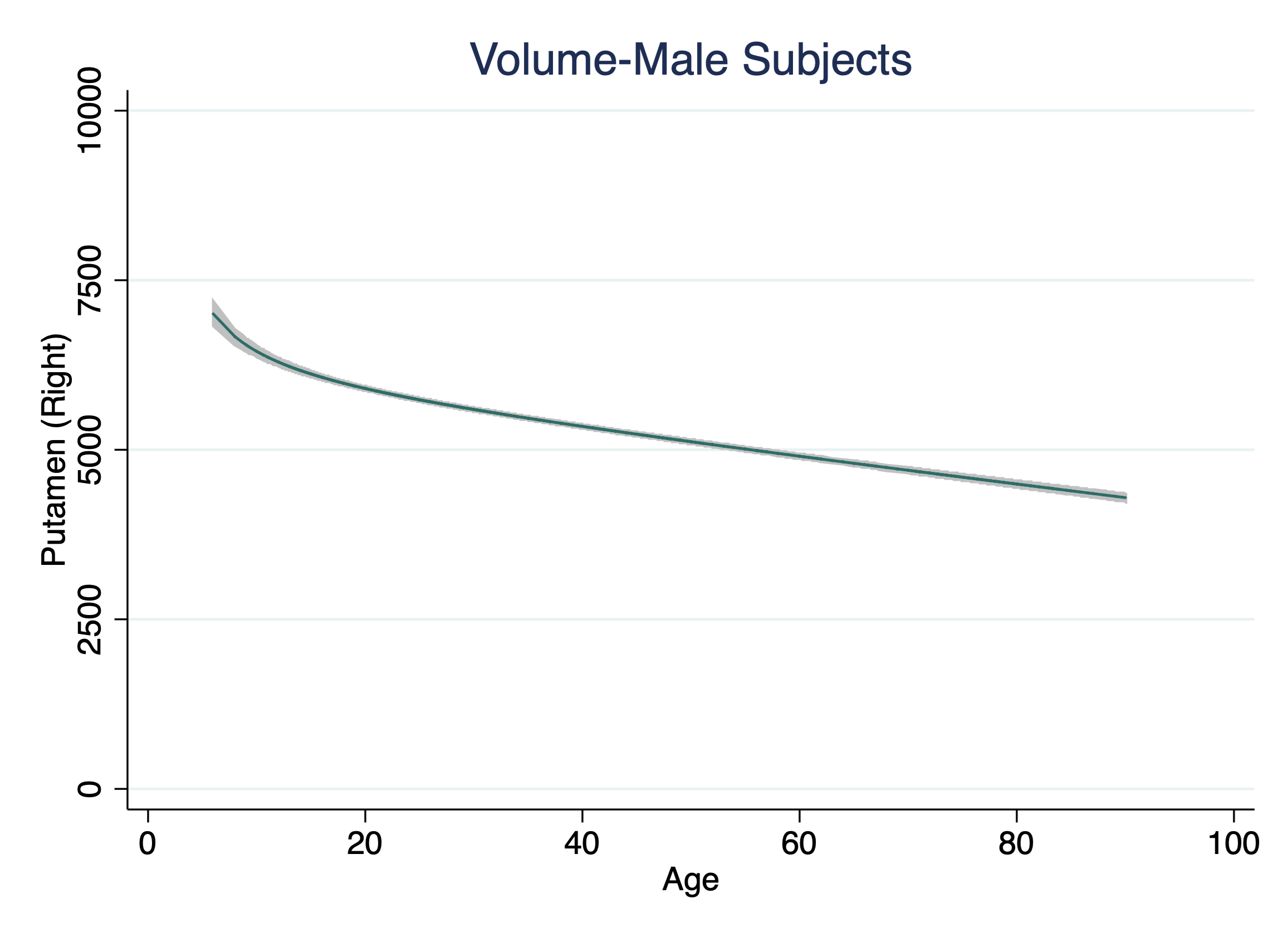

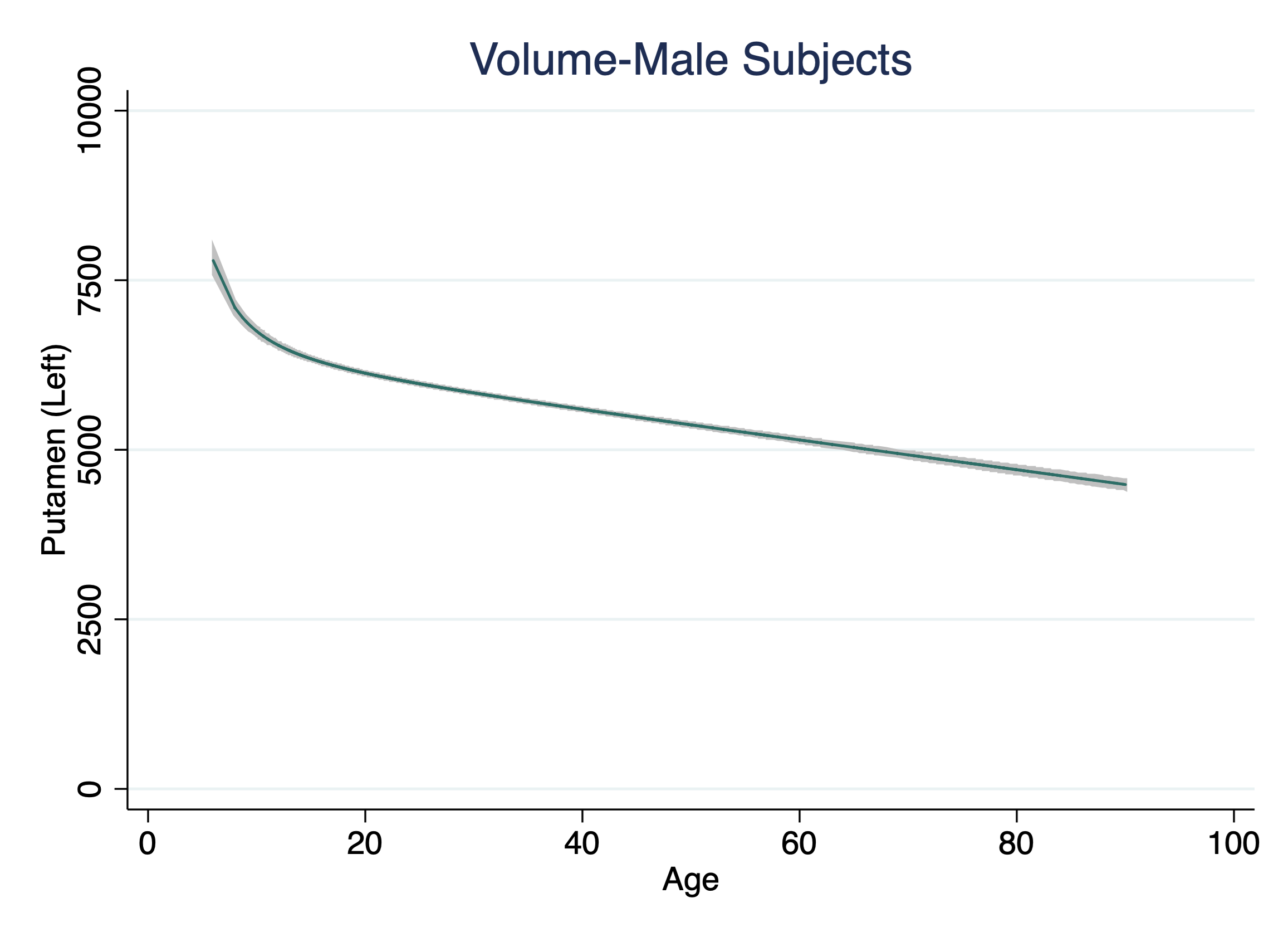

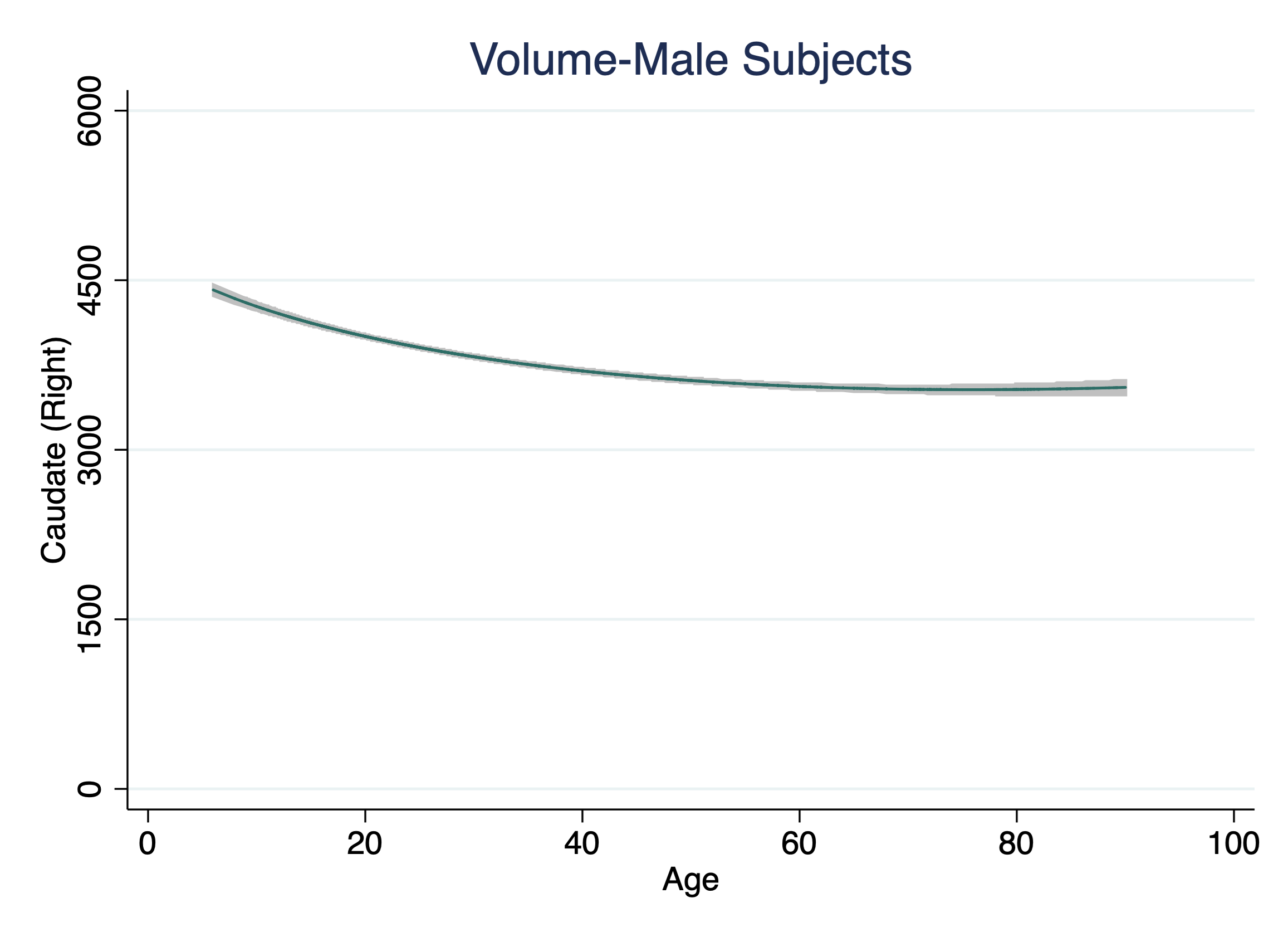

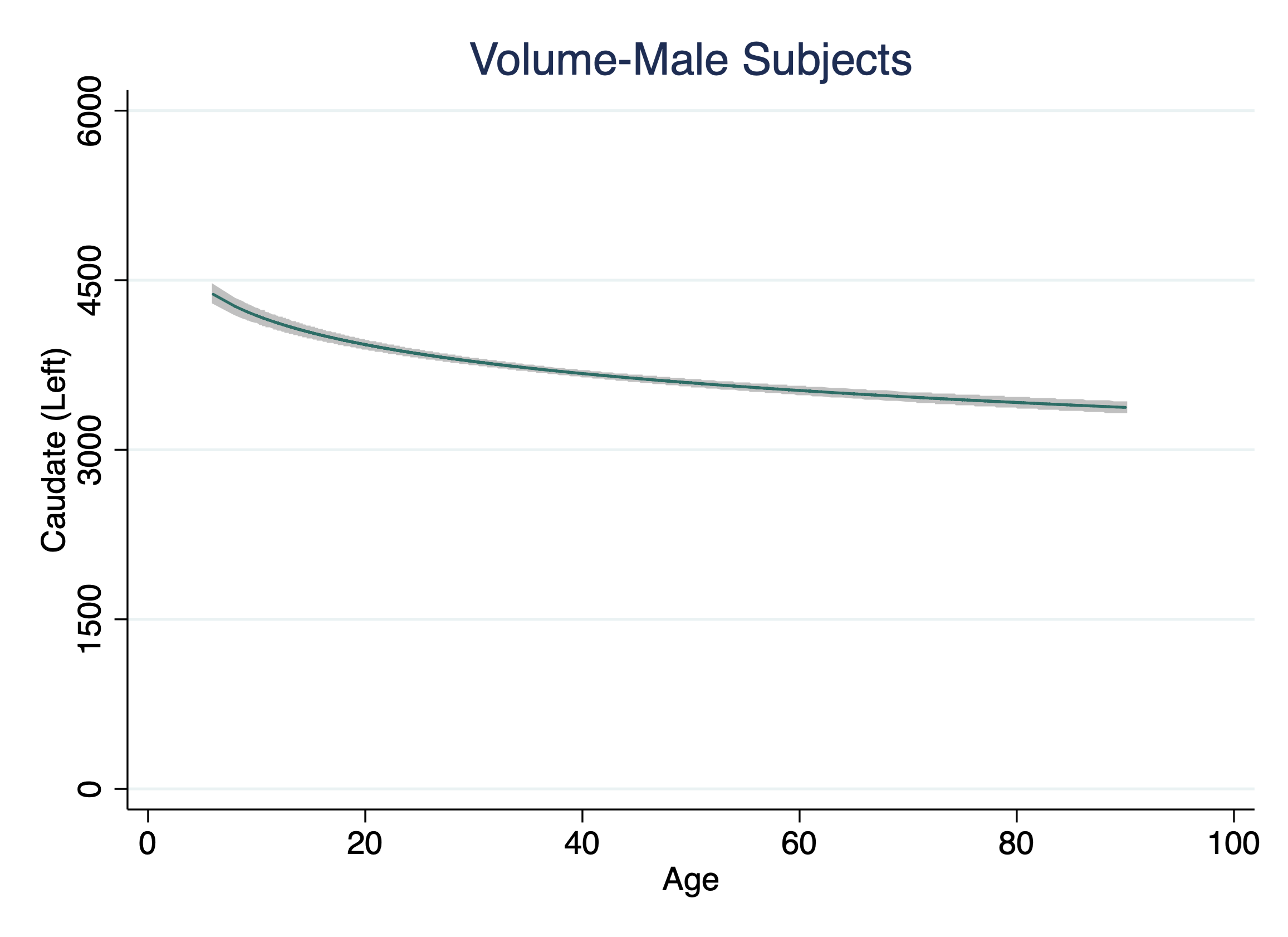

**Figure S3. Age-related trajectories in Globus Pallidus, Putamen and Caudate in Females**

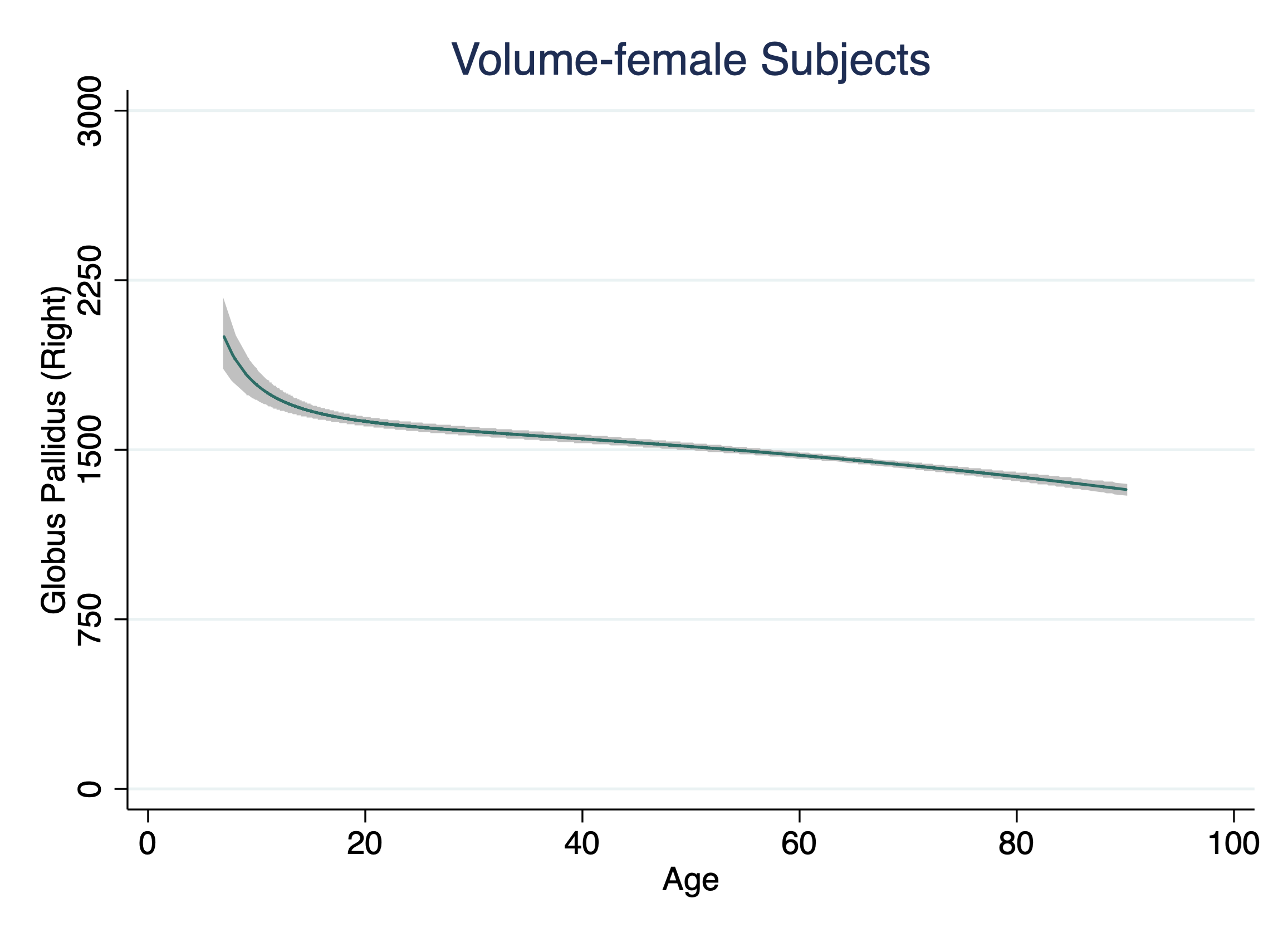

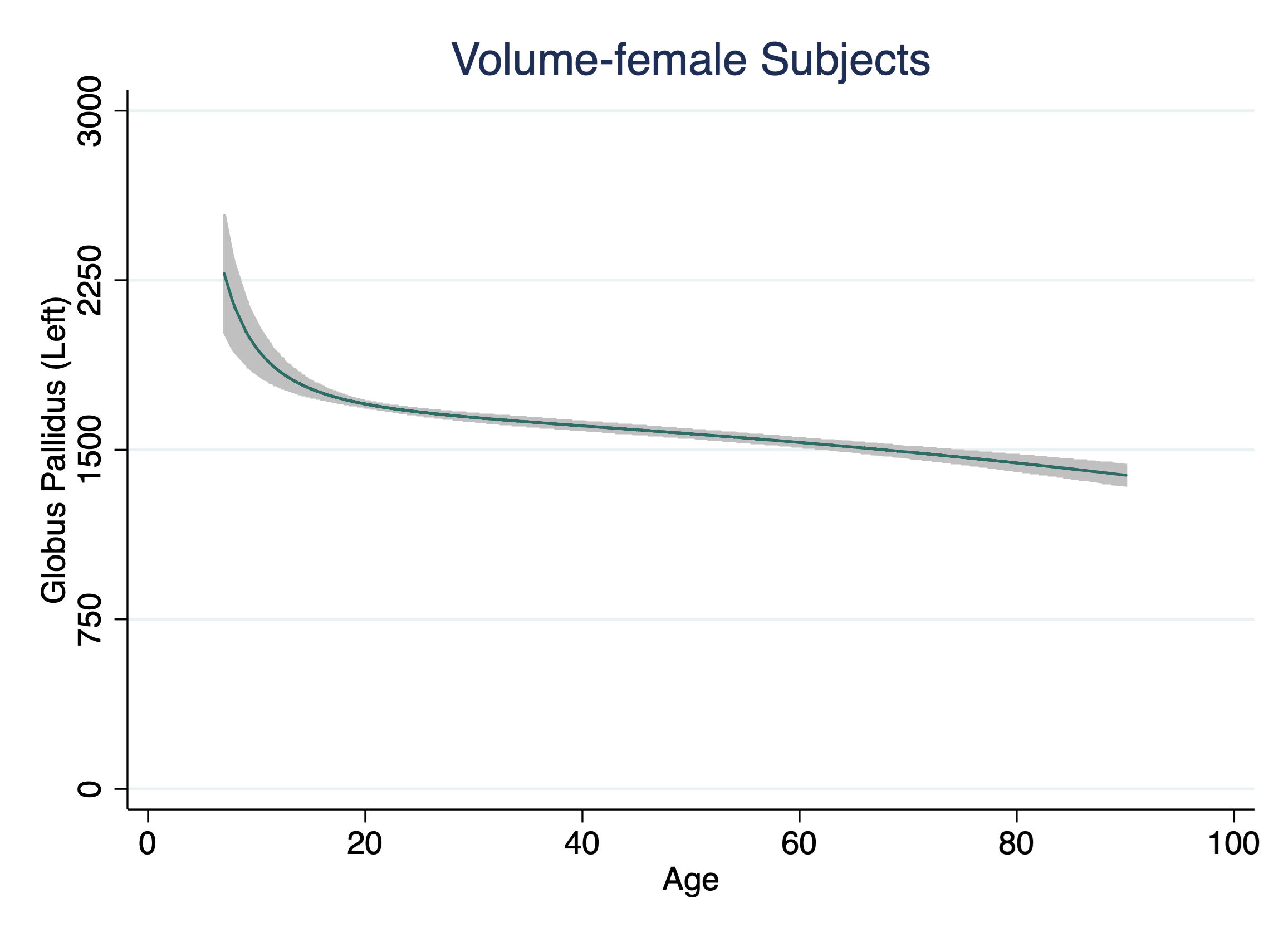

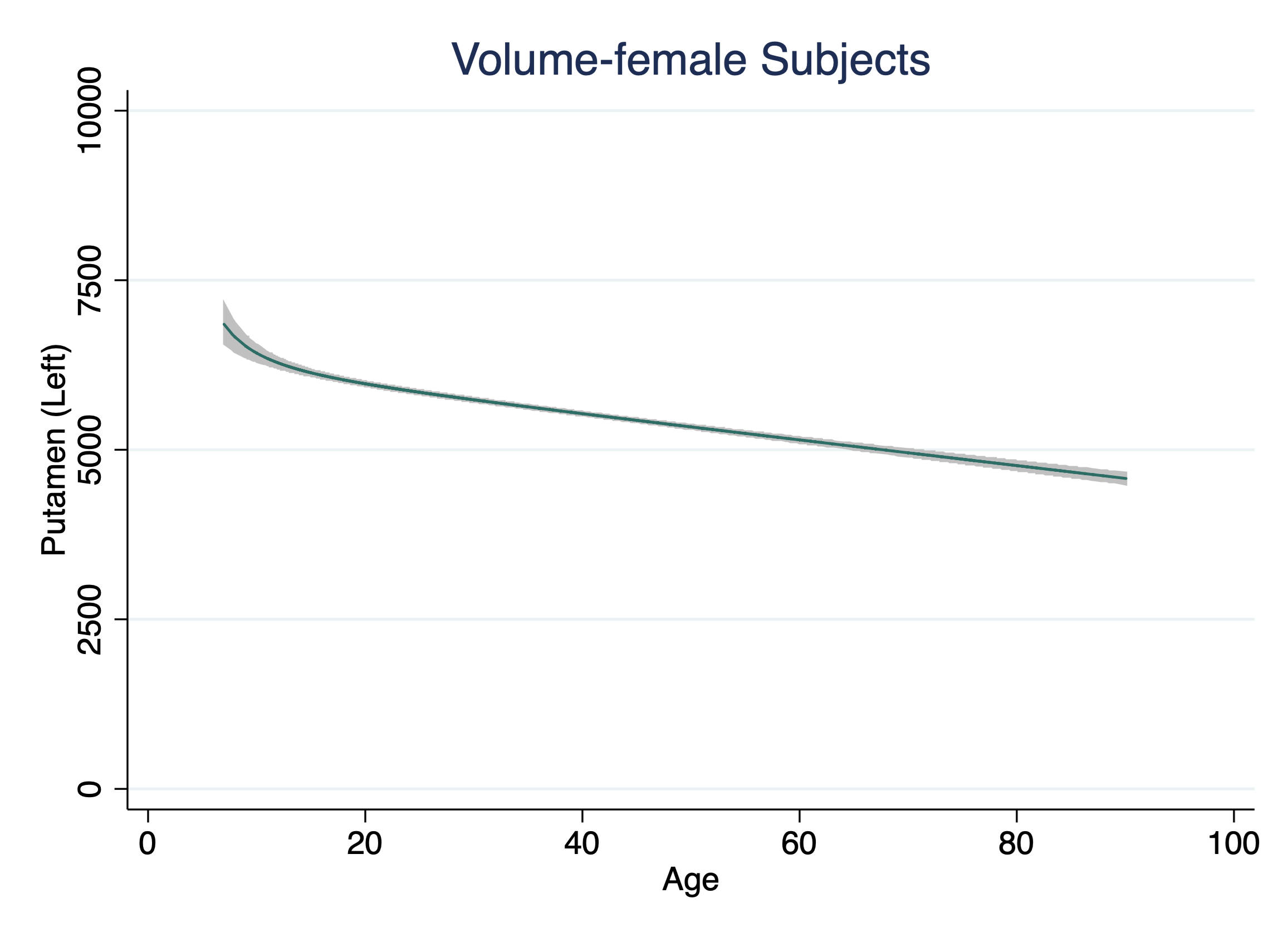

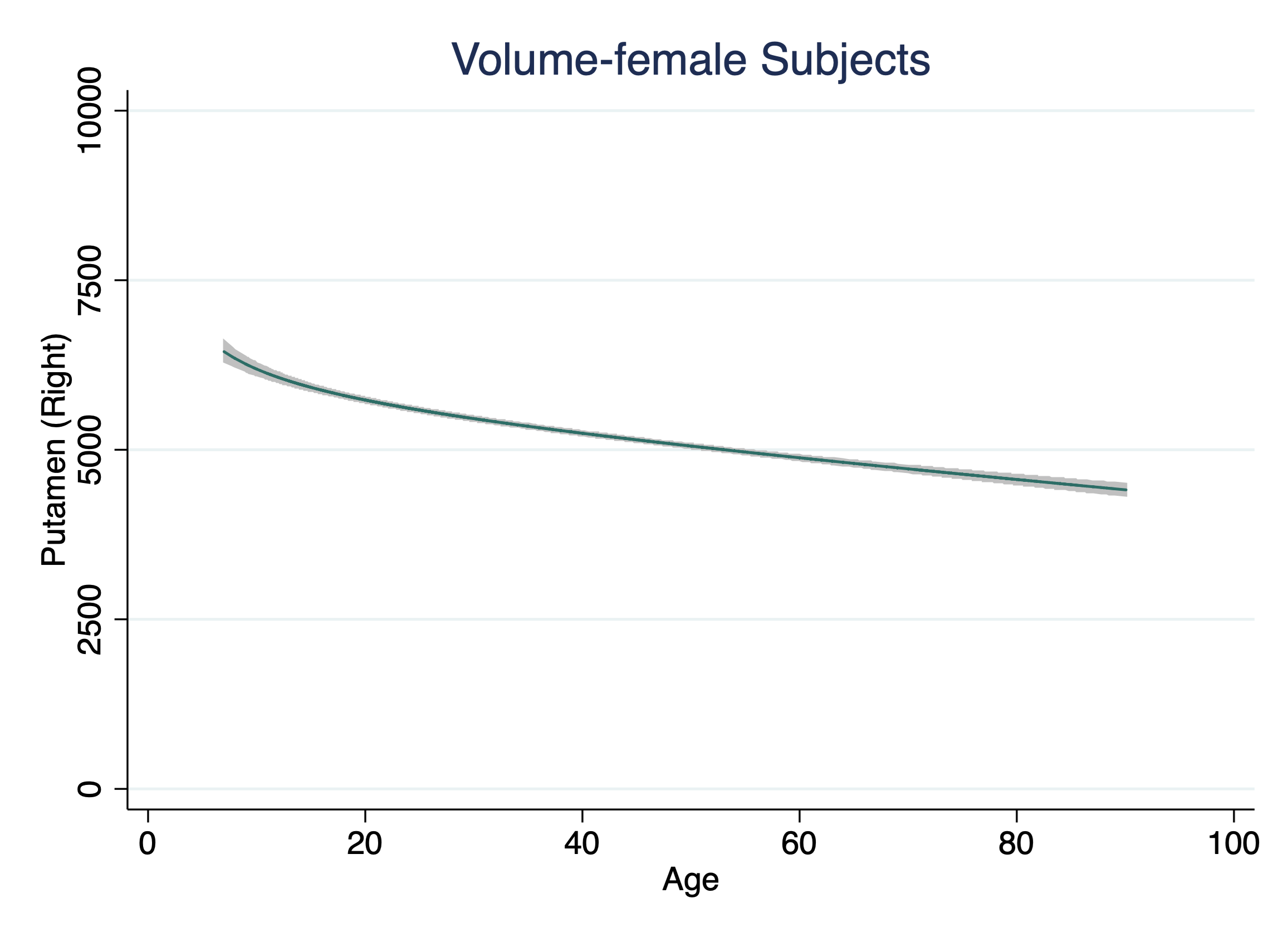

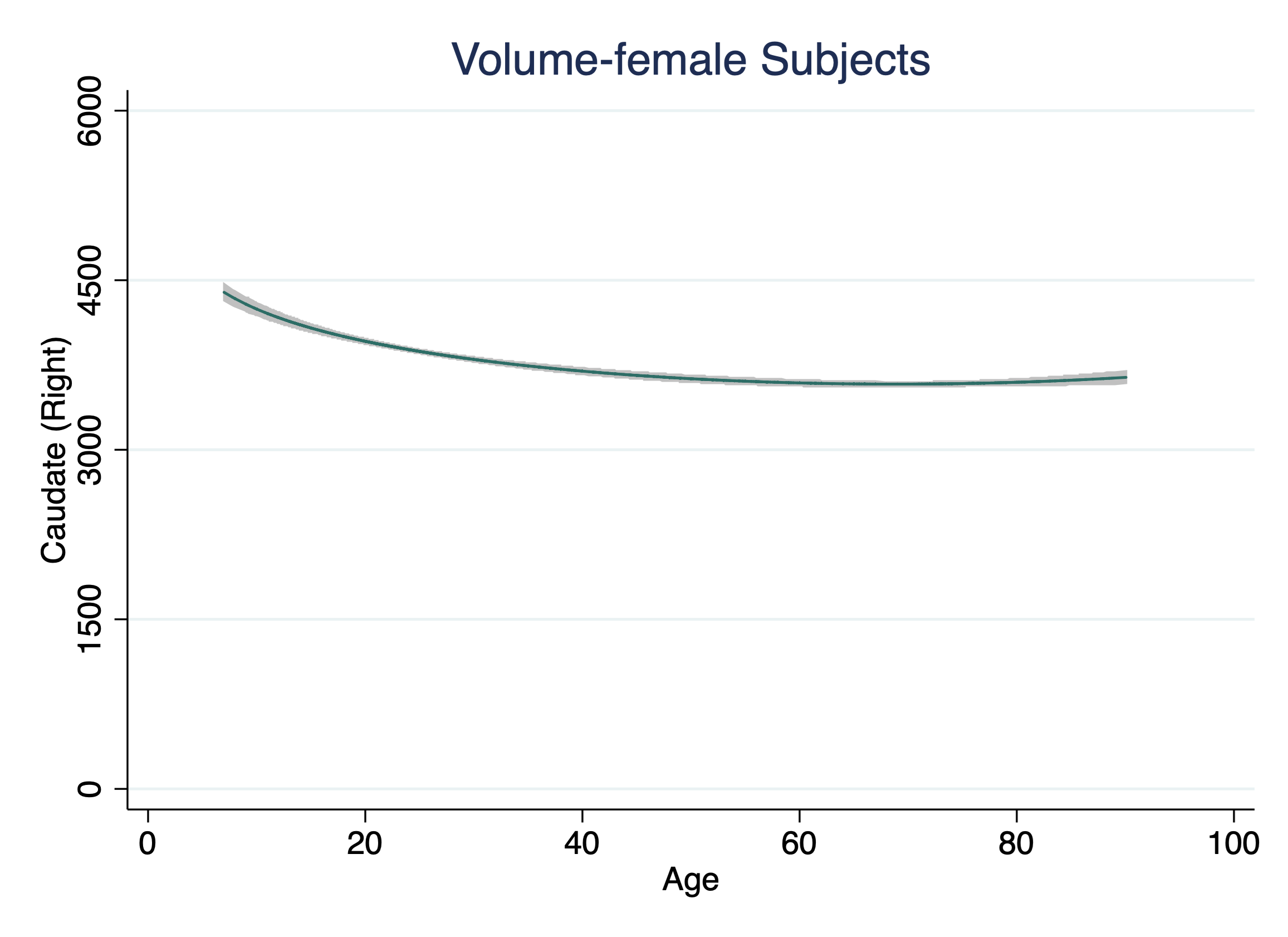

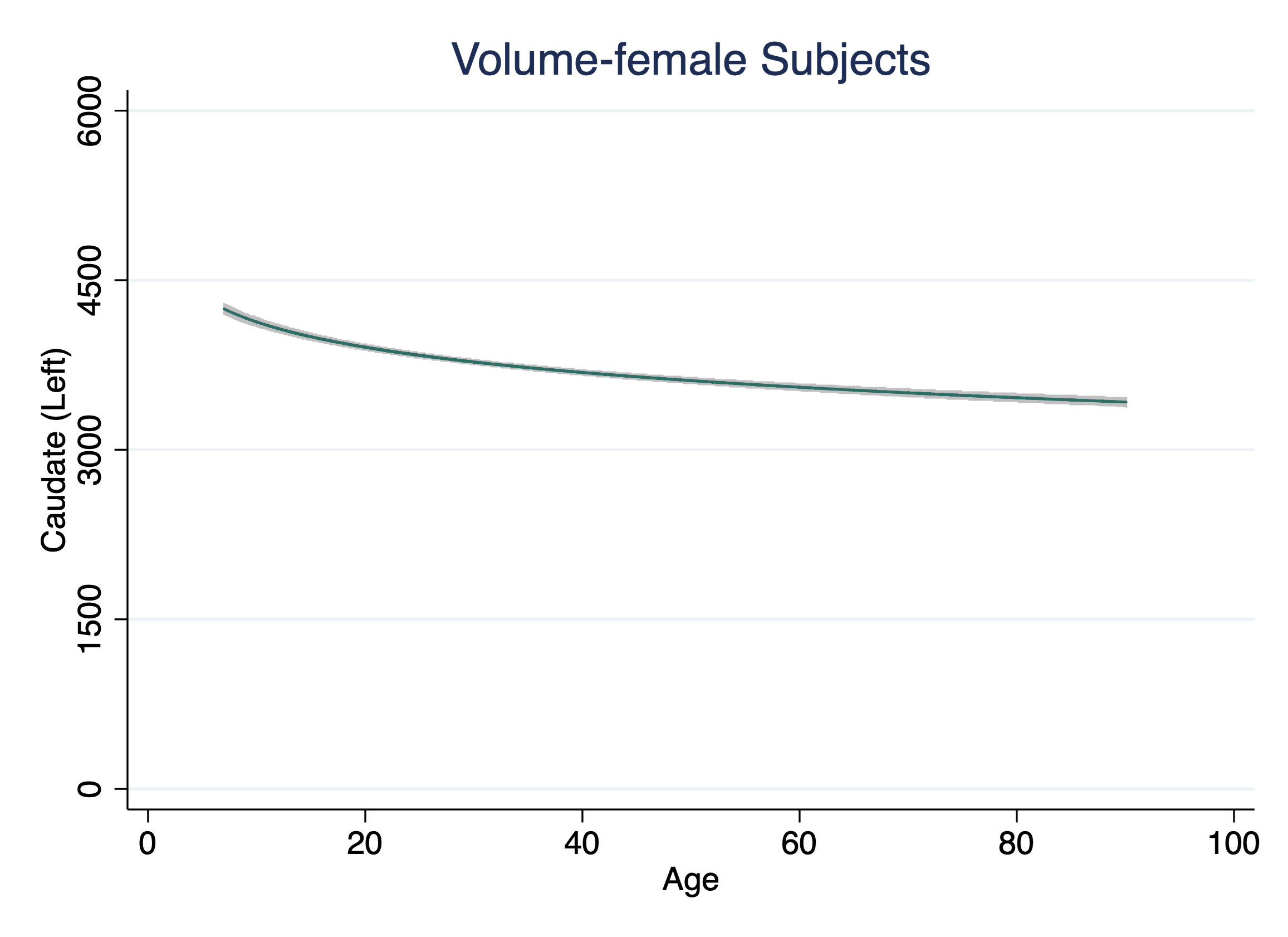

**Figure S4. Age-related Trajectories in Nucleus Accumbens**

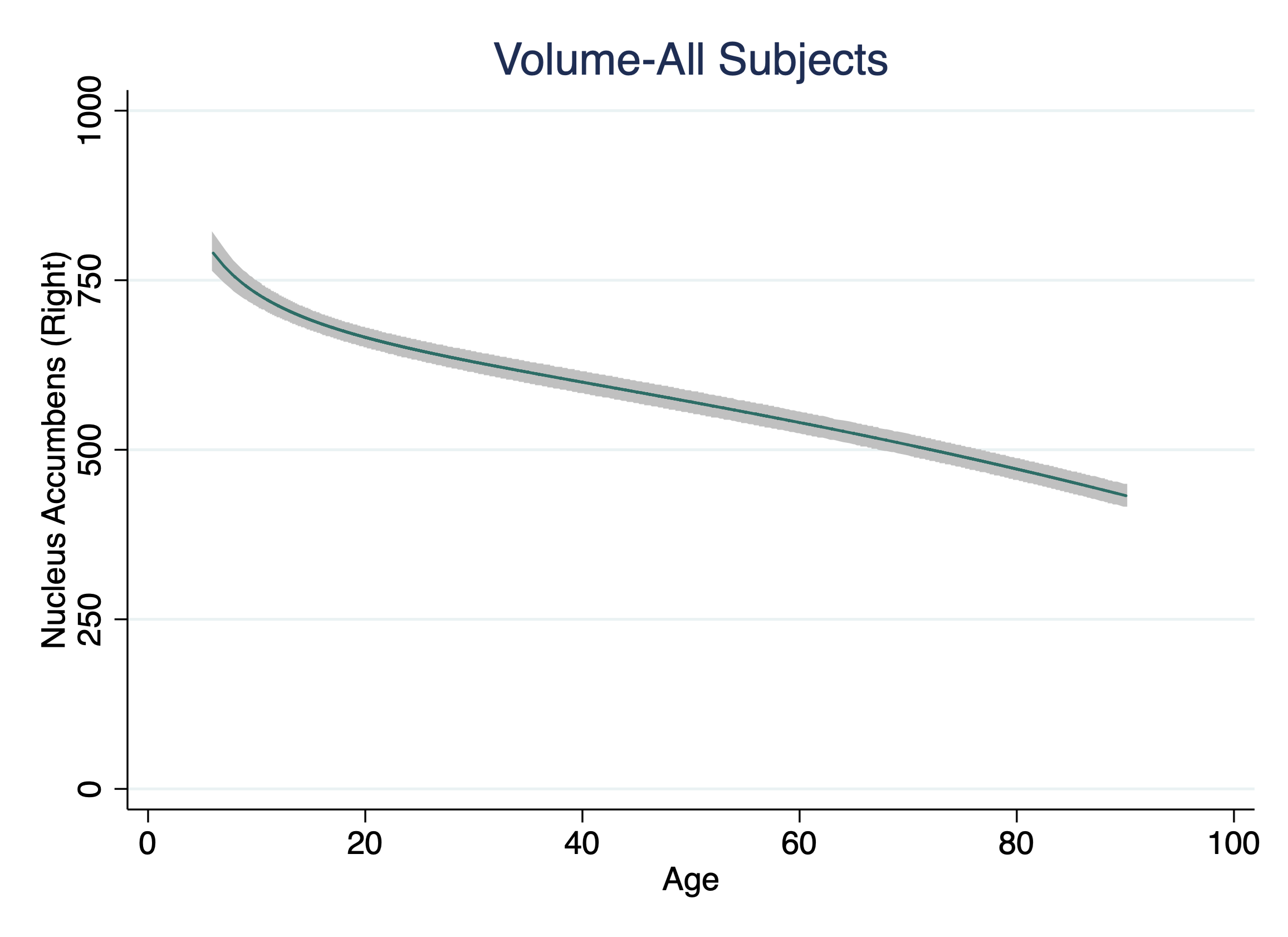

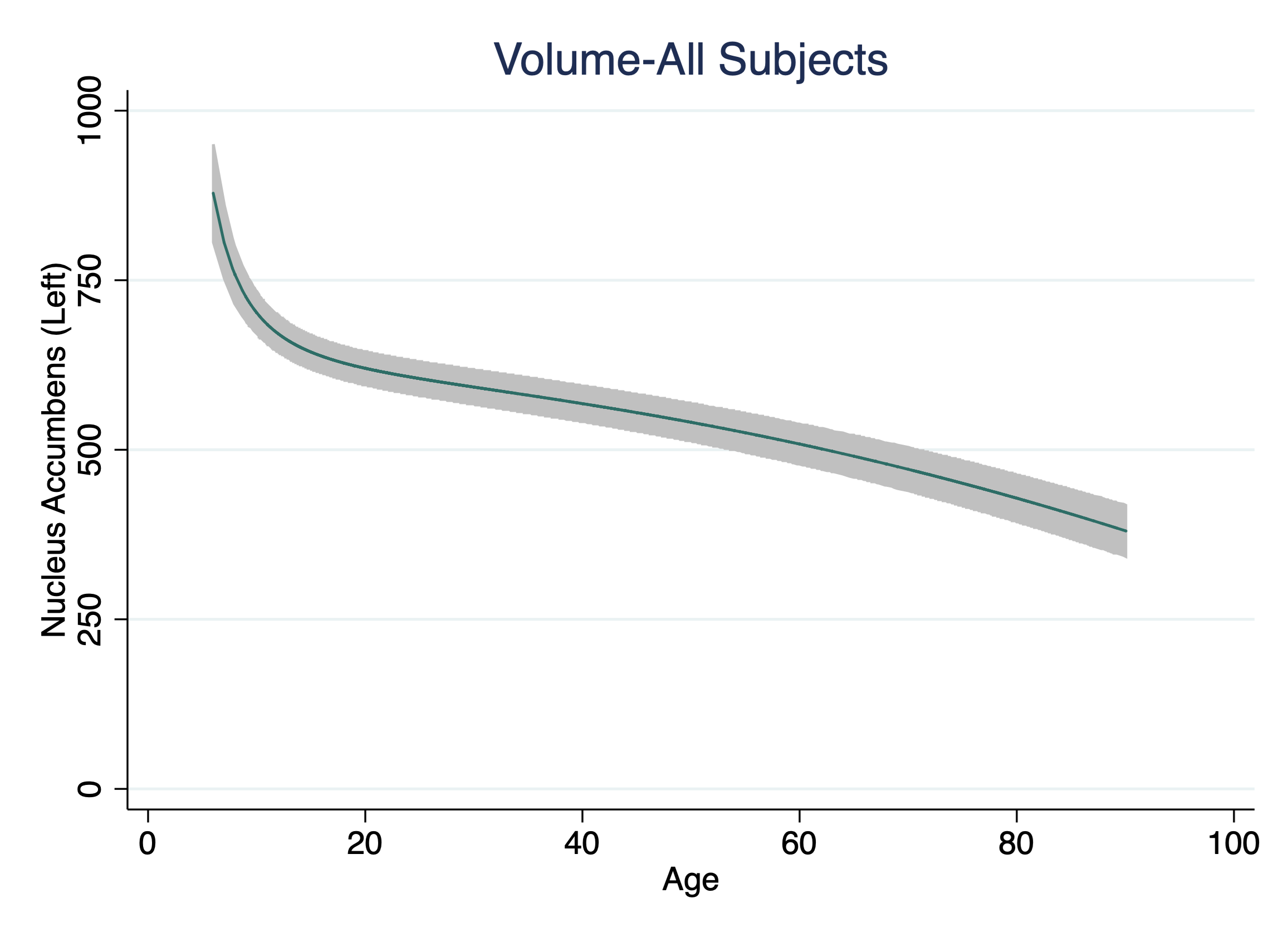

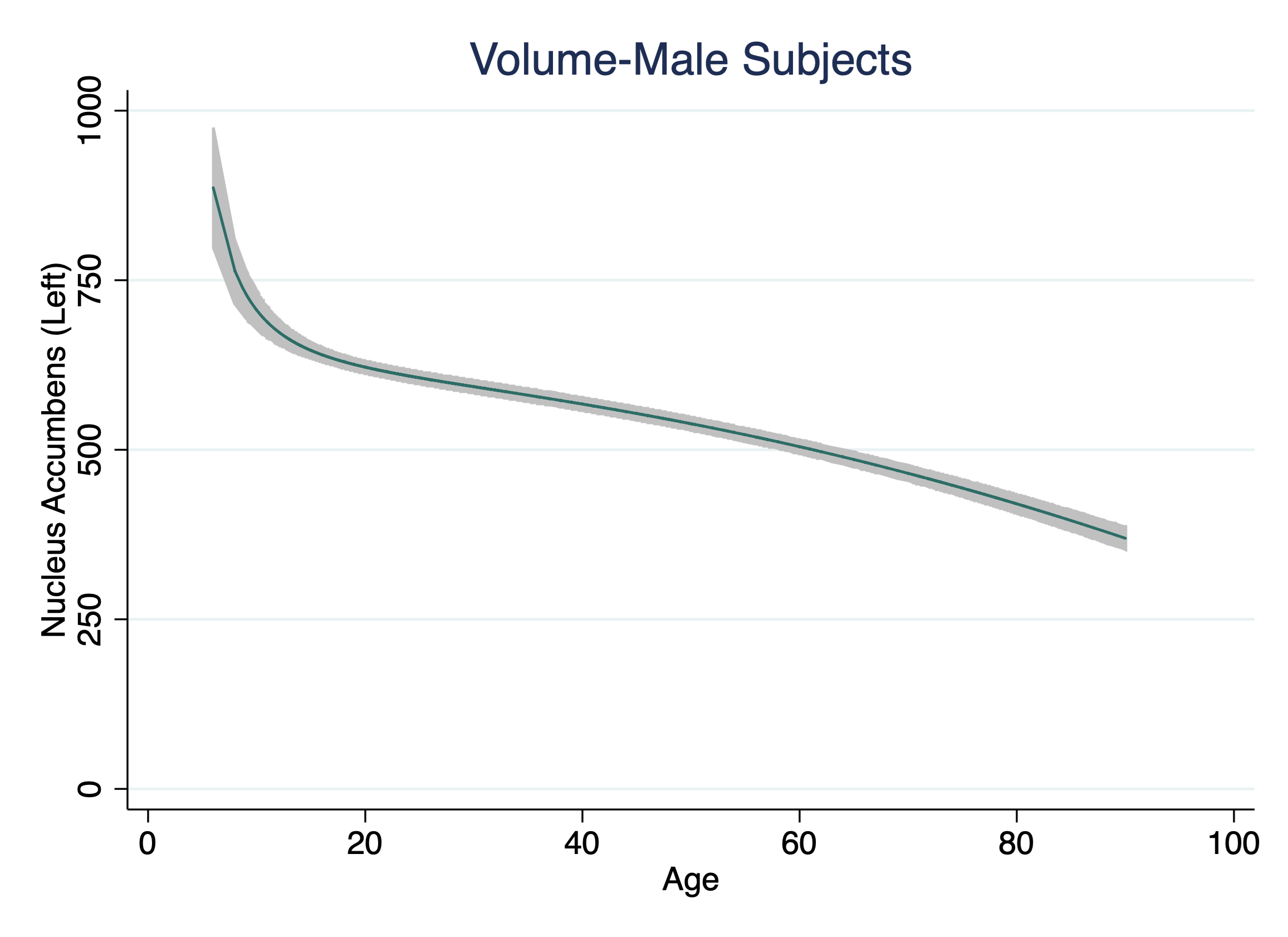

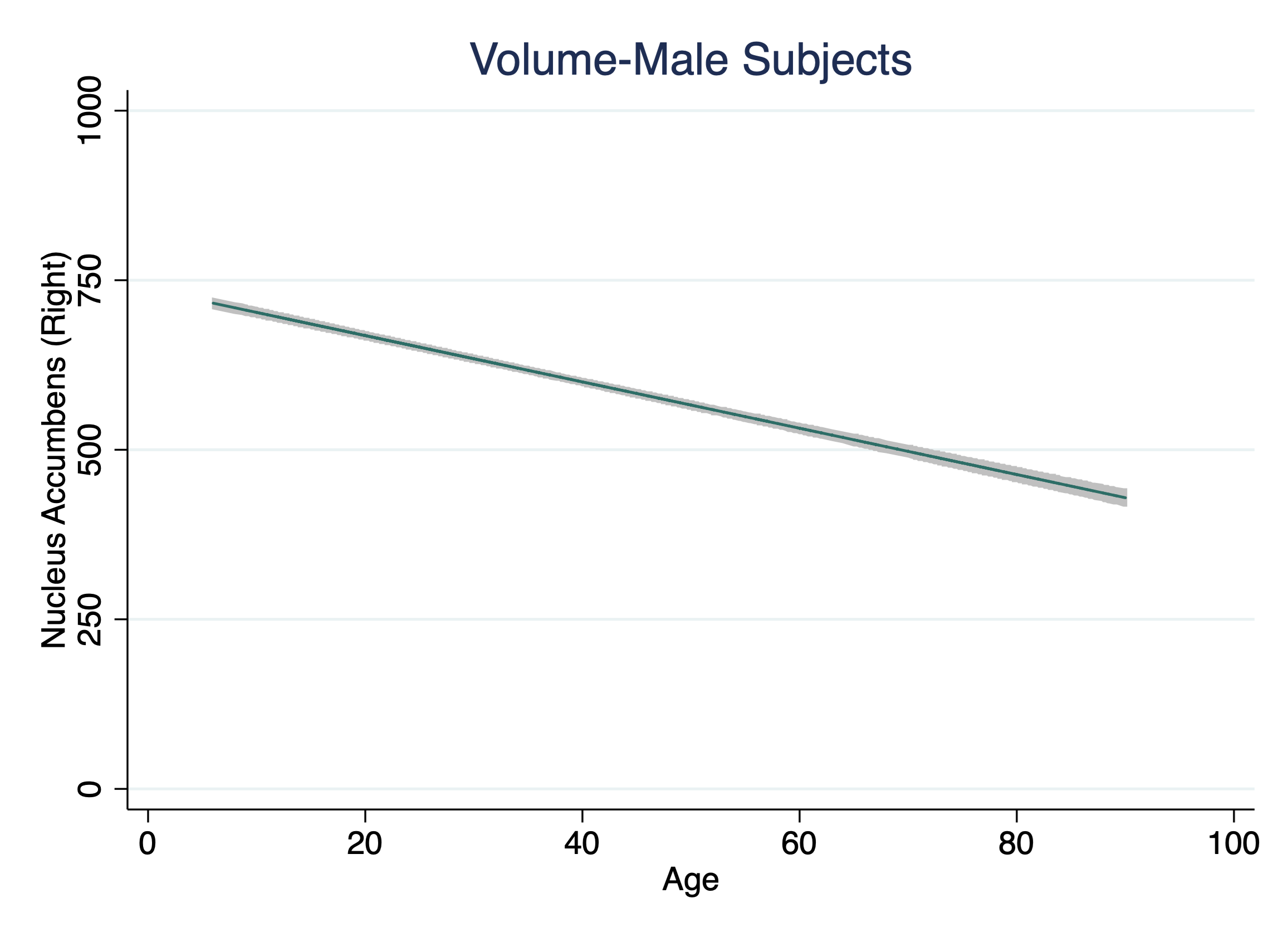

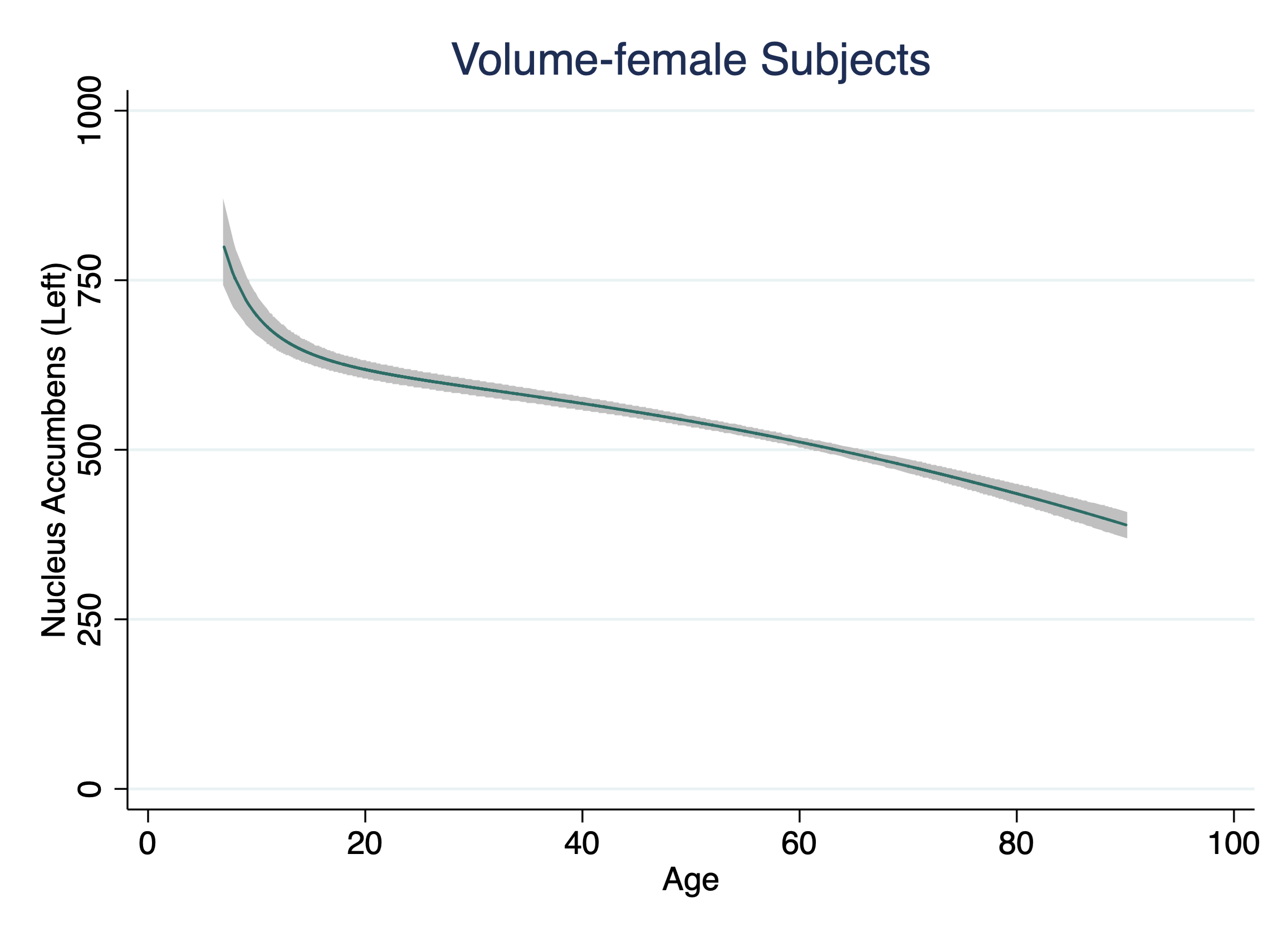

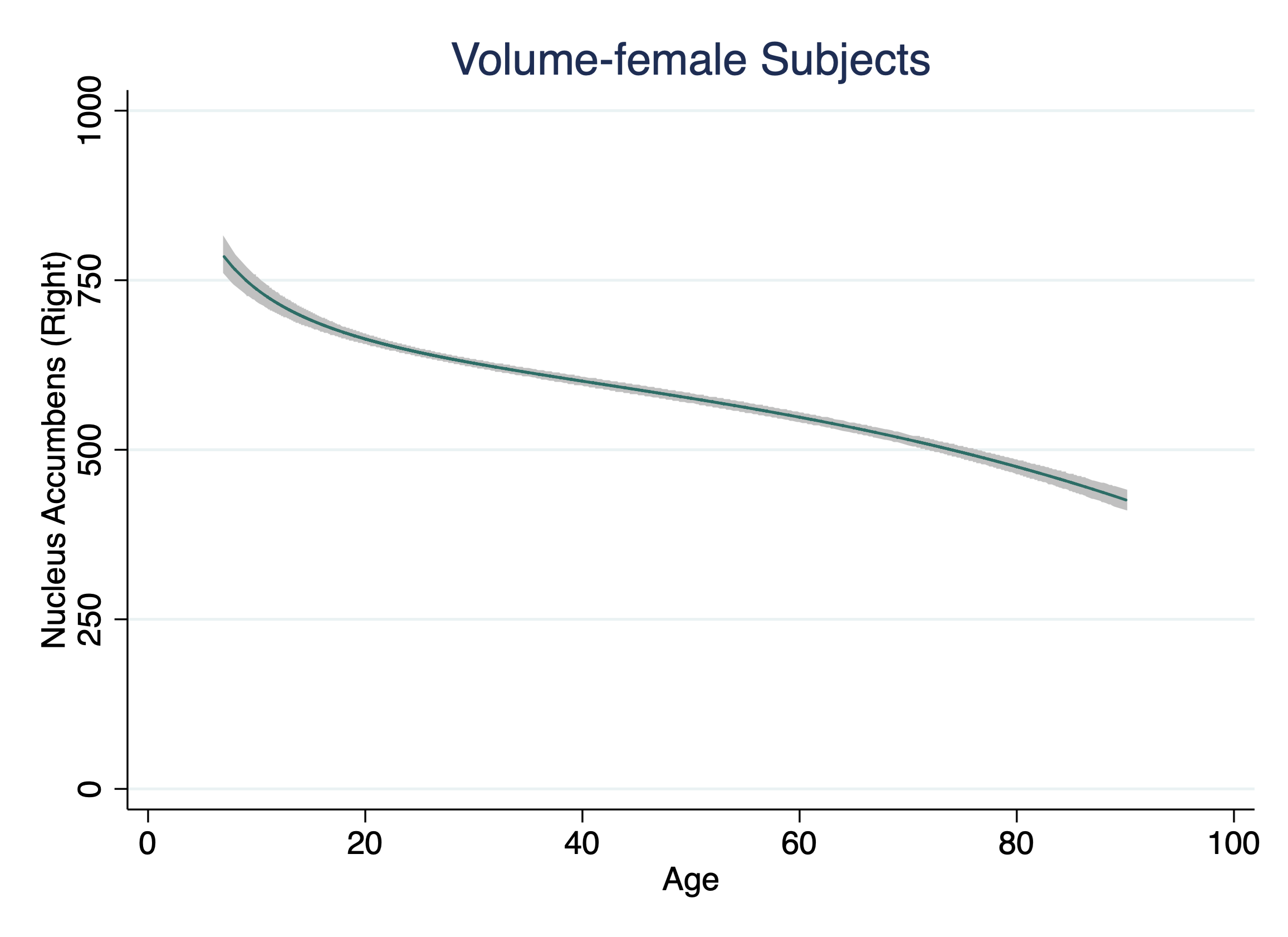

**Figure S5. Age-related Trajectories in Thalamus, Hippocampus, and Amygdala in Males**

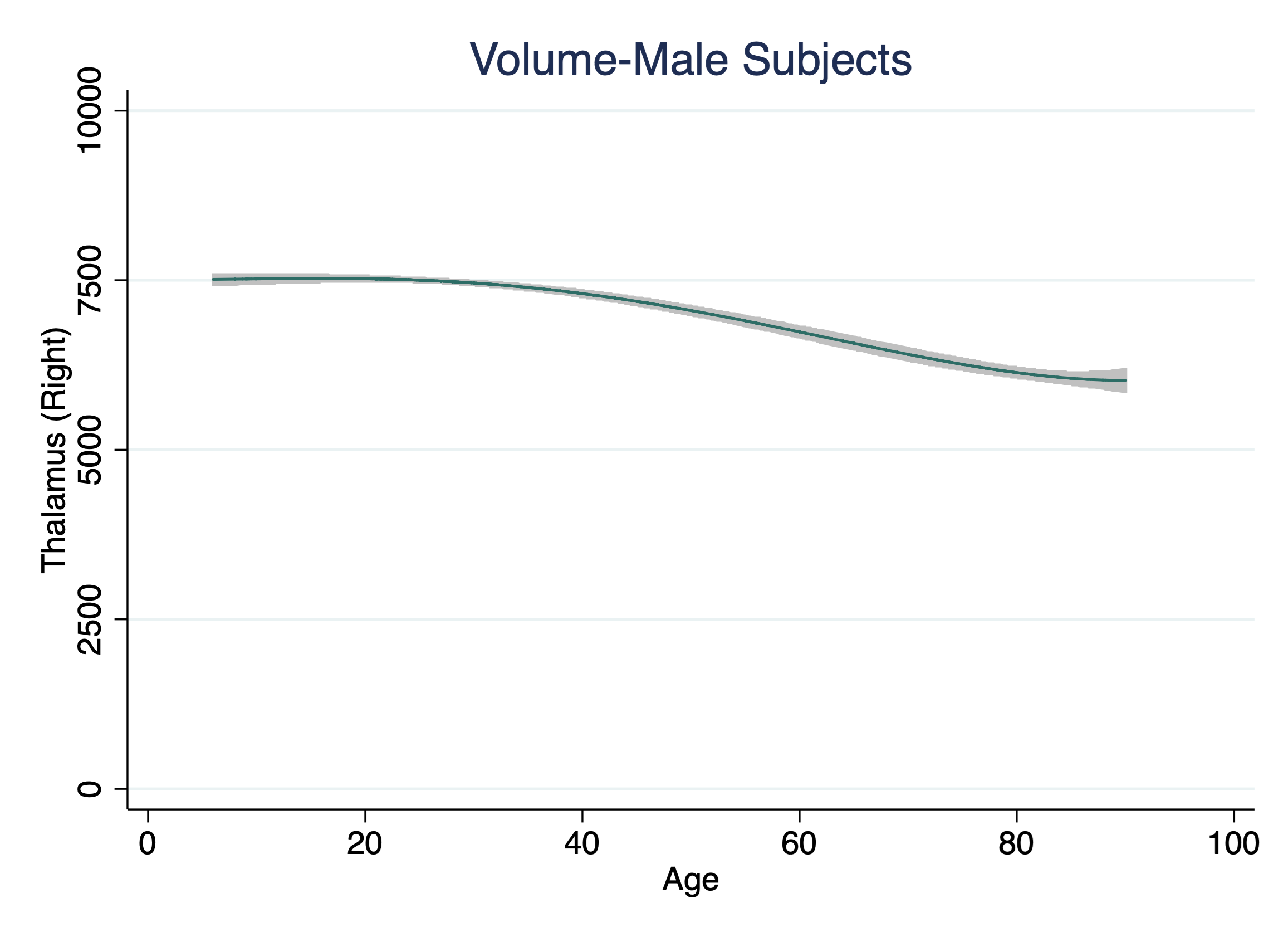

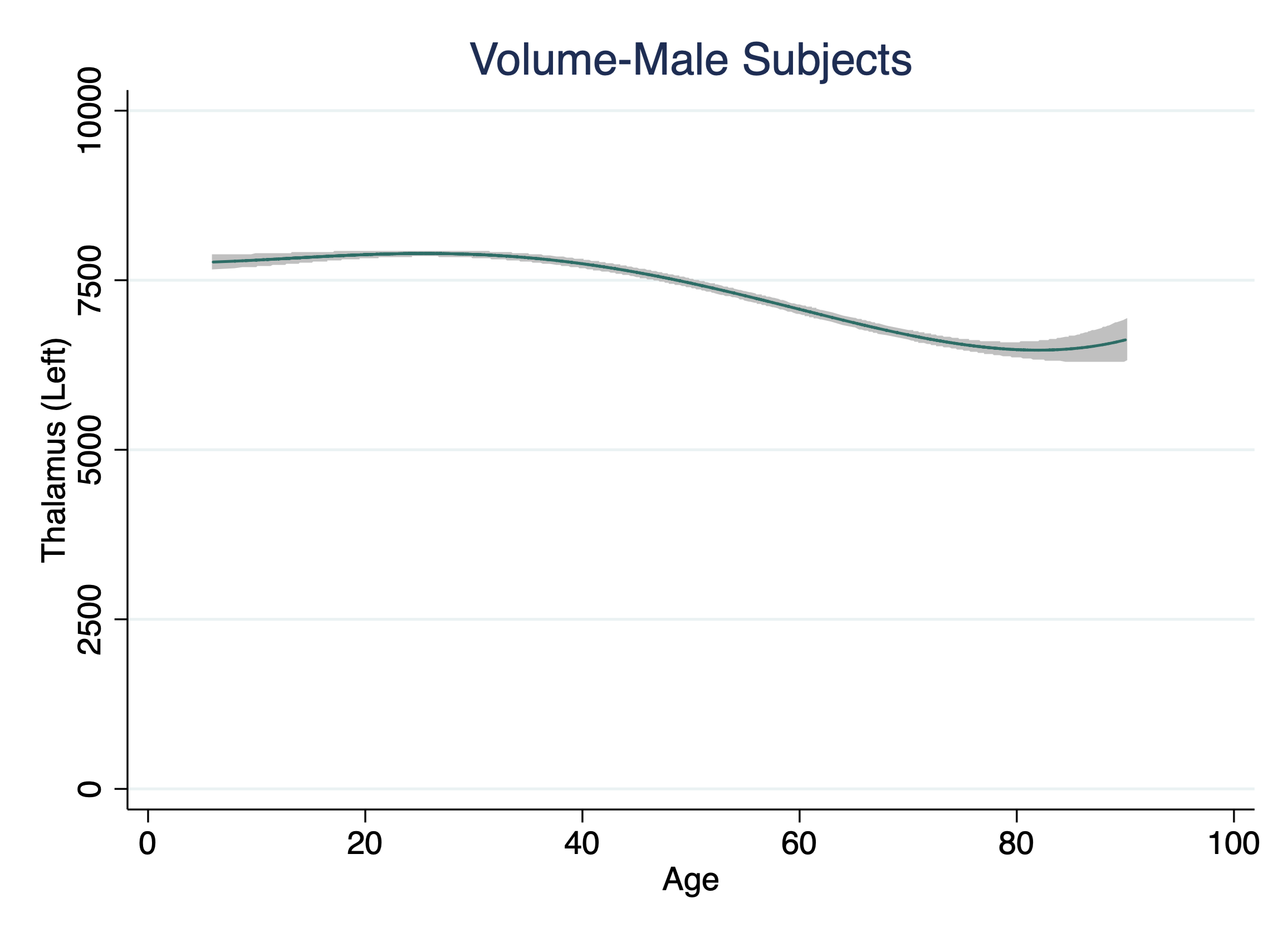

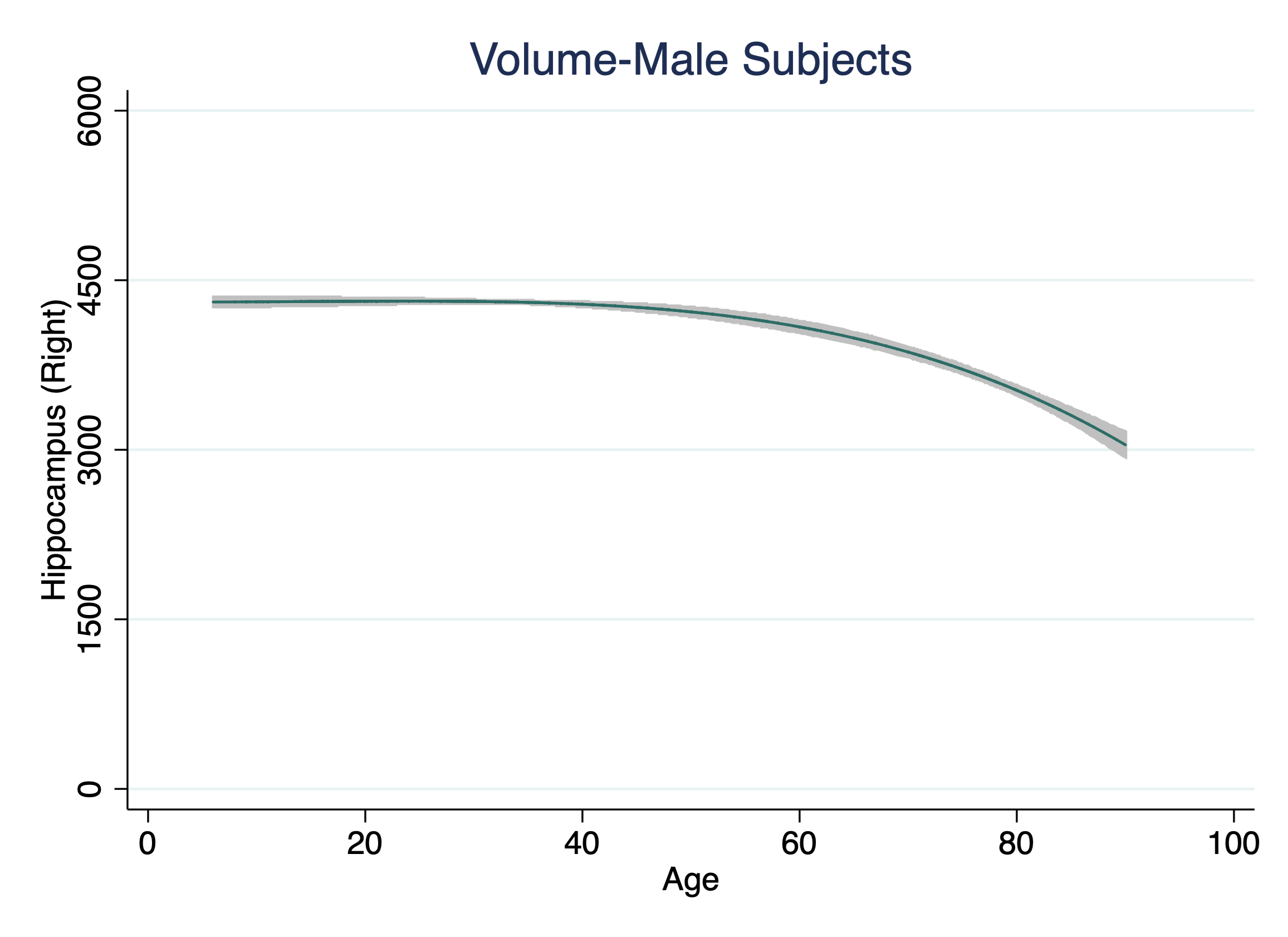

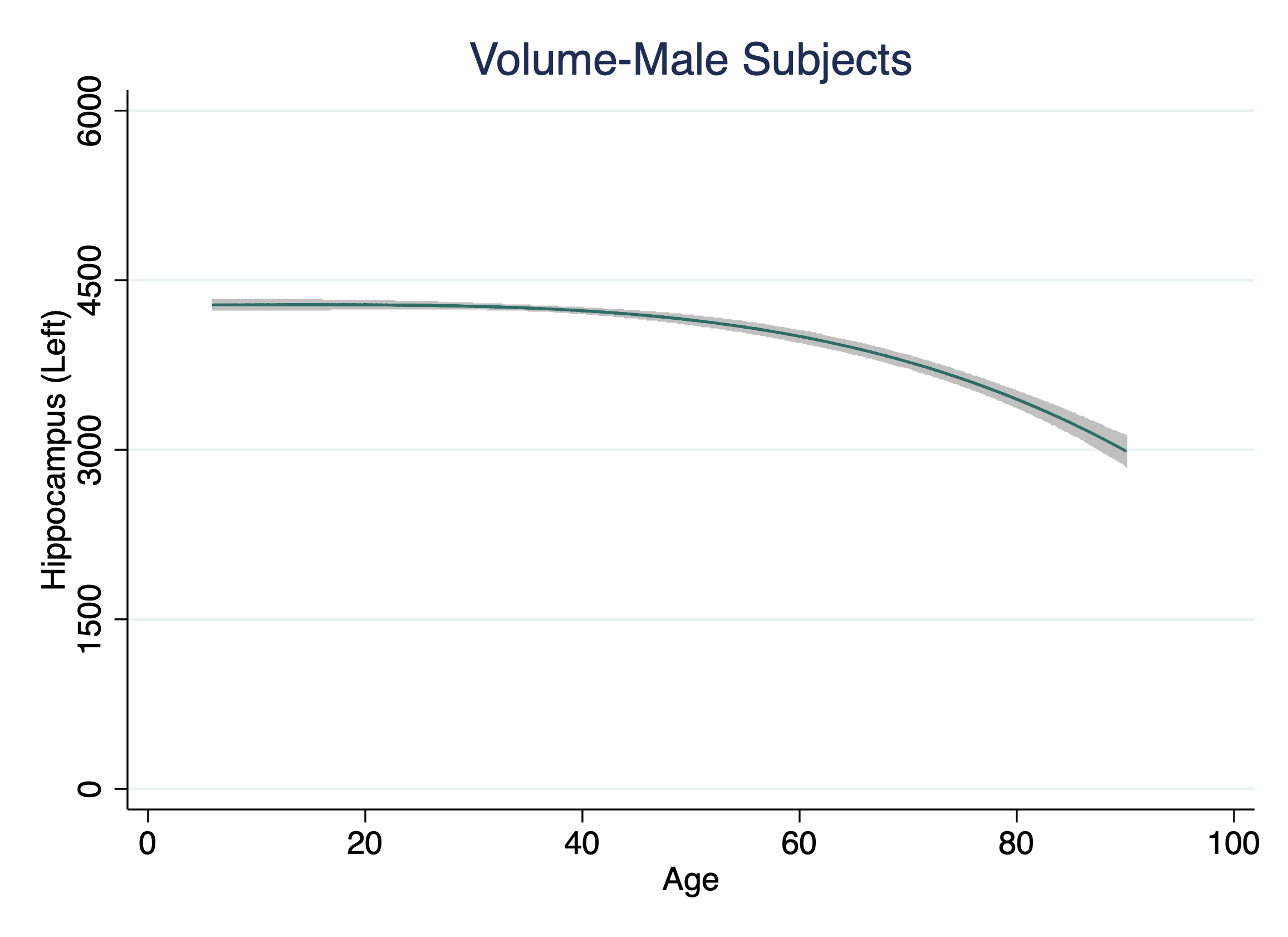

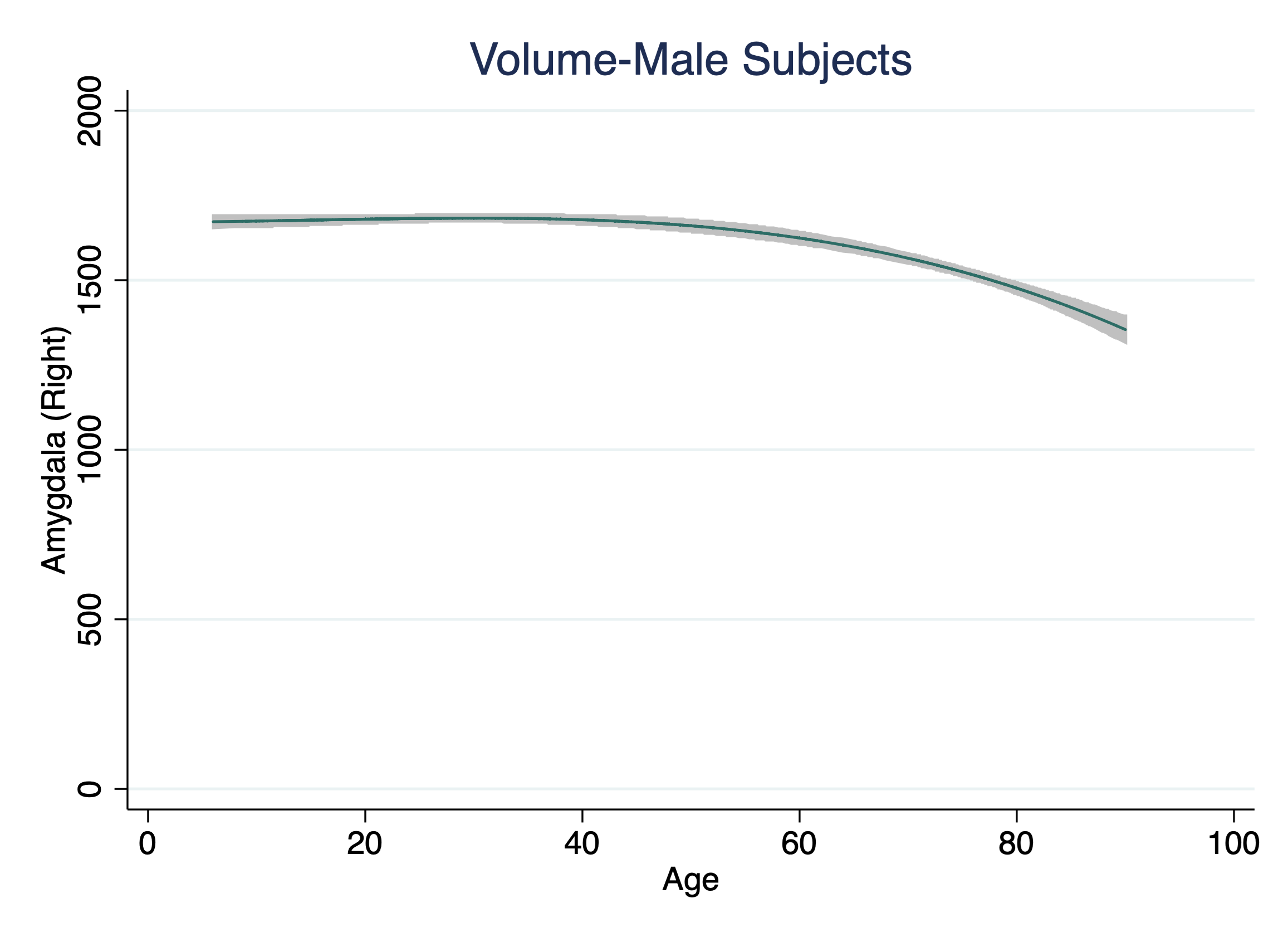

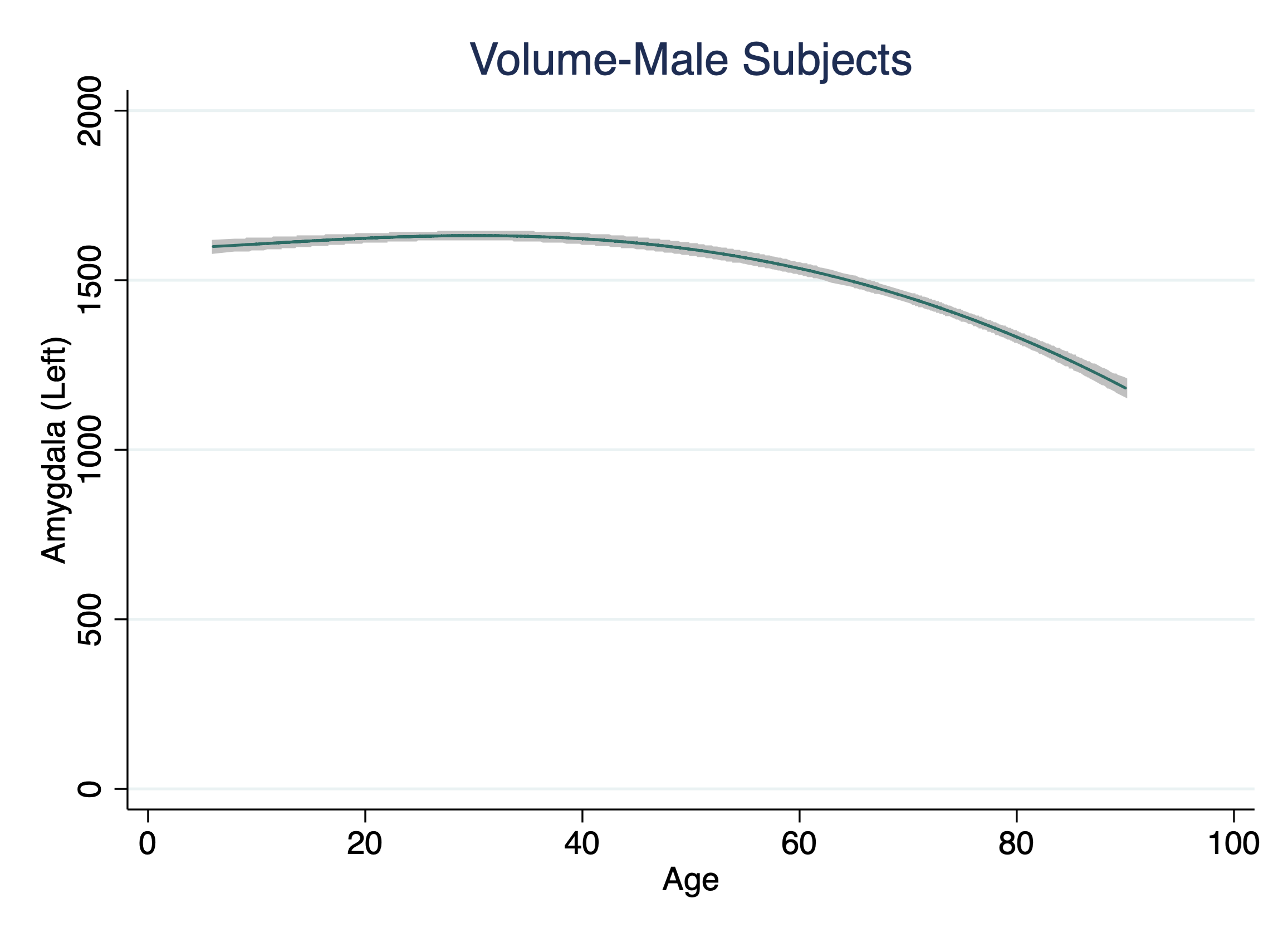

**Figure S6. Age-related Trajectories in Thalamus, Hippocampus, and Amygdala in Females**

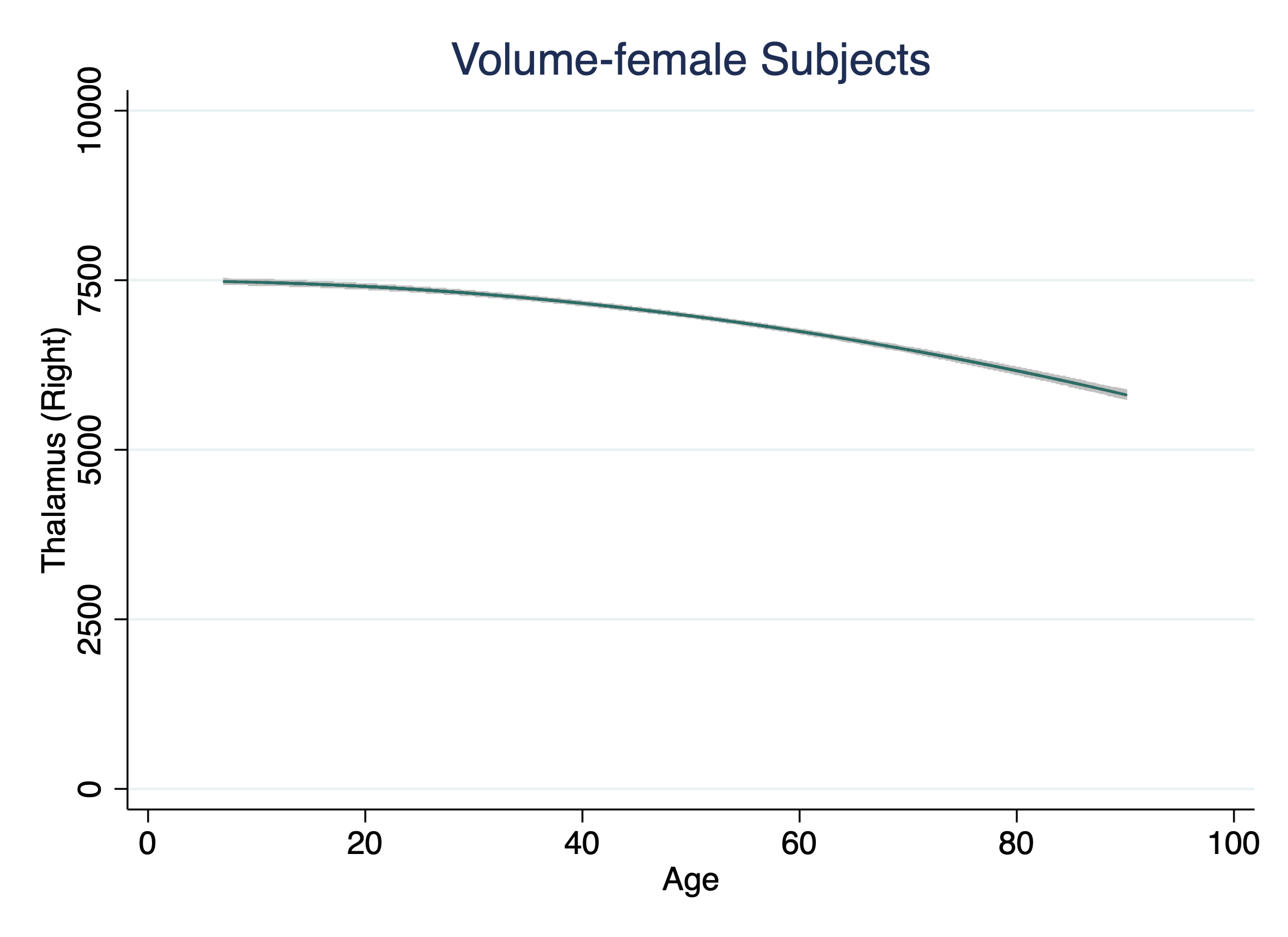

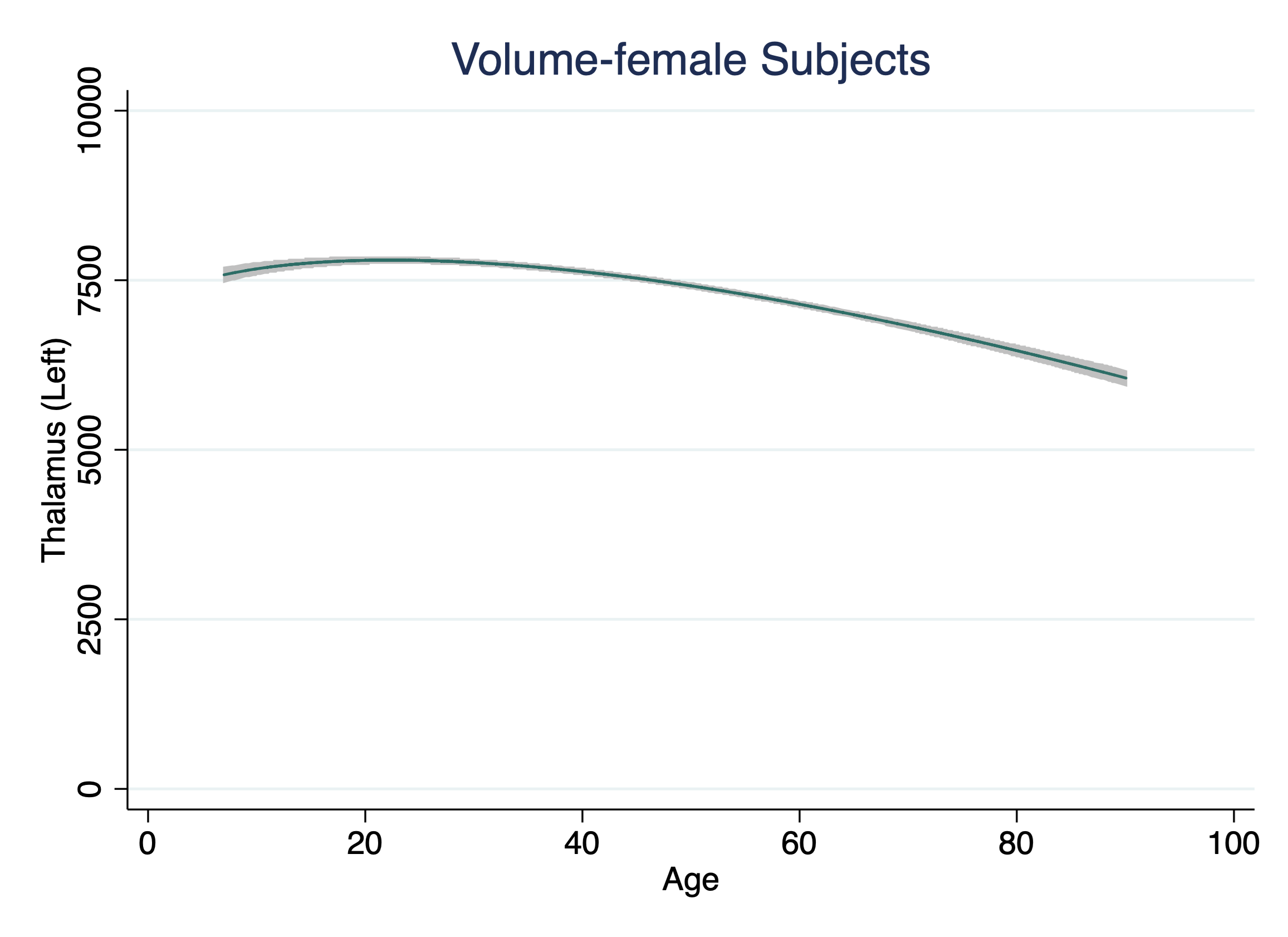

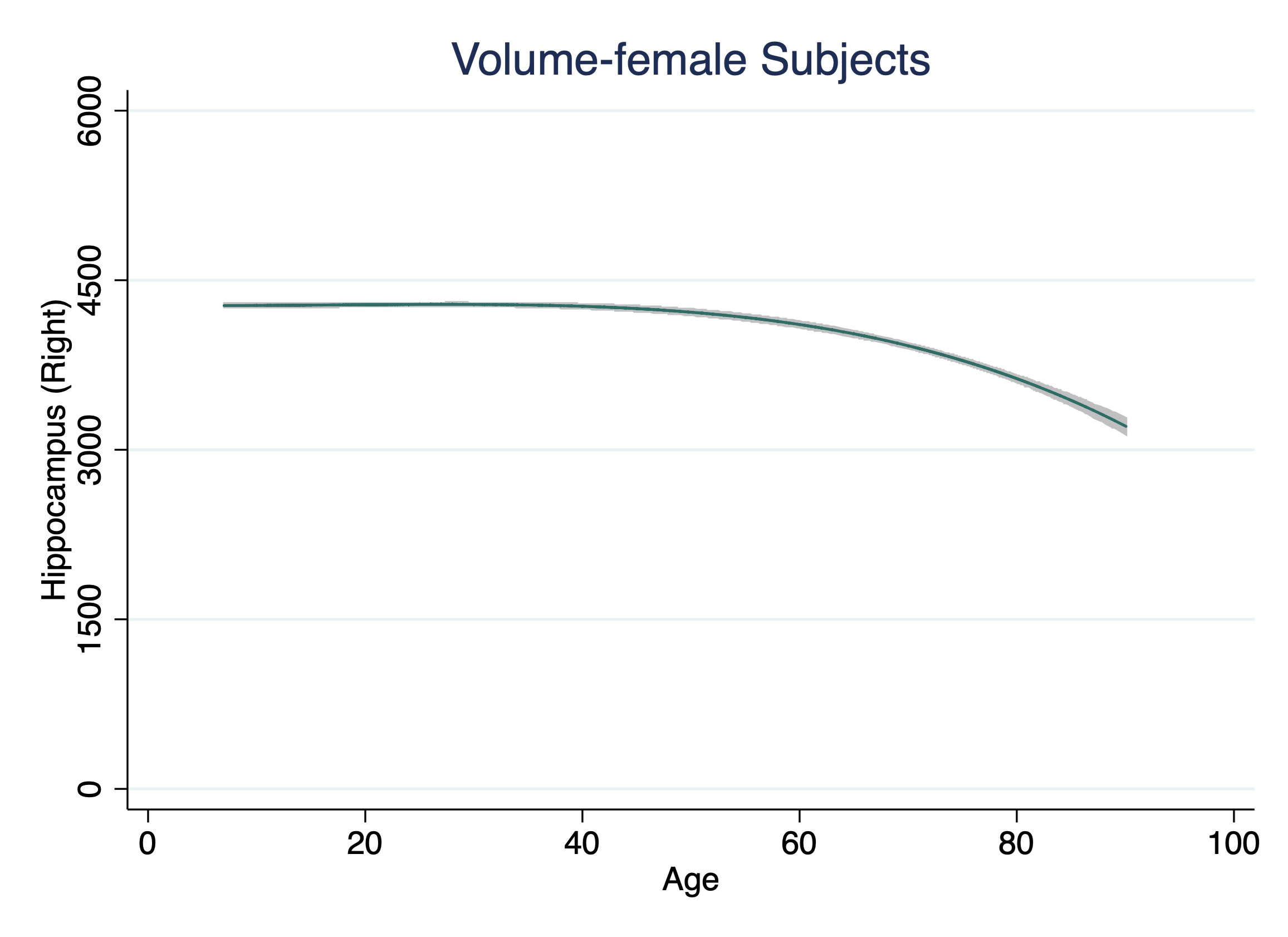

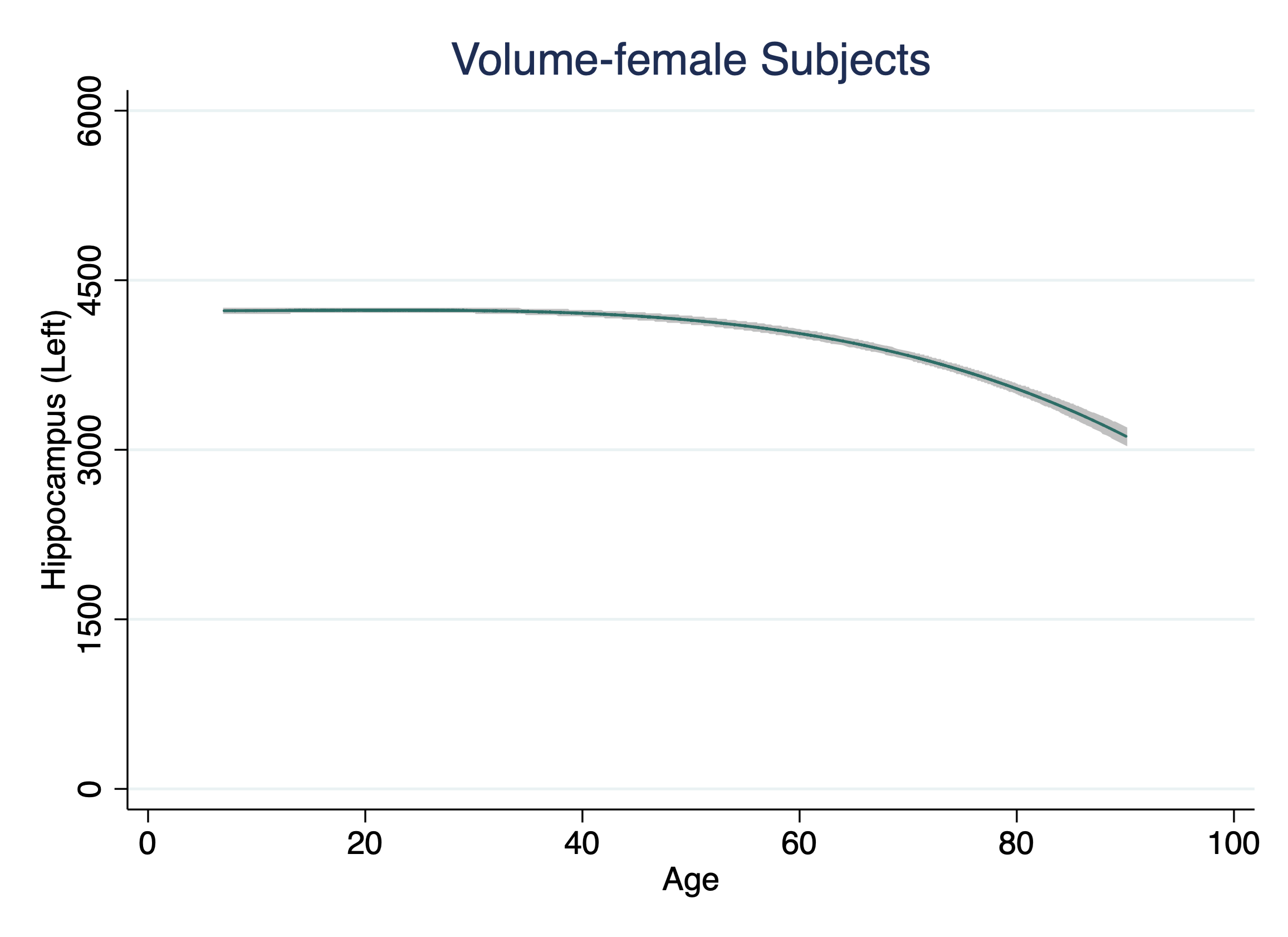

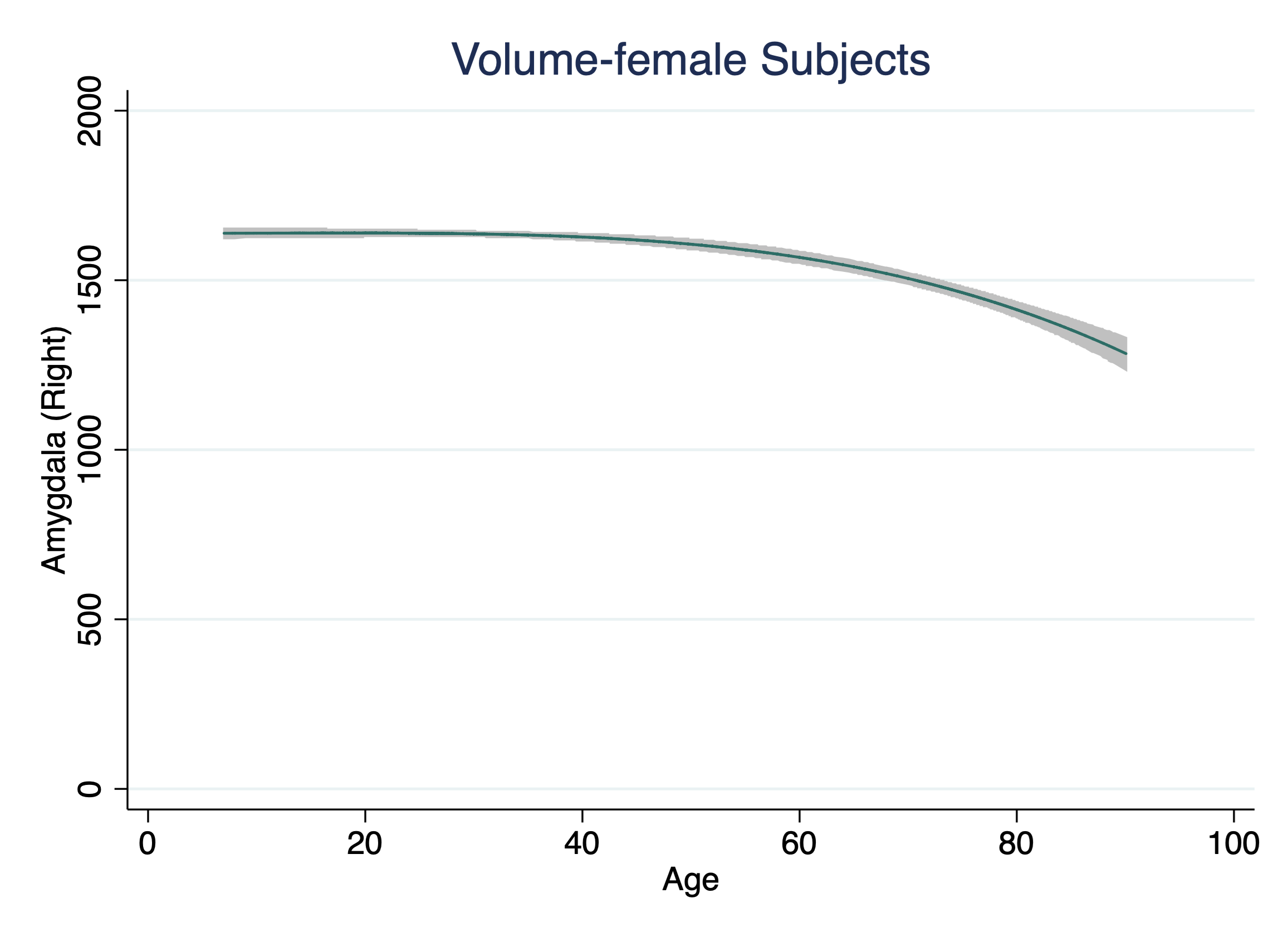

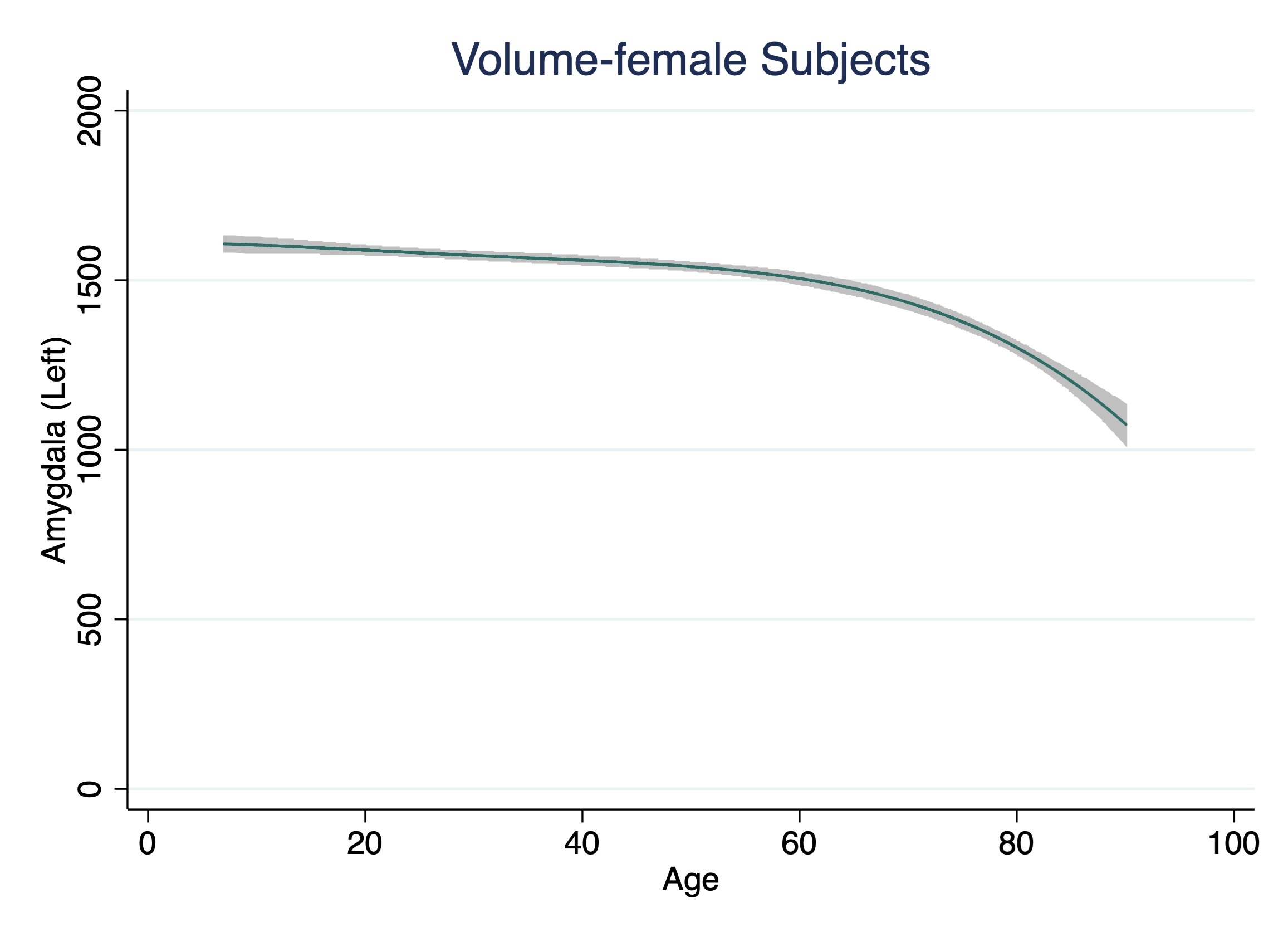

**Figure S7. Age-related Trajectories in Lateral Ventricles**

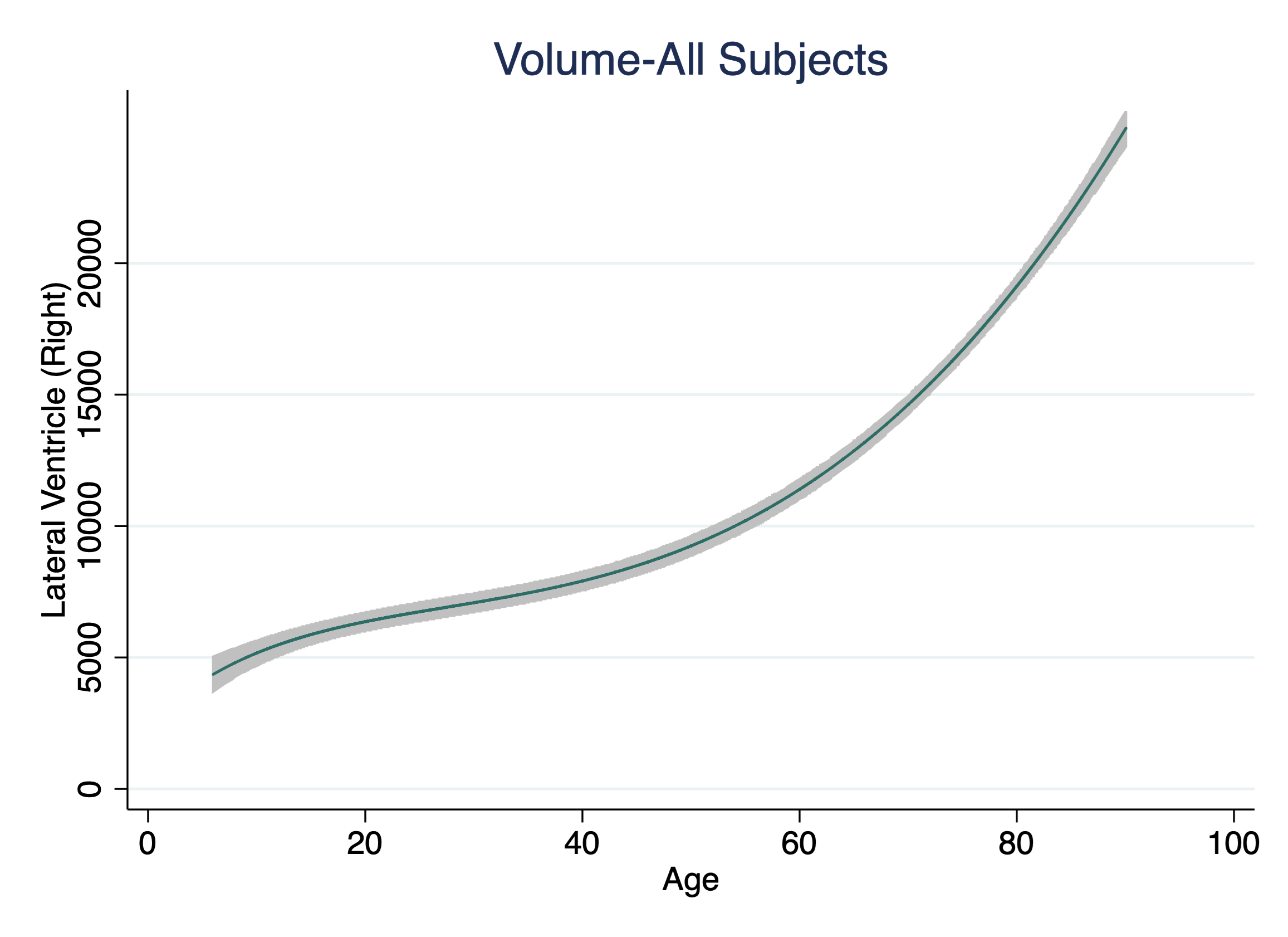

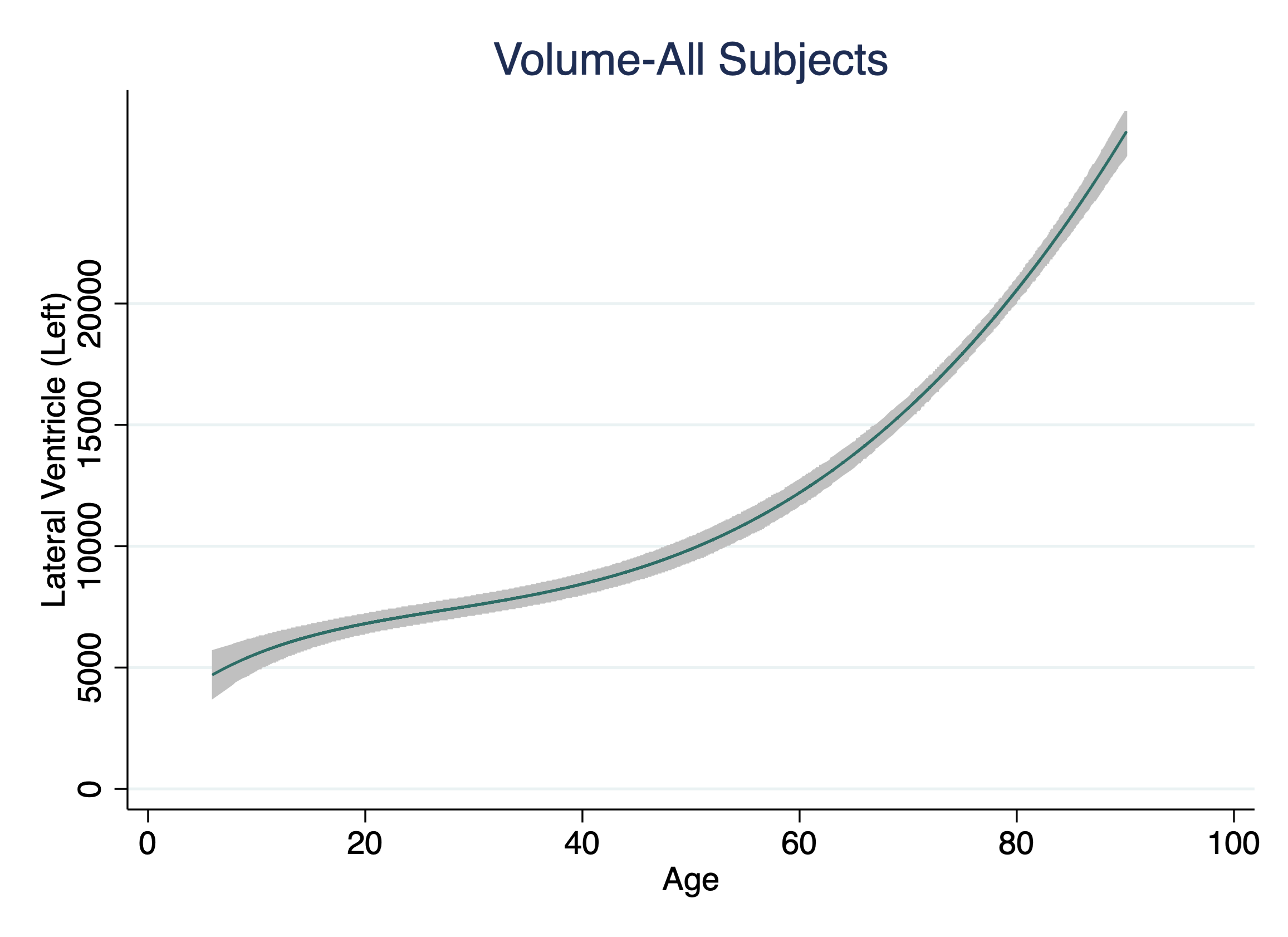

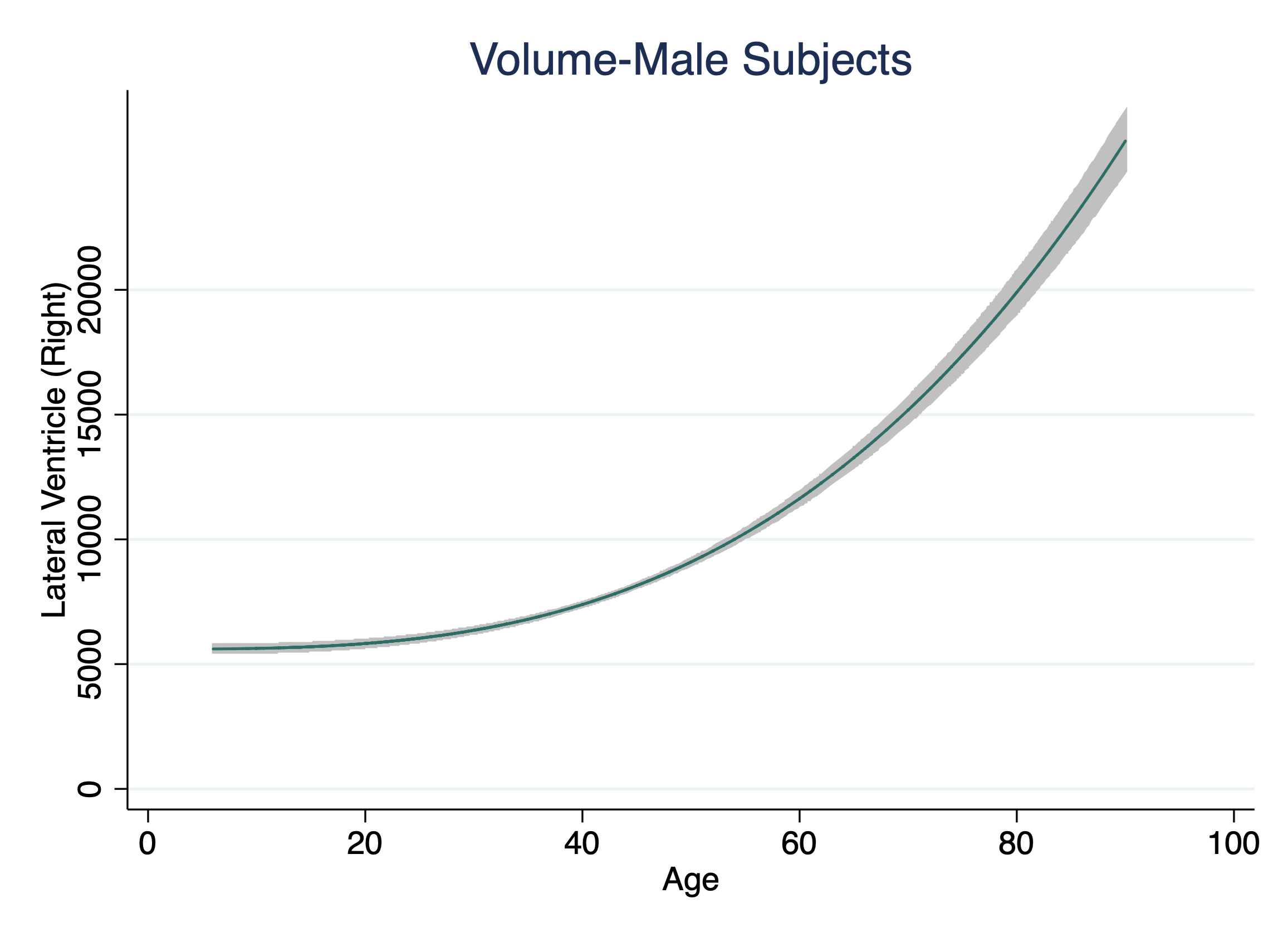

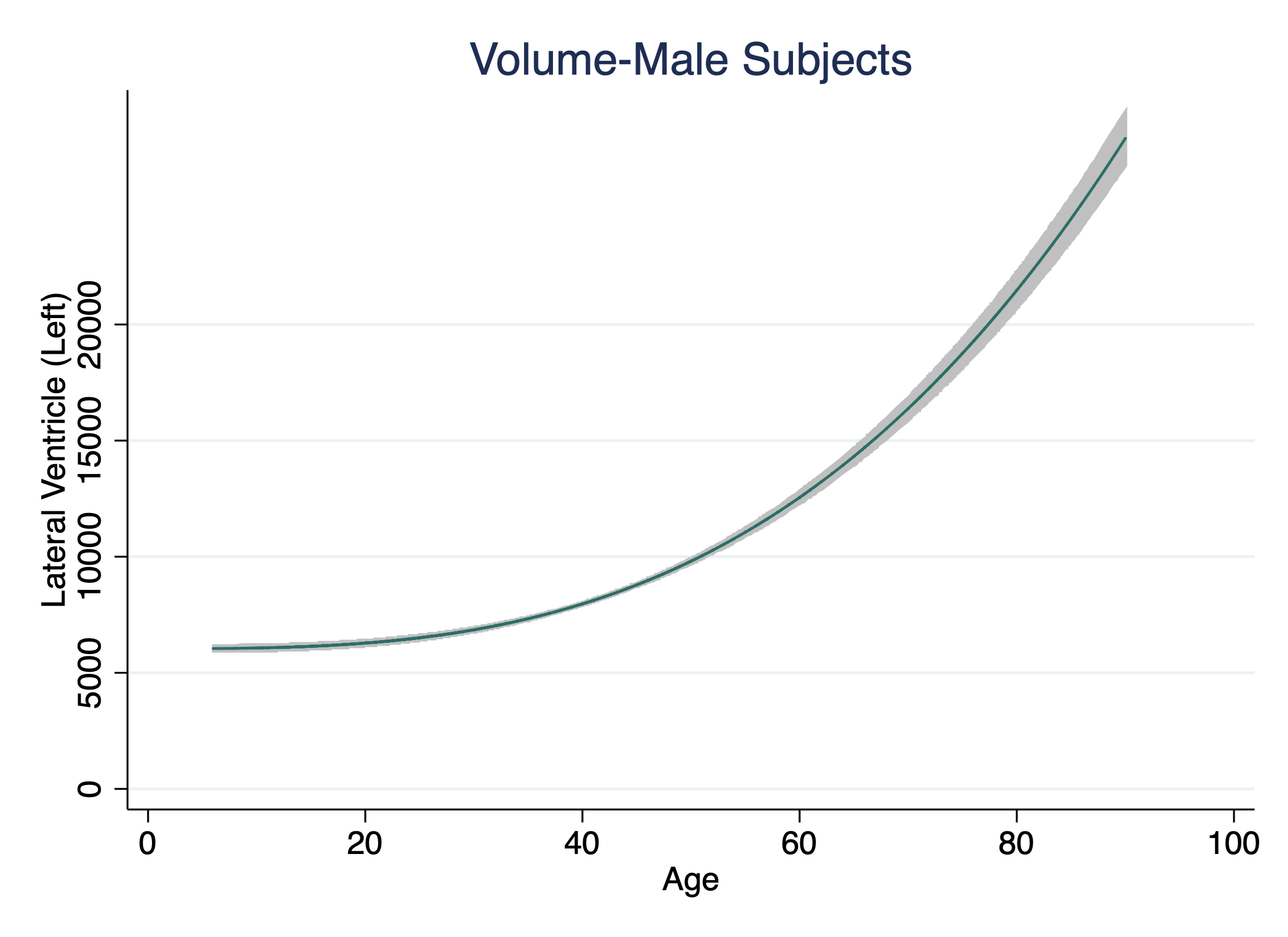

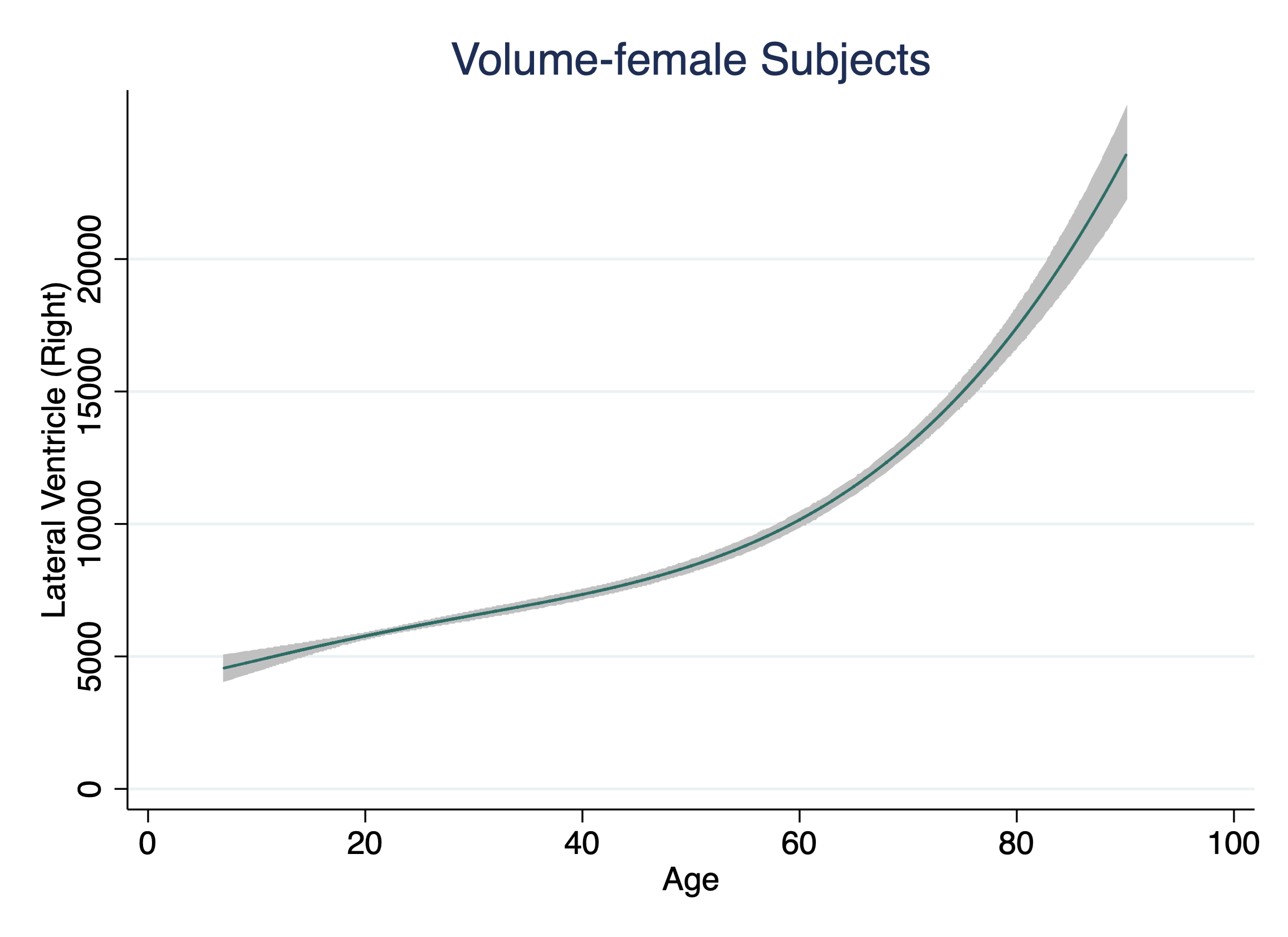

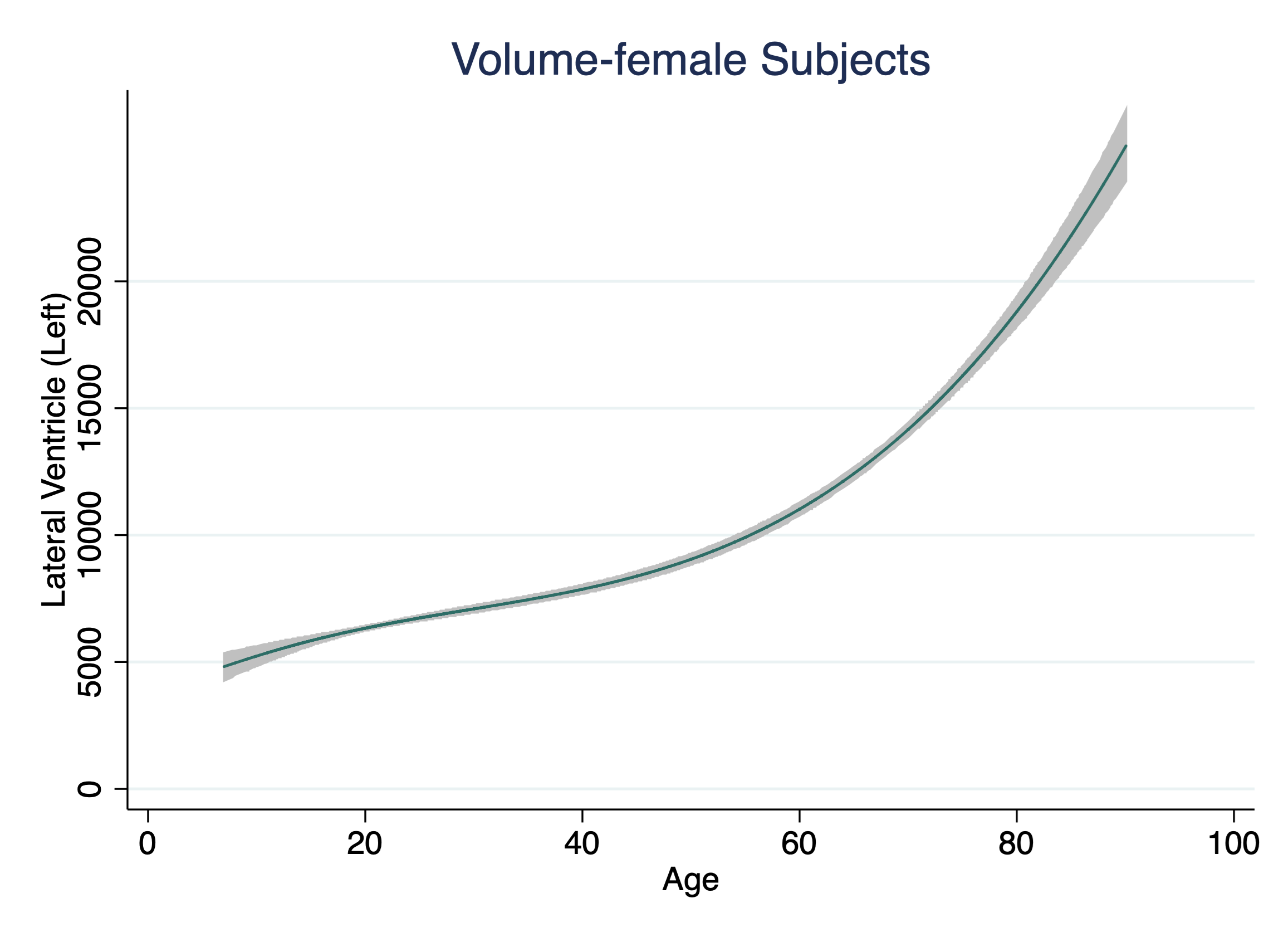

**Figure S8. Meta-analysis of the Pooled Standard Deviation of the Volume of each Subcortical Structure Stratified by Sex**

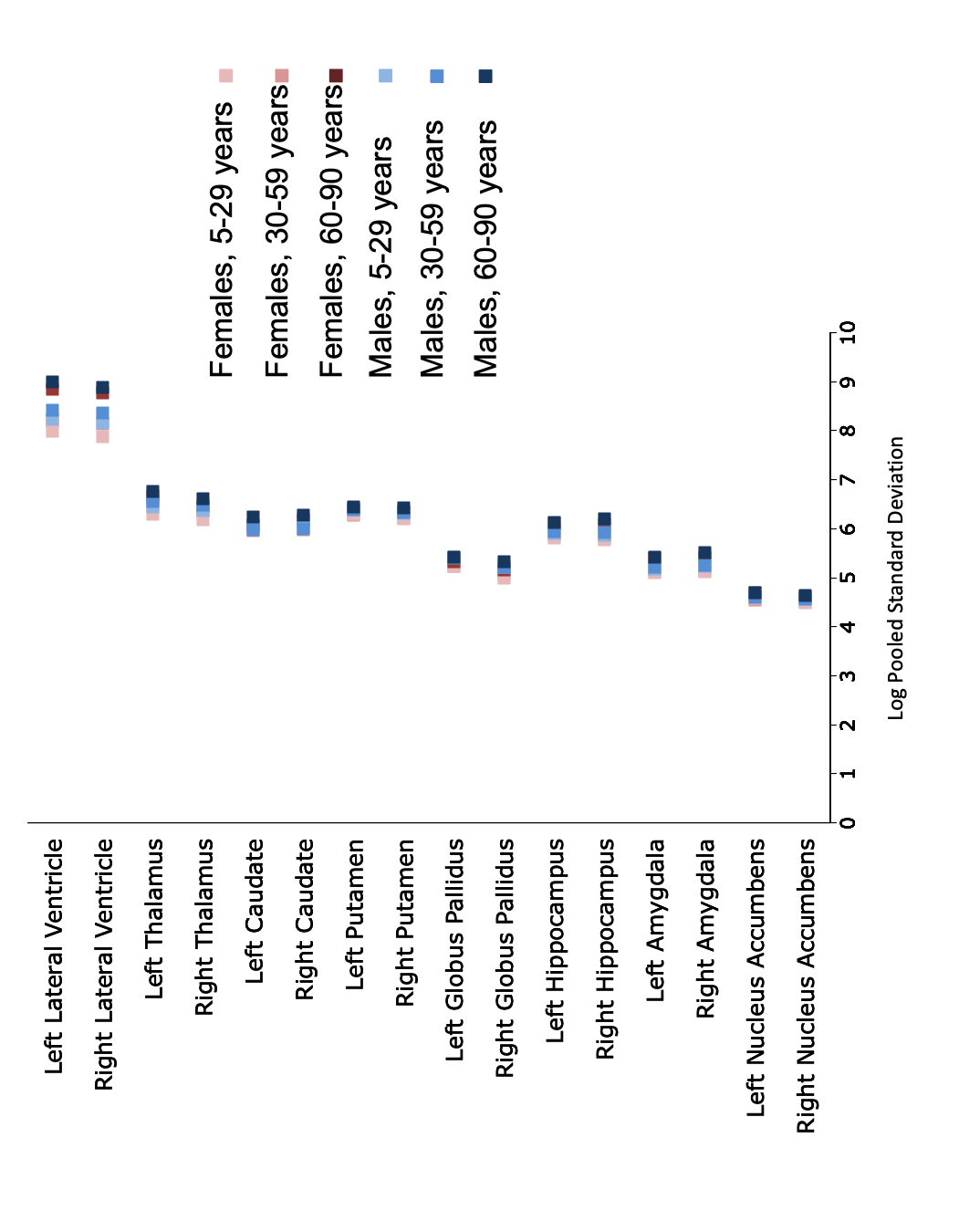

| **Table S1. Screening Process and Eligibility Criteria, Scanner, Image Acquisition Parameters and Image Segmentation Software** | | | | | |
| --- | --- | --- | --- | --- | --- |
| **Sample** | **Screening Process** | **Eligibility Criteria** | **Magnet strength/ Scanner Vendor** | **Acquisition parameters** | **Freesurfer version** |
| **ADHD-NF** | KSADS | No head trauma, no neurological and psychiatric history, no lifetime alcohol or substance abuse, no previous or current use of psychotropic medication, IQ>75. | 3T Siemens Tim Trio | T1-weighted 3D MPRAGE; TR/TE/TI/FA=2300ms/3030ms/900ms/9^o^;  Image matrix= 256 × 256; 192 sagittal slices; voxel size=1mm^3^ | 5.3 |
| **AMC** | Personal Interview | No head trauma, no medical, neurological or psychiatric history, no lifetime alcohol or substance abuse, no previous or current use of psychotropic medication, IQ>75. No history of any psychiatric disorders in 1st degree family members. | 3T Philips Intera | T1-weighted 3D MPRAGE; TE range= 3.5-4.6ms, TR range= 9-9.663ms, FA= | 5.1 |
| **Barcelona 1.5T** | KSADS | No head trauma, no medical, neurological or psychiatric history, no lifetime alcohol or substance abuse, no previous or current use of psychotropic medication, IQ>75. No history of any psychiatric disorders in 1st degree family members. | 1.5 T General Electric Signa | T1-weighted; image matrix = 256 x 256; 128 slices; voxel size 1 x 1 x 1 mm^3^. | 5.3 |
| **Barcelona 3T** | KSADS | No head trauma, no medical, neurological or psychiatric history, no lifetime alcohol or substance abuse, no previous or current use of psychotropic medication, IQ>75. No history of any psychiatric disorders in 1st degree family members. | 3 T Siemens MAGNETOM TIM Trio | T1-weighted; image matrix = 256 x 256; 240 slices; voxel size 1 x 1 x 1 mm^3^. | 5.3 |
| **Betula** | Personal Interview | No head trauma, no medical, neurological or psychiatric history, no lifetime alcohol or substance abuse, no previous or current use of psychotropic medication. | 3T General Electric Discovery  MR750 | T1-weighted MPRAGE; TR/TE/TI/FA=8.1240 ms/3.2000 ms/450ms/ 12°; image matrix = 256x256 | 5.3 |
| **BIG 1.5T and 3T** | Questionnaire for psychiatric history | No head trauma, no medical, neurological or psychiatric history, no lifetime alcohol or substance abuse, no previous or current use of psychotropic medication, IQ>70. No history of any psychiatric disorders in 1st or 2^nd^ degree family members. | 1,5 T Siemens Sonata and Avanto and 3 T Siemens Trio, TimTrio and Skyra | T1-weighted 3D MPRAGE; TR/TE/TI/sagittal slices = 1940-2730 ms/850-110 ms/2.92-4.58 ms; 176-192 sagittal slices; 1.0x1.0x1.0 mm^3^ | 5.3 |
| **BIL&GIN** | Personal Interview | No head trauma, no current neurological or psychiatric disorders, no current use of psychotropic medication, IQ>70. | 3T Phillips ACHIEVA | T1 - weighted 3D; TR/TE/TI/FA=20 ms/4.6 ms/800ms/ 10°; turbo field echo factor = 65; sense factor = 2; matrix size = 256x256x180mm 3 ; voxel size= 1.0x1.0x1.0 mm^3^ | 5.3 |
| **CAMH** | SCID | No head trauma, no neurological or psychiatric history, no alcohol or substance abuse preceding 6 months, no previous or current use of psychotropic medication, IQ>75. No history of any psychotic disorders in 1st degree family members. | 1.5 T GE (echospeed) | 124 Axial inversion recovery–prepared spoiled gradient recall images, 1.5-mm-thick slice acquisition TE=5.3ms; TR=12.3ms; time to inversion, 300.0ms; flip angle, 20°. | 5.3 |
| **Cardiff** | MINI | No head trauma, no medical history, including neurological and psychiatric history, no alcohol or substance abuse in the preceding 6 months, no previous or current use of psychotropic medication. | 3T General Electric Signa | T1-weighted 3D FSPGR; TR/TE/TI/FA=7.9ms/3.0ms/450ms/20^o^;  Image matrix= 256 × 192 x 172; voxel size=1mm^3^ | 5.3 |
| **CEG** | Teacher and parent  Conners' | No head trauma, no medical history, including neurological and psychiatric history, no alcohol or substance abuse in the preceding 6 months, no previous or current use of psychotropic medication. IQ>75 | 3T General Electric Signa | T1-weighted 3D SPGR; TR/TE/ FA=2000ms/30ms/90^o^;  Image matrix= 128 × 128; 43 slices | 5.3 |
| **CIAM** | SCID | No head trauma or psychiatric history, no previous or current use of psychotropic medication, IQ>75. | 3T Siemens Allegra | T1-weighted 3D MPRAGE; TR/TE/TI/FA= 2530 ms/1.53, 3.21, 4.89, 6.57/2.91 ms/ ms/7^o^; image matrix= 256x256; 128 sagittal slices; voxel size= 1.3x1.0x1.3 mm^3^ | 5.3 |
| **CLiNG** | Personal Interview | No head trauma, no medical, neurological or psychiatric history, no lifetime alcohol or substance abuse, no previous or current use of psychotropic medication, IQ>75. No history of any psychiatric disorders in 1st degree family members. | 3T Siemens Tim Trio | T1-weighted 3D MPRAGE; TR/TE/TI/FA=2250 ms/3.26 ms/900 ms/9°; image matrix = 256 x 256; 192 sagittal slices; voxel size= 1 mm^3^ | 5.3 |
| **CODE (1-5)** | SCID | No head trauma, no medical, neurological or psychiatric history, no lifetime alcohol or substance abuse, no previous or current use of psychotropic medication, IQ>75. No history of any psychiatric disorders in 1st degree family members. | 3T Siemens Trio (4 CODE sites); 3T Philips Achieva (1 CODE site) | Siemens: T1 mprage, 1mm isotropic voxels, 12 channel head coil, TR=1900ms, TE=2.52ms, 170/192 slices.  Philips: T1 3D-TFE, 1mm isotropic voxels, 8 channel head coil, TR=8.3ms, TE=3.8ms, 170 slices | 5.3 |
| **ENIGMA-HIV** | MINI | No head trauma, no medical, neurological or psychiatric history, no alcohol or substance abuse preceding 6 months, no previous or current use of psychotropic medication, IQ>75. No mild cognitive impairment | 3T Siemens Allegra | T1-weighted MPRAGE; TR/TE/TI/FA=2400 ms/2.38 ms/1000 ms/ 8°; 162 slices; voxel size= 1 mm^3^ | 5.1 |
| **ENIGMA-OCD (1)** | MINI-Plus | No head trauma, no medical, neurological or psychiatric history, no lifetime alcohol or substance abuse, no previous or current use of psychotropic medication, IQ>75. No cognitive impairment | 3T Siemens Allegra | T1-weighted 3D MPRAGE; TR/TE/TI/FA=2300 ms/3.93 ms/ 1100 ms/12^o^; image matrix =256×240; 160 contiguous sagittal slices; voxel size=1.3 x 1 x 1 mm^3^ | 5.3 |
| **ENIGMA-OCD (2)** | SCID-I | No medical or psychiatric history | 1.5T Siemens Sonata | T1-weighted 3D MPRAGE; TR/TE/TI/FA=2700 ms/4 ms/ 950 ms/8^o^; image matrix =256×192; 160 slices; voxel size= 1 mm^3^ | 5.3 |
| **ENIGMA-OCD (3)** | SCID-I | No medical or psychiatric history | 3T General Electric Signa | T1-weighted 3D MPRAGE; image matrix =256×256; 172 slices; voxel size= 1x0.977x0.977 mm^3^ | 5.3 |
| **ENIGMA-OCD (4)** | Personal Interview | No head trauma, no medical, neurological or psychiatric history, no lifetime alcohol or substance abuse, no previous or current use of psychotropic medication, IQ>75. | 3T Phillips Intera | T1-weighted 3D MPRAGE; TR/TE/ FA=9.69 ms/4.60 ms/8^o^; image matrix =256×256; 182 slices; voxel size=1 x 1 x 1.2 mm^3^ | 5.3 |
| **ENIGMA-OCD (5)** | SCID | No head trauma, no neurological or psychiatric history, no lifetime alcohol or substance abuse, no previous or current use of psychotropic medication, IQ>75. | 1.5T General Electric Signa | T1-weighted 3D SPGR; TR/TE/FA= 14.8 ms/ 1.7 ms/ 20º; image matrix=  256 x 256 x 124; voxel size: 0.94 x 0.94 x 1.50 mm | 5.3 |
| **ENIGMA-OCD (6)** | Personal Interview | No head trauma, no neurological or psychiatric history, no lifetime alcohol or substance abuse. | 3T Phillips Achieva | T1-weighted 3D TFE; TR/TE/TI/FA=8.2 ms/3.8ms/1026 ms/8^o^; image matrix =240×240; 190 slices; voxel size=1 mm^3^ | 5.3 |
| **ENIGMA-OCD (7)** | SCID-I/NP | No head trauma, no medical, neurological or psychiatric history, no alcohol or substance abuse in the preceding 6 months, no previous or current use of psychotropic medication, IQ>70. No history of any psychiatric disorders in 1st or 2^nd^ degree family members | 1.5 T General Electric Signa | T1-weighted 3D FSPGR; TR/TE/ FA=11.8ms/4.2ms/90^o^;  Image matrix= 256 × 256 x 130; voxel size=1.2mm^3^ | 5.3 |
| **FBIRN** | SCID-I/NP | No head trauma, no medical, neurological or psychiatric history, no alcohol or substance abuse in the preceding 5 years, no previous or current use of psychotropic medication, IQ>75. No history of any Axis-I psychotic disorders in 1st degree family members. | 3T Siemens Tim ^Trio^ or General Electric Discovery MR750 | T1-weighted SPGR; TR/TE/TI/FA=2300 ms/2.94 ms/1100 ms/9^o^; image matrix=256×256x160; voxel size=0.86x0.86x1.2mm^3^; sagittal plane acquisition | 5.1 |
| **FIDMAG** | Personal interview; structured interview in part of the sample | No head trauma, no medical, neurological or psychiatric history, no lifetime alcohol or substance abuse, no previous or current use of psychotropic medication, IQ>70. | 1.5 T General Electric Signa | T1-weighted MPRAGE; TR/TE/FA=2000 ms/4 ms/ 9^o^; image matrix=512 x 512; 180 contiguous sagittal slices; voxel size=0.56 x 0.56 x 1 mm3 | 5.3 |
| **GSP** | Structured phone screen and study specific self-report battery and clinical screen | No head trauma, no medical, neurological or psychiatric history, no lifetime alcohol or substance abuse, no current use of psychotropic medication, normal brain anatomy following brain scan. | 3T Siemens Tim Trio | T1-weighted 3D multi-echo MPRAGE; TR/TE/TI/FA =2200 ms/1.54-7 ms/  1100/7 ^o^; voxel size=1.2x1.2x1.2 mm | 4.5 |
| **HMS** | Personal Interview | No head trauma, no medical, neurological or psychiatric history, no lifetime alcohol or substance abuse, no previous or current use of psychotropic medication, IQ>75. No history of any psychiatric disorders in 1st degree family members. | 1.5T Siemens Magnetom Sonata | T1-weighted 3D MPRAGE; TR/TE/TI/FA=1900 ms/4.0 ms/700 ms/15°; image matrix = 256 x 256; 176 consecutive sagittal slices; voxel size=1 mm^3^ | 5.3 |
| **HUBIN** | SCID-I | No head trauma, no medical, neurological or psychiatric history, no lifetime alcohol or substance abuse, no previous or current use of psychotropic medication, IQ>75. No history of any psychiatric disorders in 1st degree family members. | 1.5 T General Electric Signa | T1-weighted SPGR; TR/TE/FA= 24 ms/6 ms/35 ^o^; 124 coronal slices; voxel size 0.86 x 0.86 x 1.50 mm3. | 5.3 |
| **IMH** | SCID-I/NP | No head trauma, no medical history, neurological or psychiatric history, no lifetime alcohol or substance abuse, as well as no previous or current use of psychotropic medication, IQ>75. No cognitive impairment | 3T Phillips Achieva | T1-weighted 3D MPRAGE; TR/TE/FA= 7.2ms/ 3.8ms/8^o^; image matrix=256 x 256; 180 axial slices; voxel size=0.9mm^3^ | 5.3 |
| **Indiana 1.5T** | Personal Interview | No head trauma, no medical, neurological or psychiatric history, no lifetime alcohol or substance abuse, no previous or current use of psychotropic medication, IQ>75. | 1.5T General Electric Signa Horizon LX | T1-weighted 3D SPGR; TR/TE/FA=25 ms/3 ms/ 45^o^; image matrix= 256 x  256; 124 contiguous coronal slices | 5.1 |
| **Indiana 3T** | Personal interview  Structured phone screen | No head trauma, no medical, neurological or psychiatric history, no alcohol or substance abuse in the preceding 6 months, no previous or current use of psychotropic medication, IQ>75. | 3T Siemens Skyra | T1-weighted MPRAGE; TR/TE/FA=2300 ms/2.95 ms/ 9^o^; image matrix=256  x 240; 176 contiguous sagittal slices | 5.1 |
| **KaSP** | MINI | No head trauma, no medical, neurological or psychiatric history, no lifetime alcohol or substance abuse, no previous or current use of psychotropic medication, IQ>75. No history of any psychiatric disorders in 1^st^ or 2^nd^ degree family members. | 3T General Electric | T1-weighted SPGR; TR/TI/FA=7.904 ms/450 ms/12 ^o^; image matrix= 256 x 256 mm^3^; 145 sagittal slices ; voxel size=0.934 x 0.934 x 1.2 mm3 | 5.3 |
| **MAS** | Personal interview | No head trauma, no diagnosis of dementia, schizophrenia, bipolar disorder no psychotic symptoms, no neurological disorder, no mild cognitive impairment, IQ>75. | 3T Philips Achieva Quasar Dual | TR/TE = 6.39 ms/2.9 ms; 190 coronal slices; voxel size = 1mm^3^ | 5.3 |
| **MCIC** | SCID, SCID-I/NP, CASH | No head trauma, no medical, neurological or psychiatric history, no lifetime alcohol or substance abuse, no previous or current use of psychotropic medication, IQ>75. | 1.5T Siemens Sonata-3T Siemens Trio | T1-weighted MPRAGE sequence; TR/TE/TI/FA=2530 ms/4.76 ms/1100  ms/20^o^; image matrix=256×256×128 cm; voxel size=0.625 mm^3^ | 5.3 |
| **Melbourne** | SCID-I | No head trauma, no neurological or psychiatric history, no lifetime alcohol or substance abuse, no previous or current use of psychotropic medication. No history of any psychiatric disorders in 1^st^ or 2^nd^ degree family members. | 3T GE Signa Excite | 3D BRAVO sequence 140; TR/TE/FA=7900 ms/3000 ms/13º; FOV=256 mm; matrix=256 x 256 | 5.3 |
| **Meth-CT** | SCID DSM-IV | No head trauma, no medical, neurological or psychiatric history, no lifetime alcohol or substance abuse, no previous or current use of psychotropic medication, IQ>70. | 3T Siemens Allegra | T1-weighted 3D MPRAGE; TR/graded TE/FA=2530 ms/ 1.53, 3.21, 4.89, 6.57 ms/ 7^o^; 160 contiguous sagittal slices; voxel size=1 x 1 *x 1 mm3 | 5.3 |
| **NESDA** | CIDI | No lifetime history of Axis-I diagnoses, no lifetime medical or neurological morbidity including hypertension, no lifetime substance dependence, no substance abuse in the preceding year, no medication use. | 3T Philips Achieva  SENSE-6 to 8 channel head coil | T1-weighted 3D MPRAGE; TR/TE/FA= 9 ms/3.5 ms/8^o^; image matrix=256x256; 170 sagittal slices; voxel size=1mm^3^ | 5.0 |
| **NeuroIMAGE** | KSADS-PL | No head trauma, no mild cognitive impairment, neurological or psychiatric history, no previous or current use of psychotropic medication, IQ>75. No history of any psychiatric disorders in 1st and 2nd degree family members. | 1.5 T Siemens AVANTO (Donders Centre for Cognitive Neuroimaging)  1.5 T Siemens SONATA (VU University Amsterdam) | MPRAGE 176 sagittal slices, repetition time=2,730ms, echo time=2.95ms, voxel size=1.0x1.0x1.0mm, field of view=256 mm | 5.3 |
| **Neuroventure** | DAWBA and BSI | No head trauma, no medical, neurological or psychiatric history, no lifetime alcohol or substance abuse, no previous or current use of psychotropic medication, IQ>75. | 3T SIEMENS TrioTim | T1-weighted 3D MPRAGE; TR/TE/ FA= 2300 ms/2.96 ms/9^o^; image matrix= 256x256; voxel size= 1.0x1.0x1.0 mm^3^ | 5.3 |
| **NU** | SCID | No head trauma, no medical, neurological or psychiatric history, no lifetime alcohol or substance abuse, no previous or current use of psychotropic medication, IQ>75. No history of any psychiatric disorders in 1st degree family members. | 1.5T SIEMENS Vision | T1-weighted 3D MPRAGE; TR/TE/TI/FA=2200 ms/4.13 ms/766 ms/13°; voxel size =0.8mm^3^; axial plane acquisition. | 5.3 |
| **NUIG** | SCID | No head trauma, no neurological or psychiatric history, no alcohol or substance abuse preceding 6 months, no previous or current use of psychotropic medication, IQ>75. No history of any psychiatric disorders in 1st degree family members. | Siemens Magnetom Symphony 1.5T | 3D, T1-weighted MPRAGE 4 channel head coil, FOV 230mm, TR/TE/: 1140ms/4.38ms, matrix size 256 x 256, interpolated to 512 x 512, yielding an in-plane voxel size of 0.45mm x 0.45mm^2^, slice thickness 0.9mm. | 5.1 |
| **NYU** | SCID-NP for DSM-IV | No head trauma, no medical history, including neurological and psychiatric history, no lifetime alcohol or substance abuse, no previous or current use of psychotropic medication. IQ>75. | 3T Siemens Allegra | T1-weighted 3D MPRAGE; TR/TE/TI/FA=2530ms/3.25ms/1100ms/7^o^ | 5.3 |
| **Olin** | SCID I | No head trauma, no medical, neurological or psychiatric history, no alcohol or substance abuse preceding 6 months, never prescribed with psychotropic medication, IQ>75. | 3T Siemens Alegra | T1-weighted 3D MPRAGE; TR/TE/TI/FA= 2300 ms/2.91 ms/900 ms/9^o^; image matrix= 256x240x192; 160 sagittal slices; voxel size= 1.0x1.0x1.2 mm^3^ | 5.1 |
| **Oxford** | KSADS | No head trauma, no medical, neurological or psychiatric history, no lifetime alcohol or substance abuse, no previous or current use of psychotropic medication, IQ>75. | 1.5T Siemens Sonata | T1-weighted 3D MPRAGE; TR/TE =12 ms/5.6 ms; image matrix =256×240x 208 mm^3^; voxel size=1 mm^3^ | 5.3 |
| **QTIM** | CIDI | No head trauma, no medical history, neurological and psychiatric history, no alcohol or substance abuse in the preceding 6 months, no antidepressant medication or medication affecting cognition. | 4T Bruckner | T1-weighted 3D MPRAGE: TR/TE/TI/FA = 1500 ms/3.35 ms/ 700 ms/ 8°; image matrix= 256 × 256 × 256 or 256 × 256 × 240; 256 coronal slices; voxel size= 0.9 mm^3^ | 5.1 |
| **Sao Paulo (3)** | SCID | No head trauma, neurological or psychiatric history, no lifetime alcohol or substance abuse. IQ>75 | 1.5T General Electric Signa | T1-weighted FSPGR; TR/TE/TI/FA=21.7 ms/52 ms /20^o^; 124 axial slices; voxel size= 0.86 x 0.86 x 1.5 mm3 | 5.3 |
| **SCORE** | BPRS | No head trauma, no medical, neurological or psychiatric history, no lifetime history of alcohol or substance abuse, no previous or current use of psychotropic medication, IQ>75. No history of any psychiatric disorders in 1st degree family members. | 3T Siemens Magnetom Verio | T1-weighted 3D-MPRAGE; TR/TE/TI/FA =2000 ms/3.37 ms/1000 ms/8^o^; image matrix=256x256x176; 176 consecutive sagittal slices; voxel size= 1 mm^3^ | 6.0 |
| **SHIP-2** | Personal Interview | No head trauma, no neurological and psychiatric history, no alcohol or substance abuse in the preceding 6 months, no previous or current use of psychotropic medication. IQ>75. | 1.5T Siemens Avanto | T1-weighted 3D MPRAGE; TR/TE/ FA=1900ms/3.4ms/15^o^; voxel size=1mm^3^ | 5.3 |
| **SHIP-TREND** | Personal Interview | No head trauma, no neurological and psychiatric history, no alcohol or substance abuse in the preceding 6 months, no previous or current use of psychotropic medication. IQ>75. | 1.5T Siemens Avanto | T1-weighted 3D MPRAGE; TR/TE/ FA=1900ms/3.4ms/15^o^; voxel size=1 mm^3^ | 5.3 |
| **Staged-Dep** | SCID-I | No head trauma, no medical, neurological or psychiatric history, no lifetime alcohol or substance abuse, no previous or current use of psychotropic medication, IQ>75. No history of any psychiatric disorders in 1st degree family members. | 3T Phillips Achieva | T1-weighted 3D-MPRAGE; TR/TE/TI/FA =6.7 ms/3.2 ms/200 ms/88^o^; °; image matrix = 288 x 288; 170 consecutive sagittal slices; voxel size= 0.896×0.896×1.2 mm^3^ | 5.1 |
| **Stanford** | SCID | No head trauma, no medical, neurological or psychiatric history, no lifetime alcohol or substance abuse, no previous or current use of psychotropic medication. no mild cognitive impairment. | 1.5T General Electric Signa Excite | T1-weighted SPGR; TR/TE/TI/FA=8.3-10.3 ms/1.7-3.0 ms/300 ms/15^o^; image matrix= 256 x 192; 176 contiguous sagittal slices; voxel size=0.86x0.86x1.5 mm^3^; sagittal plan acquisition | 5.3 |
| **StrokeMRI** | Personal interview | No head trauma, no medical, neurological or psychiatric history, no lifetime alcohol or substance abuse, no previous or current use of psychotropic medication, IQ>75. | 3T General Electric Signa HDxt | T1-weighted FSPGR; TR/TE/TI/FA=7.8 s/2.956 ms/450 ms/12°; 170 slices; voxel size= 1.0x1.0x1.2 mm | 5.3 |
| **Sydney** | SCID | No head trauma, no medical history, neurological or psychiatric history, no alcohol or substance abuse preceding 6 months, as well as no previous or current use of psychotropic medication, IQ>75. | 3T General Electric Discovery MR750 | T1-weighted 3D MPRAGE; TR/TE/FA= 7264ms/ 2784ms/15^o^; image matrix  =256 x 256 x 196; voxel size=0.9mm^3^ | 5.1 |
| **TOP** | PRIME-MD | No head trauma, no organic or other psychotic disorder (ICD codes 290-299), no substance abuse in the preceding 6 months, no previous or current use of psychotropic medication, IQ>75. No history of any psychiatric disorders in 1st degree family members. | 1.5T Siemens Magnetom Sonata | T1-weighted SPGR; TR/TE/TI/FA=2730 ms/3.93 ms/1000 ms/71^o^; voxel size = 1.33x0.94x1mm^3^; sagittal plane acquisition | 5.3 |
| **TS-EUROTRAIN** | KSADS | No head trauma, no medical, neurological or psychiatric history, no alcohol or substance abuse preceding 6 months, no previous or current use of psychotropic medication, IQ>75. No history of any psychiatric disorders in 1^st^ or 2^nd^ degree family members. | 3T Siemens Tim Trio and Prisma | T1-weighted 3D MPRAGE; TR/TE/FA=2300 ms/2.98 ms/9°; image matrix = 256 x 256; 176 sagittal slices; voxel size= 1x1x1.2 mm^3^ | 5.3 |
| **Tuebingen** | SCID I and II | No head trauma, no medical history, neurological or psychiatric history, no lifetime alcohol or substance abuse as well as no previous or current use of psychotropic medication, IQ>75. No history of any psychiatric disorders in 1st degree family members. | 1.5T Siemens Avanto | T1-weighted 3D MPRAGE; TR/TE/FA= 2250ms/ 3.93ms/8^o^; image matrix  =256 x 256; voxel size=1mm^3^ | 5.3 |
| **UMCU** | CASH | No head trauma, no medical, neurological or psychiatric history, no lifetime alcohol or substance abuse, no previous or current use of psychotropic medication, IQ>75. No history of any psychiatric disorders in 1st degree family members. | 1.5T Philips Intera and Achieva | T1-weighted 3D FFE; TE/TR/FA= 4.6 ms/0 ms**/ 0˚;** 160-180 contiguous coronal slices; voxel size=1x1x1.2 mm^3^ | 5.1 |
| **UNIBA** | SCID-NP | No head trauma, no medical, neurological or psychiatric history, no lifetime alcohol or substance abuse, no previous or current use of psychotropic medication, IQ>75. No history of any psychiatric disorders in 1st degree family members. | 3T General Electric | T1-weighted 3D SPGR; TE/FA = min full/ 6°; image matrix= 256×256 x124 | 5.3 |
| **UPENN** | SCID | No head trauma, no medical history, including neurological and psychiatric history, no alcohol or substance abuse preceding 6 months, no previous or current use of psychotropic medication, IQ>75. No history of any psychiatric disorders in 1st degree family members. | 3T Siemens Tim Trio | T1-weighted 3D MPRAGE; TR/TE/TI/FA=1810 ms/3.51 ms/1100 ms/9^o^; image matrix= 256 × 192;160 axial slices | 5.3 |
| **Yale** | KSADS-PL | No head trauma, neurological or psychiatric history, no alcohol or substance abuse in the preceding 6 months, no previous or current use of psychotropic medication, IQ>75. | 3T General Electric Signa | T1-weighted 3D MPRAGE; image matrix =256×256; voxel size=0.976 x 0.976 x 1 mm3 | 5.3 |
| **Abbreviations of Terms**: BDI = Behavioural Descriptive Interview; BSI = Brief Symptom Inventory; CASH = Comprehensive assessment of symptoms and history; CDR = Clinical Dementia Rating; CIDI = Composite International Diagnostic Interview; DAWBA = Development and Well-Being Assessment; DISC-IV = Diagnostic Interview Schedule for Children; DSM = Diagnostic and Statistical Manual of Mental Disorders (DSM); FA=flip angle; FSPGR=fast spoiled gradient echo sequence; GRE=spoiled gradient echo sequence; ICD= International Classification of Diseases; IR= inversion recovery; KSADS-PL= Kiddie Schedule for Affective Disorders and Schizophrenia-Present and Lifetime; MADRS = Montgomery-Asberg Depression Rating Scale; MINI = Mini International Neuropsychiatric Interview; MMSE = Mini Mental State Exam; PRIME-MD = Primary Care Evaluation of Mental Disorder; SCID = Structured Clinical Interview for DSM Disorders; SCID-I/NP = SCID Non-Patient version; SPGR=spoiled gradient recalled sequence; STAI = State-Trait Anxiety Inventory; STAS = State-trait anger scale; TE=echo time; TI=inversion time; TR=repetition time; TFE=turbo field echo sequence; YBOCS = Yale-Brown Obsessive Compulsive Scale  **Abbreviations of samples:** ADHD-NF = Attention Deficit Hyperactivity Disorder- Neurofeedback Study; AMC = Amsterdam Medisch Centrum; Barcelona = University of Barcelona; Betula = Swedish longitudinal study on aging, memory, and dementia; BIG = Brain Imaging Genetics; BIL&GIN = a multimodal multidimensional database for investigating hemispheric specialization; CAMH = Centre for Addiction and Mental Health; Cardiff = Cardiff University; CEG = Cognitive-experimental and Genetic study of ADHD and Control Sibling Pairs; CIAM = Cortical Inhibition and Attentional Modulation study; CLiNG = Clinical Neuroscience Göttingen; CODE = formerly Cognitive Behavioral Analysis System of Psychotherapy (CBASP) study; ENIGMA-HIV = Enhancing NeuroImaging Genetics through Meta-Analysis-Human Immunodeficiency Virus Working Group; ENIGMA-OCD = Enhancing NeuroImaging Genetics through Meta-Analysis- Obsessive Compulsive Disorder Working Group; FBIRN = Function Biomedical Informatics Research Network; FIDMAG = Fundación para la Investigación y Docencia Maria Angustias Giménez; GSP = Brain Genomics Superstruct Project; HMS = Homburg Multidiagnosis Study; HUBIN = Human Brain Informatics; IMH=Institute of Mental Health, Singapore; Indiana = Indiana University School of Medicine; KaSP= The Karolinska Schizophrenia Project; MAS = Memory and Ageing Study; MCIC = MIND Clinical Imaging Consortium formed by the Mental Illness and Neuroscience Discovery (MIND) Institute now the Mind Research Network; Melbourne = University of Melbourne; Meth-CT = methamphetamine use, University of Cape Town; NESDA = The Netherlands Study of Depression and Anxiety; NeuroIMAGE = Dutch part of the International Multicenter ADHD Genetics (IMAGE) study; Neuroventure: the imaging part of the Co-Venture Trial funded by the Canadian Institutes of Health Research (CIHR); NU = Northwestern University; NUIG = National University of Ireland Galway; NYU = New York University; Olin = Olin Neuropsychiatric Research Center; Oxford =Oxford University; QTIM = Queensland Twin Imaging; Sao Paulo = University of Sao Paulo; SCORE: University of Basel Study; SHIP-2 and SHIP TREND = Study of Health in Pomerania; Staged-Dep= Stages of Depression Study; Stanford = Stanford University; StrokeMRI = Stroke Magnetic Resonance Imaging; Sydney = University of Sydney; TOP = Tematisk Område Psykoser (Thematically Organized Psychosis Research); TS-EUROTRAIN = European-Wide Investigation and Training Network on the Etiology and Pathophysiology of Gilles de la Tourette Syndrome; Tuebingen = University of Tuebingen; UMCU = Universitair Medisch Centrum Utrecht; UNIBA = University of Bari Aldo Moro; UPENN=University of Pennsylvania; Yale = Yale University | | | | | |

| **Table S2. Age at Maximum Fitted Value for Each Subcortical Volume** | | |
| --- | --- | --- |
| **Region** | **Age (years) at**  **maximum volume**  **Males Only** | **Age (years) at**  **maximum volume**  **Females Only** |
| **Left Lateral Ventricle** | 90 | 90 |
| **Right Lateral Ventricle** | 90 | 90 |
| **Left Thalamus** | 25 | 22 |
| **Right Thalamus** | 16 | 7 |
| **Left Amygdala** | 30 | 7 |
| **Right Amygdala** | 30 | 18 |
| **Left Hippocampus** | 14 | 23 |
| **Right Hippocampus** | 24 | 27 |
| **Left Caudate** | 6 | 7 |
| **Right Caudate** | 6 | 7 |
| **Left Putamen** | 6 | 7 |
| **Right Putamen** | 6 | 7 |
| **Left Nucleus Accumbens** | 6 | 7 |
| **Right Nucleus Accumbens** | 6 | 7 |
| **Left Globus Pallidus** | 6 | 7 |
| **Right Globus Pallidus** | 6 | 7 |

| **Table S3. Variance Explained by Age in Fractional Polynomial Model** | |
| --- | --- |
| **Region** | **R-Squared** |
| **Left Lateral Ventricle** | 0.43 |
| **Right Lateral Ventricle** | 0.43 |
| **Left Thalamus** | 0.26 |
| **Right Thalamus** | 0.29 |
| **Left Caudate** | 0.14 |
| **Right Caudate** | 0.14 |
| **Left Putamen** | 0.35 |
| **Right Putamen** | 0.38 |
| **Left Globus Pallidus** | 0.20 |
| **Right Globus Pallidus** | 0.20 |
| **Left Hippocampus** | 0.22 |
| **Right Hippocampus** | 0.20 |
| **Left Amygdala** | 0.15 |
| **Right Amygdala** | 0.09 |
| **Left Nucleus Accumbens** | 0.23 |
| **Right Nucleus Accumbens** | 0.28 |

| **Table S4. Pearson's Correlation Coefficient Between Age and Subcortical Volumes** | | | | | | | | | |
| --- | --- | --- | --- | --- | --- | --- | --- | --- | --- |
|  | **All Participants** | | | **Males Only** | | | **Females Only** | | |
| **Region** | **6-29 years** | **30-59 years** | **60-90 years** | **6-29 years** | **30-59 years** | **60-90 years** | **6-29 years** | **30-59 years** | **60-90 years** |
| **Left Lateral Ventricle** | 0.099 | 0.256 | 0.498 | 0.106 | 0.275 | 0.494 | 0.092 | 0.237 | 0.525 |
| **Right Lateral Ventricle** | 0.097 | 0.263 | 0.509 | 0.1 | 0.29 | 0.506 | 0.094 | 0.232 | 0.535 |
| **Left Thalamus** | -0.002 | -0.292 | -0.275 | -0.006 | -0.297 | -0.222 | 0.001 | -0.284 | -0.327 |
| **Right Thalamus** | -0.048 | -0.333 | -0.311 | -0.063 | -0.328 | -0.279 | -0.035 | -0.336 | -0.347 |
| **Left Caudate** | -0.161 | -0.224 | -0.041 | -0.171 | -0.237 | -0.05 | -0.152 | -0.21 | -0.034 |
| **Right Caudate** | -0.178 | -0.225 | 0 | -0.194 | -0.249 | -0.018 | -0.164 | -0.197 | 0.017 |
| **Left Putamen** | -0.225 | -0.381 | -0.203 | -0.252 | -0.422 | -0.243 | -0.206 | -0.33 | -0.167 |
| **Right Putamen** | -0.253 | -0.404 | -0.182 | -0.266 | -0.442 | -0.215 | -0.248 | -0.356 | -0.151 |
| **Left Globus Pallidus** | -0.225 | -0.261 | -0.194 | -0.25 | -0.272 | -0.209 | -0.212 | -0.243 | -0.18 |
| **Right Globus Pallidus** | -0.185 | -0.234 | -0.268 | -0.213 | -0.209 | -0.265 | -0.164 | -0.262 | -0.271 |
| **Left Hippocampus** | -0.033 | -0.119 | -0.48 | -0.038 | -0.119 | -0.471 | -0.029 | -0.117 | -0.494 |
| **Right Hippocampus** | -0.029 | -0.1 | -0.472 | -0.04 | -0.114 | -0.478 | -0.02 | -0.082 | -0.472 |
| **Left Amygdala** | -0.017 | -0.11 | -0.379 | 0.005 | -0.128 | -0.356 | -0.038 | -0.08 | -0.398 |
| **Right Amygdala** | -0.016 | -0.086 | -0.288 | -0.008 | -0.105 | -0.253 | -0.025 | -0.054 | -0.32 |
| **Left Nucleus Accumbens** | -0.152 | -0.275 | -0.29 | -0.158 | -0.276 | -0.263 | -0.147 | -0.274 | -0.317 |
| **Right Nucleus Accumbens** | -0.19 | -0.295 | -0.254 | -0.179 | -0.318 | -0.203 | -0.2 | -0.269 | -0.304 |

| **Table S5. Inter-individual Variation in Subcortical Volumes** | | | | | | | |
| --- | --- | --- | --- | --- | --- | --- | --- |
| **Region** | **Inter-individual variation**  **Mean (standard deviation)** | | |  | **P value for sex differences** | | |
|  | **6-29**  **years** | **30-59**  **years** | **60-90**  **years** | **Unadjusted P value for F test** | **6-29 years** | **30-59 years** | **60-90 years** |
| **Left Lateral Ventricle** | 2655 (2686) | 3053 (2791) | 4262 (3922) | **0** | 0 | 0 | 0.57 |
| **Right Lateral Ventricle** | 2456 (2456) | 2835 (2721) | 3982 (3564) | **0** | 0 | 0 | 0.33 |
| **Left Thalamus** | 490 (384) | 537 (422) | 538 (464) | **0.001** | 0 | 0.59 | 0.01 |
| **Right Thalamus** | 441 (353) | 469 (375) | 470 (435) | 0.46 | 0 | **0.0003** | **0.002** |
| **Left Caudate** | 329 (251) | 317 (248) | 330 (299) | 0.26 | **0.001** | 0.87 | 0.003 |
| **Right Caudate** | 335 (258) | 318 (252) | 348 (305) | 0.06 | 0.09 | 0.18 | 0.15 |
| **Left Putamen** | 453 (347) | 441 (349) | 465 (378) | 0.17 | 0.01 | 0.38 | 0.18 |
| **Right Putamen** | 416 (323) | 416 (329) | 438 (369) | 0.70 | 0 | 0.06 | 0.47 |
| **Left Globus Pallidus** | 164 (132) | 166 (131) | 170 (136) | 0.23 | **0.0001** | **0.0002** | 0.02 |
| **Right Globus Pallidus** | 132 (109) | 136 (112) | 136 (114) | 0.34 | 0 | 0.02 | 0.01 |
| **Left Hippocampus** | 274 (224) | 296 (233) | 327 (268) | **0** | 0 | 0.01 | 0.29 |
| **Right Hippocampus** | 267 (213) | 290 (225) | 316 (272) | **0** | **0.0003** | 0.03 | **0.002** |
| **Left Amygdala** | 139 (108) | 147 (113) | 161 (133) | **0.0003** | 0.09 | 0.21 | 0.06 |
| **Right Amygdala** | 144 (112) | 151 (116) | 165 (135) | **0** | 0 | 0.13 | 0.01 |
| **Left Nucleus Accumbens** | 81 (62) | 79 (62) | 87 (67) | 0.06 | **0.002** | 0.13 | 0.31 |
| **Right Nucleus Accumbens** | 74 (57) | 76 (59) | 80 (62) | 0.01 | 0 | 0.10 | 0.005 |

| **Table S6. Centile Values for Subcortical Volumes – All participants** | | | | | | | | | | |
| --- | --- | --- | --- | --- | --- | --- | --- | --- | --- | --- |
| **Region** | **Age** | **C0.4** | **C2** | **C10** | **C25** | **C50** | **C75** | **C90** | **C98** | **C99.6** |
| **Left Lateral Ventricle** | 6 | 461.8519555 | 894.7297485 | 1748.846917 | 2713.851644 | 4196.186199 | 6299.316444 | 9004.812873 | 14321.6895 | 20981.14186 |
|  | 10 | 561.0399805 | 1050.324586 | 1978.673552 | 2996.510817 | 4528.639848 | 6675.224504 | 9427.54924 | 14876.44619 | 21828.67015 |
|  | 15 | 715.4569463 | 1286.0452 | 2315.923872 | 3401.829901 | 4993.420511 | 7185.901198 | 9983.318919 | 15571.07122 | 22871.34034 |
|  | 20 | 888.9811434 | 1553.142386 | 2693.295259 | 3846.94594 | 5491.542958 | 7721.674292 | 10565.22146 | 16350.43842 | 24205.64415 |
|  | 30 | 1215.960024 | 2019.670411 | 3315.683683 | 4566.08221 | 6299.199965 | 8626.091416 | 11627.8828 | 18002.44114 | 27344.38312 |
|  | 40 | 1869.557414 | 2676.35326 | 3999.164773 | 5324.589078 | 7214.595761 | 9766.800728 | 12971.91369 | 19266.30836 | 27361.5774 |
|  | 50 | 2599.426811 | 3450.042558 | 4869.016995 | 6321.753795 | 8421.330955 | 11252.71481 | 14731.1009 | 21190.8785 | 28777.9865 |
|  | 60 | 3449.187985 | 4537.874475 | 6297.124079 | 8044.572191 | 10505.29721 | 13746.15838 | 17655.78692 | 24813.48612 | 33157.23626 |
|  | 70 | 4751.263108 | 6590.900139 | 9184.469438 | 11450.15491 | 14358.16912 | 17980.58459 | 22329.80348 | 30792.95134 | 42051.50959 |
|  | 80 | 7044.493425 | 10147.75941 | 13786.3353 | 16466.68403 | 19530.11093 | 23101.03015 | 27362.52749 | 36184.29895 | 49514.24824 |
|  | 90 | 12153.54983 | 15349.39789 | 18936.89909 | 21522.53713 | 24401.65401 | 27597.12142 | 31132.00635 | 37561.90865 | 45660.63769 |
| **Right Lateral Ventricle** | 6 | 380.6197808 | 805.5153755 | 1625.525122 | 2517.118742 | 3845.271719 | 5701.417357 | 8107.60877 | 13028.3138 | 19651.85246 |
|  | 10 | 474.1462813 | 951.069876 | 1839.135925 | 2783.129691 | 4169.824593 | 6090.051536 | 8568.746614 | 13639.77406 | 20504.13844 |
|  | 15 | 618.8601629 | 1166.728518 | 2144.184888 | 3154.636947 | 4612.813582 | 6607.227291 | 9164.678772 | 14388.69522 | 21491.73496 |
|  | 20 | 786.8394028 | 1404.542059 | 2465.99618 | 3536.609385 | 5057.834092 | 7116.877516 | 9743.972126 | 15111.33364 | 22456.03872 |
|  | 30 | 1179.613725 | 1896.214011 | 3067.482844 | 4217.165621 | 5827.179853 | 7986.688869 | 10729.28423 | 16328.85332 | 24029.03632 |
|  | 40 | 1711.545223 | 2492.125266 | 3749.379342 | 4985.992793 | 6725.376585 | 9056.759659 | 11989.08167 | 17830.38971 | 25556.79432 |
|  | 50 | 2260.744832 | 3127.19154 | 4520.004582 | 5897.299612 | 7845.870061 | 10465.35871 | 13752.36112 | 20232.45603 | 28646.87116 |
|  | 60 | 2953.891102 | 4047.833663 | 5758.404864 | 7407.735288 | 9694.294009 | 12716.62315 | 16466.61884 | 23813.85707 | 33362.62971 |
|  | 70 | 4394.015349 | 6051.171844 | 8403.95084 | 10474.21693 | 13145.00238 | 16480.72118 | 20483.37651 | 28235.19702 | 38455.58347 |
|  | 80 | 6832.900078 | 9478.857911 | 12713.54551 | 15199.05329 | 18110.49757 | 21518.23425 | 25505.37666 | 33336.53503 | 44217.72505 |
|  | 90 | 9234.513271 | 13086.09097 | 17184.8523 | 19968.61653 | 22979.77037 | 26357.1005 | 30304.64271 | 38386.7179 | 50502.38467 |
| **Left**  **Thalamus** | 6 | 6448.50376 | 6728.597024 | 7102.338261 | 7408.470428 | 7764.611095 | 8141.513452 | 8502.459662 | 8998.258009 | 9416.920071 |
|  | 10 | 6428.189665 | 6719.430468 | 7105.499815 | 7420.452633 | 7786.454956 | 8174.46643 | 8547.745517 | 9064.82136 | 9506.650013 |
|  | 15 | 6384.88139 | 6692.588764 | 7095.668131 | 7421.997394 | 7800.257818 | 8202.177071 | 8591.609354 | 9138.543188 | 9615.165506 |
|  | 20 | 6316.372477 | 6643.596263 | 7065.455404 | 7403.403019 | 7793.607483 | 8209.203645 | 8615.449725 | 9196.06732 | 9714.981165 |
|  | 30 | 6198.417059 | 6560.338179 | 7017.002855 | 7377.715911 | 7792.269814 | 8235.783487 | 8675.359671 | 9320.649385 | 9919.834371 |
|  | 40 | 6036.882576 | 6409.032118 | 6878.970882 | 7250.659333 | 7678.662342 | 8137.92618 | 8594.859242 | 9269.331239 | 9899.936234 |
|  | 50 | 5773.215044 | 6145.384764 | 6614.435276 | 6984.92209 | 7411.331138 | 7869.049282 | 8325.012432 | 8999.705223 | 9632.721959 |
|  | 60 | 5459.266532 | 5827.288739 | 6287.203852 | 6647.946239 | 7061.161704 | 7503.306441 | 7943.204662 | 8594.525817 | 9207.052492 |
|  | 70 | 4987.658147 | 5433.320063 | 5937.861947 | 6305.44238 | 6711.13182 | 7143.563793 | 7587.728463 | 8298.024494 | 9048.615606 |
|  | 80 | 4139.749705 | 4910.263826 | 5601.108456 | 6016.459718 | 6432.801279 | 6875.808113 | 7378.176245 | 8385.295707 | 9864.9464 |
|  | 90 | 4029.099738 | 4785.753696 | 5467.660053 | 5879.856622 | 6293.510175 | 6731.461339 | 7222.25562 | 8183.943707 | 9549.455184 |
| **Right Thalamus** | 6 | 6455.230736 | 6705.233687 | 7044.685462 | 7326.370743 | 7656.536558 | 8006.982756 | 8341.878579 | 8798.13088 | 9178.035797 |
|  | 10 | 6357.461758 | 6613.823429 | 6959.33071 | 7244.552735 | 7578.020595 | 7931.910054 | 8270.883624 | 8735.244438 | 9125.187659 |
|  | 15 | 6221.422491 | 6492.503744 | 6850.439762 | 7141.725059 | 7479.975376 | 7838.887917 | 8185.048639 | 8666.857674 | 9081.355456 |
|  | 20 | 6071.786172 | 6376.343104 | 6760.28997 | 7062.72407 | 7408.572292 | 7775.575079 | 8135.376377 | 8655.202232 | 9128.281464 |
|  | 30 | 5768.312664 | 6200.822319 | 6676.811247 | 7016.223675 | 7386.388246 | 7779.361263 | 8184.787245 | 8842.560478 | 9554.04948 |
|  | 40 | 5727.801884 | 6106.505345 | 6552.516721 | 6886.796719 | 7259.759802 | 7654.665836 | 8050.295064 | 8652.218087 | 9244.566369 |
|  | 50 | 5510.825843 | 5866.654705 | 6295.527548 | 6622.150855 | 6988.868539 | 7376.001299 | 7758.99258 | 8327.024043 | 8865.838731 |
|  | 60 | 5219.433621 | 5587.143591 | 6018.162839 | 6339.576415 | 6696.304697 | 7071.594748 | 7444.829002 | 8007.078906 | 8553.633807 |
|  | 70 | 4721.172061 | 5215.991092 | 5716.263885 | 6049.762826 | 6400.068793 | 6767.507265 | 7152.353256 | 7805.816229 | 8562.859893 |
|  | 80 | 3925.856321 | 4741.227943 | 5401.318189 | 5765.740538 | 6113.832697 | 6478.000785 | 6897.146643 | 7776.210641 | 9146.980588 |
|  | 90 | 4325.592123 | 4798.356999 | 5264.653294 | 5569.407249 | 5885.309856 | 6213.522422 | 6555.355611 | 7133.537414 | 7800.683431 |
| **Left**  **Caudate** | 6 | 3062.444473 | 3342.828942 | 3694.127676 | 3965.07933 | 4263.973662 | 4563.805221 | 4837.243022 | 5194.139011 | 5480.980941 |
|  | 10 | 2997.169966 | 3263.567984 | 3599.385243 | 3860.151482 | 4149.86898 | 4443.002387 | 4712.838679 | 5069.119901 | 5359.194221 |
|  | 15 | 2919.034278 | 3170.317462 | 3488.905043 | 3738.064143 | 4017.182666 | 4302.602463 | 4568.527111 | 4925.116052 | 5220.647832 |
|  | 20 | 2854.898809 | 3094.154237 | 3398.636485 | 3638.141205 | 3908.496619 | 4187.897242 | 4451.541035 | 4811.116338 | 5115.148562 |
|  | 30 | 2770.687985 | 2995.501105 | 3281.623733 | 3508.012483 | 3766.651839 | 4039.354285 | 4303.628052 | 4678.107622 | 5010.061483 |
|  | 40 | 2680.122907 | 2902.222648 | 3180.394748 | 3398.761697 | 3648.77063 | 3915.814885 | 4180.726124 | 4571.053205 | 4935.655973 |
|  | 50 | 2577.615944 | 2805.184314 | 3081.835971 | 3294.88237 | 3537.658513 | 3799.63932 | 4066.22448 | 4478.067108 | 4889.323964 |
|  | 60 | 2483.776283 | 2737.571202 | 3029.90839 | 3246.110804 | 3487.75339 | 3748.680147 | 4020.144465 | 4461.230613 | 4936.259811 |
|  | 70 | 2345.795836 | 2669.038423 | 3007.190025 | 3238.551907 | 3485.697271 | 3747.946726 | 4024.279877 | 4494.582262 | 5039.395316 |
|  | 80 | 2050.714375 | 2534.605648 | 2965.381031 | 3224.142669 | 3480.416757 | 3743.578575 | 4024.102995 | 4528.300286 | 5161.296223 |
|  | 90 | 1501.637947 | 2300.196817 | 2906.300091 | 3209.133028 | 3478.591044 | 3744.184799 | 4034.342107 | 4600.075546 | 5386.093034 |
| **Right**  **Caudate** | 6 | 3131.587877 | 3418.211551 | 3778.988004 | 4057.790803 | 4365.049222 | 4672.069802 | 4950.243831 | 5309.615651 | 5594.636684 |
|  | 10 | 3061.66463 | 3330.597454 | 3672.610455 | 3939.67137 | 4236.930016 | 4537.244385 | 4812.384752 | 5172.450474 | 5461.981522 |
|  | 15 | 2976.728978 | 3226.616569 | 3547.775375 | 3801.431696 | 4087.086372 | 4379.631398 | 4651.527515 | 5013.548117 | 5310.224845 |
|  | 20 | 2905.139046 | 3140.192771 | 3444.586917 | 3687.232677 | 3963.34659 | 4249.826963 | 4519.978984 | 4886.32596 | 5192.859308 |
|  | 30 | 2815.278087 | 3032.351181 | 3315.291696 | 3543.504768 | 3807.579378 | 4088.254479 | 4360.885063 | 4745.81455 | 5083.85433 |
|  | 40 | 2715.415653 | 2928.531077 | 3202.886393 | 3423.070444 | 3678.817579 | 3954.327937 | 4228.066051 | 4629.050543 | 4998.833137 |
|  | 50 | 2610.168211 | 2831.69561 | 3108.345003 | 3325.846638 | 3576.611652 | 3848.257474 | 4123.348091 | 4541.851382 | 4949.477429 |
|  | 60 | 2513.738114 | 2765.97143 | 3063.72473 | 3287.926979 | 3540.463355 | 3812.539348 | 4091.972646 | 4533.713112 | 4990.565684 |
|  | 70 | 2362.785988 | 2691.005057 | 3041.211522 | 3284.3071 | 3545.039586 | 3819.658157 | 4103.46203 | 4569.048518 | 5081.176256 |
|  | 80 | 2049.519052 | 2560.378915 | 3018.856583 | 3296.379081 | 3571.217608 | 3850.308876 | 4141.087298 | 4643.041771 | 5239.989229 |
|  | 90 | 1475.181351 | 2329.057819 | 2996.971947 | 3331.434871 | 3627.366797 | 3914.852646 | 4221.558111 | 4796.842482 | 5557.709816 |
| **Left**  **Putamen** | 6 | 5204.98834 | 5593.482684 | 6085.145885 | 6467.154866 | 6890.300998 | 7315.467299 | 7702.81297 | 8206.376223 | 8608.408903 |
|  | 10 | 4975.395582 | 5352.378859 | 5828.528241 | 6198.109864 | 6607.515092 | 7019.326286 | 7395.271952 | 7885.654317 | 8278.898791 |
|  | 15 | 4701.977618 | 5065.415789 | 5523.051122 | 5877.672744 | 6270.468381 | 6666.125269 | 7028.329408 | 7502.99593 | 7886.012393 |
|  | 20 | 4491.320954 | 4845.859851 | 5290.583247 | 5634.447957 | 6015.205265 | 6399.283443 | 6751.968163 | 7216.603184 | 7594.198396 |
|  | 30 | 4316.050793 | 4672.201986 | 5114.268197 | 5453.911629 | 5829.413106 | 6209.29209 | 6560.628081 | 7029.437549 | 7417.1379 |
|  | 40 | 4113.447658 | 4471.700719 | 4909.545247 | 5242.653571 | 5609.779947 | 5982.29866 | 6329.924352 | 6801.581362 | 7200.768273 |
|  | 50 | 3855.017564 | 4216.438019 | 4648.498104 | 4972.457464 | 5327.62257 | 5689.137628 | 6030.381235 | 6503.728615 | 6916.959238 |
|  | 60 | 3610.13325 | 3991.057946 | 4432.752419 | 4757.165084 | 5110.006944 | 5470.459321 | 5815.906935 | 6309.651782 | 6759.235295 |
|  | 70 | 3302.880697 | 3725.934885 | 4196.929769 | 4533.17877 | 4894.802627 | 5265.971692 | 5629.173267 | 6170.234685 | 6692.357483 |
|  | 80 | 2960.492197 | 3460.619305 | 3988.5641 | 4351.051806 | 4734.699278 | 5130.879947 | 5529.834825 | 6159.380872 | 6817.745852 |
|  | 90 | 2583.078064 | 3213.896979 | 3836.520611 | 4241.650715 | 4660.561326 | 5096.428609 | 5553.033632 | 6333.850318 | 7246.797333 |
| **Right**  **Putamen** | 6 | 5201.685061 | 5506.303304 | 5900.024479 | 6214.460525 | 6574.263525 | 6951.288327 | 7311.627036 | 7809.814628 | 8236.769641 |
|  | 10 | 4962.07778 | 5265.791776 | 5656.23927 | 5966.774468 | 6321.2289 | 6692.210015 | 7046.877263 | 7538.19172 | 7960.747163 |
|  | 15 | 4674.61585 | 4977.554615 | 5364.102846 | 5669.775395 | 6017.477985 | 6380.805608 | 6728.311541 | 7211.047388 | 7628.283026 |
|  | 20 | 4438.917145 | 4743.476563 | 5128.7904 | 5431.494097 | 5774.464696 | 6132.181765 | 6474.483761 | 6951.467296 | 7366.012039 |
|  | 30 | 4191.467988 | 4510.777566 | 4906.310897 | 5212.029976 | 5555.034382 | 5911.090797 | 6252.110509 | 6730.72332 | 7152.072224 |
|  | 40 | 3932.242521 | 4265.65662 | 4667.785101 | 4972.329178 | 5309.926219 | 5658.453331 | 5992.833033 | 6466.6868 | 6890.890263 |
|  | 50 | 3667.788925 | 4018.159961 | 4426.928691 | 4728.823946 | 5058.704842 | 5397.25848 | 5723.109266 | 6191.061617 | 6619.305819 |
|  | 60 | 3405.390672 | 3787.00591 | 4214.445007 | 4520.654222 | 4849.668901 | 5185.342392 | 5510.225328 | 5985.448345 | 6433.152777 |
|  | 70 | 3098.144917 | 3538.740811 | 4008.053553 | 4331.941438 | 4673.083313 | 5019.073498 | 5356.826318 | 5863.272905 | 6358.685383 |
|  | 80 | 2735.324398 | 3281.285124 | 3827.106379 | 4186.416403 | 4555.617919 | 4927.580124 | 5294.992944 | 5863.915233 | 6447.448971 |
|  | 90 | 2300.180689 | 3022.094398 | 3691.902183 | 4107.098616 | 4520.488326 | 4933.589051 | 5347.98548 | 6016.417293 | 6743.263513 |
| **Left**  **Pallidum** | 6 | 1458.503664 | 1611.251674 | 1788.615308 | 1919.81557 | 2064.73902 | 2216.944718 | 2368.574106 | 2598.243621 | 2823.351595 |
|  | 10 | 1363.713599 | 1512.856323 | 1686.203542 | 1814.464122 | 1956.050335 | 2104.503175 | 2252.003602 | 2474.473224 | 2691.30013 |
|  | 15 | 1255.16412 | 1399.906568 | 1568.259132 | 1692.807137 | 1830.137574 | 1973.781486 | 2115.993959 | 2329.2965 | 2535.6827 |
|  | 20 | 1172.385469 | 1313.785168 | 1478.205783 | 1599.729759 | 1733.487742 | 1872.976422 | 2010.512688 | 2215.552823 | 2412.416384 |
|  | 30 | 1115.606895 | 1257.752433 | 1422.435614 | 1543.620357 | 1676.282012 | 1813.570332 | 1947.651626 | 2144.872814 | 2331.117827 |
|  | 40 | 1054.480313 | 1199.659105 | 1367.148398 | 1489.811101 | 1623.338768 | 1760.466918 | 1893.14619 | 2085.798415 | 2264.890347 |
|  | 50 | 987.1217384 | 1133.22151 | 1300.717704 | 1422.617421 | 1554.439734 | 1688.68181 | 1817.303437 | 2001.633789 | 2170.340605 |
|  | 60 | 969.3963866 | 1117.802301 | 1286.468297 | 1408.247952 | 1538.935462 | 1670.808454 | 1795.877617 | 1972.763772 | 2132.186411 |
|  | 70 | 909.0797686 | 1065.244477 | 1241.026725 | 1366.875595 | 1500.875556 | 1634.865125 | 1760.685287 | 1936.397221 | 2092.47575 |
|  | 80 | 774.5969852 | 953.5761669 | 1152.000816 | 1292.326137 | 1440.218916 | 1586.494933 | 1722.327375 | 1909.472227 | 2073.221757 |
|  | 90 | 620.1108492 | 846.670487 | 1092.006358 | 1261.986995 | 1438.377424 | 1610.26036 | 1767.661493 | 1981.138297 | 2164.840864 |
| **Right Pallidum** | 6 | 1430.319015 | 1546.368157 | 1683.301775 | 1786.445139 | 1902.665246 | 2027.888319 | 2156.46496 | 2359.633338 | 2569.73742 |
|  | 10 | 1351.333779 | 1464.769242 | 1597.771035 | 1697.412606 | 1809.260214 | 1929.442071 | 2052.662477 | 2247.270675 | 2448.561759 |
|  | 15 | 1264.133715 | 1374.997139 | 1503.91187 | 1599.815764 | 1706.939408 | 1821.640831 | 1939.025219 | 2124.30818 | 2316.015672 |
|  | 20 | 1202.127204 | 1312.430636 | 1439.586728 | 1533.491553 | 1637.848042 | 1749.182224 | 1862.907629 | 2042.318409 | 2228.026818 |
|  | 30 | 1153.202808 | 1269.594139 | 1401.289323 | 1497.035797 | 1602.295983 | 1713.746728 | 1827.158457 | 2005.908123 | 2191.138462 |
|  | 40 | 1096.583292 | 1219.211953 | 1355.132259 | 1452.273642 | 1557.831519 | 1668.703822 | 1781.089242 | 1958.096554 | 2141.801589 |
|  | 50 | 1032.87303 | 1161.945104 | 1301.760572 | 1399.827594 | 1505.061579 | 1614.657754 | 1725.31127 | 1899.512971 | 2080.663029 |
|  | 60 | 989.1892857 | 1128.731177 | 1276.045783 | 1377.259981 | 1484.408256 | 1594.99776 | 1706.207666 | 1881.258244 | 2063.737278 |
|  | 70 | 926.3869121 | 1075.729663 | 1228.867945 | 1331.709375 | 1438.995814 | 1548.677408 | 1658.529008 | 1831.471537 | 2012.286645 |
|  | 80 | 843.9560222 | 1001.755315 | 1158.308984 | 1260.819304 | 1366.077297 | 1472.605175 | 1578.865019 | 1746.242747 | 1921.851212 |
|  | 90 | 790.954227 | 965.4476954 | 1132.277341 | 1238.502228 | 1345.719145 | 1453.078715 | 1559.733048 | 1727.889394 | 1905.029392 |
| **Left**  **Hippocampus** | 6 | 3209.830065 | 3538.235306 | 3860.271045 | 4069.194823 | 4280.565316 | 4488.822548 | 4688.706726 | 4985.63953 | 5274.968176 |
|  | 10 | 3210.781104 | 3535.618644 | 3858.769005 | 4070.493114 | 4285.469859 | 4496.831244 | 4698.134298 | 4992.826859 | 5274.371985 |
|  | 15 | 3204.041788 | 3525.414024 | 3850.301773 | 4065.488114 | 4284.80481 | 4499.842245 | 4702.79512 | 4994.901829 | 5267.686082 |
|  | 20 | 3179.085091 | 3498.384397 | 3825.710921 | 4044.523928 | 4268.191285 | 4486.855718 | 4691.470516 | 4981.382614 | 5246.49005 |
|  | 30 | 3116.95536 | 3440.130027 | 3778.684837 | 4008.157873 | 4243.529345 | 4472.151249 | 4682.69826 | 4972.730913 | 5228.223952 |
|  | 40 | 3054.217157 | 3380.625619 | 3727.940243 | 3965.573035 | 4209.627824 | 4445.109407 | 4658.926459 | 4946.468001 | 5191.896344 |
|  | 50 | 2993.277262 | 3320.904668 | 3673.317833 | 3915.895667 | 4164.967921 | 4403.707643 | 4617.789538 | 4899.847028 | 5134.268672 |
|  | 60 | 2841.214691 | 3181.383599 | 3547.911866 | 3800.147789 | 4058.161647 | 4303.396756 | 4520.547021 | 4801.278181 | 5029.119615 |
|  | 70 | 2504.917523 | 2882.976389 | 3283.669914 | 3556.116592 | 3831.835581 | 4090.510785 | 4316.162673 | 4602.226822 | 4829.128277 |
|  | 80 | 2052.820835 | 2483.900327 | 2924.949439 | 3217.73673 | 3509.024316 | 3777.747012 | 4008.308376 | 4294.927219 | 4517.400869 |
|  | 90 | 1606.310368 | 2089.462455 | 2565.354839 | 2871.709562 | 3170.282446 | 3440.70951 | 3668.917794 | 3947.469996 | 4159.528416 |
| **Right**  **Hippocampus** | 6 | 3446.052089 | 3661.469099 | 3914.272171 | 4101.584866 | 4306.117797 | 4514.95312 | 4714.07258 | 4995.609279 | 5248.034315 |
|  | 10 | 3430.061551 | 3654.181825 | 3915.01952 | 4106.961027 | 4315.506639 | 4527.593007 | 4729.266961 | 5013.87968 | 5268.787955 |
|  | 15 | 3400.413931 | 3635.745877 | 3906.611899 | 4104.132285 | 4317.34664 | 4533.065944 | 4737.486876 | 5025.297219 | 5282.736403 |
|  | 20 | 3353.27216 | 3599.809079 | 3880.205856 | 4082.713039 | 4299.820227 | 4518.303919 | 4724.612246 | 5014.396144 | 5273.28325 |
|  | 30 | 3271.2435 | 3544.823485 | 3847.733458 | 4061.902664 | 4288.148336 | 4513.254065 | 4724.2578 | 5019.246109 | 5282.195413 |
|  | 40 | 3174.65865 | 3480.784859 | 3809.146713 | 4035.75661 | 4271.268243 | 4502.736198 | 4718.042132 | 5017.65387 | 5284.225228 |
|  | 50 | 3041.693556 | 3390.229554 | 3749.918281 | 3991.244541 | 4237.488784 | 4476.270575 | 4696.566374 | 5001.68405 | 5272.711028 |
|  | 60 | 2822.968991 | 3225.192331 | 3620.588534 | 3877.031721 | 4133.208983 | 4377.879753 | 4601.572589 | 4909.830406 | 5183.194139 |
|  | 70 | 2484.861213 | 2949.994172 | 3380.828859 | 3648.980726 | 3910.27181 | 4155.52208 | 4377.468667 | 4681.605368 | 4950.823021 |
|  | 80 | 2067.789722 | 2599.301759 | 3063.606166 | 3339.139281 | 3600.034278 | 3840.139906 | 4054.985028 | 4347.601859 | 4606.134361 |
|  | 90 | 1651.118602 | 2230.699887 | 2724.145218 | 3003.32851 | 3259.569845 | 3490.409728 | 3694.49297 | 3970.734402 | 4214.403041 |
| **Left**  **Amygdala** | 6 | 1136.194941 | 1245.505854 | 1378.141267 | 1478.892501 | 1590.518801 | 1705.278868 | 1814.548737 | 1967.468687 | 2102.114189 |
|  | 10 | 1145.495335 | 1254.704083 | 1387.494147 | 1488.601099 | 1600.913871 | 1716.763379 | 1827.491075 | 1983.226191 | 2121.154711 |
|  | 15 | 1151.234848 | 1259.903032 | 1392.362156 | 1493.503692 | 1606.215009 | 1722.957055 | 1835.073573 | 1993.764461 | 2135.365634 |
|  | 20 | 1146.433235 | 1254.208896 | 1385.828052 | 1486.561424 | 1599.129817 | 1716.15989 | 1829.054396 | 1989.810281 | 2134.28961 |
|  | 30 | 1134.664629 | 1244.099771 | 1377.587714 | 1479.747464 | 1594.073108 | 1713.314842 | 1828.905094 | 1994.741395 | 2145.231288 |
|  | 40 | 1110.546039 | 1225.60089 | 1364.969706 | 1471.040296 | 1589.332113 | 1712.48395 | 1831.864639 | 2003.47448 | 2159.778552 |
|  | 50 | 1064.412673 | 1187.668016 | 1334.29559 | 1444.16195 | 1565.218829 | 1689.931928 | 1809.841866 | 1980.970472 | 2135.89525 |
|  | 60 | 978.0071728 | 1117.699151 | 1276.606425 | 1391.278178 | 1513.974521 | 1637.015781 | 1752.640716 | 1913.907877 | 2056.722738 |
|  | 70 | 804.9612075 | 992.0285441 | 1186.27888 | 1317.183696 | 1450.609368 | 1578.90727 | 1695.431918 | 1852.715576 | 1987.847409 |
|  | 80 | 528.6658953 | 785.5743947 | 1031.81151 | 1182.634586 | 1326.606796 | 1457.978731 | 1572.700776 | 1722.256726 | 1846.963958 |
|  | 90 | 349.0550664 | 597.7200569 | 875.8065097 | 1040.420254 | 1188.660238 | 1317.212917 | 1425.342563 | 1561.936578 | 1672.968571 |
| **Right**  **Amygdala** | 6 | 1220.487846 | 1317.097432 | 1440.797638 | 1538.944478 | 1650.948976 | 1768.424624 | 1881.232101 | 2038.707716 | 2175.562942 |
|  | 10 | 1219.150822 | 1317.007335 | 1442.182282 | 1541.502128 | 1654.978032 | 1774.29706 | 1889.300898 | 2050.750372 | 2192.072155 |
|  | 15 | 1214.596914 | 1313.894115 | 1440.719056 | 1541.328985 | 1656.434586 | 1777.849467 | 1895.433453 | 2061.724331 | 2208.668469 |
|  | 20 | 1204.050887 | 1304.548272 | 1432.641149 | 1534.190848 | 1650.492589 | 1773.532745 | 1893.255987 | 2063.845814 | 2216.069758 |
|  | 30 | 1190.954062 | 1294.169965 | 1425.010481 | 1528.498095 | 1647.202815 | 1773.52155 | 1897.659777 | 2077.419596 | 2241.292455 |
|  | 40 | 1176.759591 | 1284.09693 | 1418.495667 | 1523.87603 | 1644.290406 | 1772.524127 | 1899.244328 | 2084.924906 | 2257.156202 |
|  | 50 | 1137.449503 | 1254.135348 | 1396.330633 | 1505.323287 | 1627.889628 | 1756.918557 | 1883.666513 | 2069.186342 | 2241.87933 |
|  | 60 | 1054.637489 | 1190.271077 | 1347.901053 | 1463.977201 | 1590.690879 | 1720.888435 | 1846.624477 | 2028.334004 | 2196.063224 |
|  | 70 | 915.2287776 | 1087.008763 | 1272.241839 | 1400.841004 | 1535.550589 | 1669.518662 | 1796.030694 | 1975.870694 | 2140.078891 |
|  | 80 | 714.477325 | 940.826773 | 1162.31484 | 1305.125847 | 1447.826302 | 1584.999943 | 1711.923799 | 1890.273587 | 2052.467631 |
|  | 90 | 507.6301665 | 783.5132412 | 1043.730732 | 1199.666132 | 1347.972067 | 1485.851758 | 1611.245516 | 1786.398716 | 1946.160484 |
| **Left Nucleus**  **Accumbens** | 6 | 509.3369909 | 560.4184841 | 629.4269536 | 686.7857237 | 754.6437111 | 828.0265489 | 900.071784 | 1002.300364 | 1091.950146 |
|  | 10 | 458.3792538 | 508.751442 | 576.4644938 | 632.5205011 | 698.671797 | 770.128433 | 840.3183835 | 940.162862 | 1028.106402 |
|  | 15 | 406.0645393 | 456.1769606 | 522.9965844 | 577.9434024 | 642.5049003 | 712.0848747 | 780.4457886 | 877.9926835 | 964.423105 |
|  | 20 | 373.0829286 | 424.3734726 | 491.9933426 | 547.0725277 | 611.3761519 | 680.4094219 | 748.1820008 | 845.1494629 | 931.5865044 |
|  | 30 | 349.2403228 | 404.1216383 | 474.1379547 | 529.6124416 | 593.1225965 | 660.3665002 | 725.9519993 | 819.8330895 | 904.1453579 |
|  | 40 | 309.3652364 | 369.4266387 | 444.2740041 | 502.3737539 | 567.8927645 | 636.4631127 | 702.8809435 | 797.6673148 | 882.8753486 |
|  | 50 | 272.2874076 | 333.6828459 | 409.4274302 | 467.585028 | 532.414372 | 599.302284 | 663.0643628 | 752.2097612 | 830.4676983 |
|  | 60 | 252.9652083 | 313.6505376 | 388.2519102 | 445.179573 | 507.9983186 | 571.7605516 | 631.2414436 | 711.8677256 | 780.0074334 |
|  | 70 | 180.7827393 | 251.5697303 | 337.9710893 | 403.2580129 | 474.4331524 | 545.465816 | 610.3700521 | 695.9001277 | 765.7832906 |
|  | 80 | 96.70762039 | 176.5210051 | 275.8783393 | 349.4228883 | 427.8446346 | 504.271982 | 572.4526354 | 659.7782337 | 728.9196727 |
|  | 90 | 56.14584401 | 128.7813802 | 231.0222546 | 306.3907411 | 384.8817681 | 459.4456231 | 524.3976837 | 605.4521442 | 667.9165314 |
| **Right Nucleus**  **Accumbens** | 6 | 520.2835117 | 572.0952781 | 639.8024443 | 694.2143861 | 756.5557293 | 821.6701392 | 883.4405385 | 967.7234992 | 1038.663622 |
|  | 10 | 492.3327807 | 543.6601531 | 610.6250437 | 664.3649907 | 725.8748013 | 790.075919 | 850.9619036 | 934.0476094 | 1004.017239 |
|  | 15 | 459.7529342 | 510.582179 | 576.7524395 | 629.7545035 | 690.3376418 | 753.5112302 | 813.397383 | 895.1279804 | 964.0019979 |
|  | 20 | 432.4273376 | 483.1945903 | 549.1341752 | 601.8479698 | 662.0174585 | 724.6988756 | 784.0942044 | 865.1682407 | 933.5398196 |
|  | 30 | 393.5582945 | 445.8903059 | 513.535994 | 567.3897757 | 628.6809067 | 692.404365 | 752.7411936 | 835.1414278 | 904.7536324 |
|  | 40 | 356.5435666 | 411.0184329 | 480.9478296 | 536.2810783 | 598.9709491 | 663.9223504 | 725.2985028 | 809.0611655 | 879.8758114 |
|  | 50 | 316.2260633 | 373.6368969 | 446.3749232 | 503.2652046 | 567.1363895 | 632.798314 | 694.4904212 | 778.3128495 | 848.9814379 |
|  | 60 | 273.4829908 | 335.4499014 | 412.2238826 | 471.133594 | 536.3047312 | 602.4507556 | 663.9887162 | 746.9097447 | 816.38217 |
|  | 70 | 227.8520501 | 296.662306 | 379.0444006 | 440.5399904 | 507.2118142 | 573.7365293 | 634.8331285 | 716.2724036 | 783.9440501 |
|  | 80 | 176.4863346 | 254.1511658 | 343.155199 | 407.1938945 | 474.8592009 | 540.9816308 | 600.7898235 | 679.5289723 | 744.3596607 |
|  | 90 | 125.5274097 | 211.3059302 | 308.0423107 | 374.8989652 | 443.4540063 | 508.8724462 | 567.0588543 | 642.6648769 | 704.3411915 |

| **Table S7. Centile Values for Subcortical Volumes in Males** | | | | | | | | | | |
| --- | --- | --- | --- | --- | --- | --- | --- | --- | --- | --- |
| **Region** | **Age** | **C0.4** | **C2** | **C10** | **C25** | **C50** | **C75** | **C90** | **C98** | **C99.6** |
| **Left**  **Lateral**  **Ventricle** | 6 | 387.0341483 | 747.7458825 | 1529.590799 | 2495.891861 | 4084.389304 | 6438.690321 | 9497.069649 | 15319.11225 | 22083.30252 |
|  | 10 | 469.4135503 | 870.1070923 | 1711.845436 | 2730.007865 | 4380.696222 | 6805.501484 | 9944.139215 | 15927.35755 | 22923.05365 |
|  | 15 | 589.9165877 | 1044.971143 | 1966.624997 | 3052.825239 | 4783.765896 | 7297.842374 | 10536.60367 | 16720.81967 | 24010.67094 |
|  | 20 | 734.4263364 | 1250.058675 | 2258.871759 | 3417.796993 | 5232.800655 | 7838.129412 | 11177.04747 | 17561.15312 | 25149.05668 |
|  | 30 | 1084.32676 | 1730.268654 | 2916.625203 | 4214.811374 | 6180.03309 | 8935.796947 | 12431.41521 | 19138.41729 | 27258.7732 |
|  | 40 | 1506.154999 | 2290.846348 | 3651.924377 | 5074.435341 | 7158.713875 | 10016.70425 | 13609.41229 | 20544.24135 | 29134.69746 |
|  | 50 | 2143.265524 | 3133.186351 | 4752.707936 | 6365.083626 | 8646.063434 | 11698.11612 | 15496.08119 | 22873.79996 | 32248.53619 |
|  | 60 | 3342.382923 | 4694.231748 | 6758.402859 | 8696.51393 | 11322.33947 | 14726.15403 | 18895.57196 | 27006.98639 | 37547.62145 |
|  | 70 | 5497.09661 | 7405.51768 | 10090.45994 | 12440.91912 | 15465.15856 | 19236.98927 | 23761.95863 | 32538.09596 | 44156.27957 |
|  | 80 | 8799.650934 | 11397.71831 | 14731.76072 | 17430.48237 | 20713.15125 | 24644.8925 | 29266.74777 | 38221.9742 | 50328.55302 |
|  | 90 | 12888.2044 | 16214.83154 | 20095.72739 | 22993.4485 | 26330.83778 | 30191.75544 | 34680.5429 | 43491.60124 | 55854.02524 |
| **Right**  **Lateral**  **Ventricle** | 6 | 336.0282848 | 660.6972485 | 1366.792306 | 2244.498739 | 3707.767202 | 5937.879193 | 8959.537912 | 15120.96895 | 22953.82835 |
|  | 10 | 420.3927699 | 788.9949825 | 1559.134947 | 2488.469746 | 4005.028322 | 6278.317694 | 9325.521698 | 15499.82106 | 23342.8108 |
|  | 15 | 546.4238729 | 974.6136596 | 1829.087162 | 2824.874954 | 4408.955663 | 6736.31631 | 9815.284417 | 16005.53507 | 23861.41105 |
|  | 20 | 693.4670613 | 1184.994366 | 2126.648085 | 3189.837471 | 4842.014953 | 7224.976698 | 10339.84757 | 16559.95199 | 24453.4472 |
|  | 30 | 1026.638353 | 1644.537447 | 2751.117203 | 3935.271558 | 5703.639645 | 8176.263192 | 11346.86592 | 17623.95396 | 25627.88632 |
|  | 40 | 1393.644148 | 2142.719243 | 3417.666523 | 4726.576993 | 6622.968881 | 9216.441513 | 12504.77193 | 19017.09978 | 27439.76209 |
|  | 50 | 1936.659697 | 2881.431031 | 4413.945413 | 5925.560771 | 8052.656204 | 10901.20013 | 14477.51005 | 21577.84728 | 30921.14663 |
|  | 60 | 2977.675858 | 4256.446065 | 6208.825305 | 8038.624447 | 10516.61837 | 13737.60692 | 17710.06186 | 25546.48303 | 35944.29621 |
|  | 70 | 4955.734116 | 6735.239874 | 9250.988731 | 11460.18577 | 14307.55346 | 17861.66671 | 22124.65593 | 30379.4894 | 41269.79532 |
|  | 80 | 8154.368606 | 10523.55698 | 13602.20521 | 16120.55317 | 19198.56949 | 22877.99214 | 27160.85204 | 35276.08098 | 45864.6518 |
|  | 90 | 12203.10255 | 15119.84876 | 18621.55349 | 21303.14658 | 24429.7376 | 28036.29249 | 32144.94558 | 39843.4295 | 49883.09905 |
| **Left**  **Thalamus** | 6 | 6440.882403 | 6774.288808 | 7199.211104 | 7529.320013 | 7892.3134 | 8251.580389 | 8571.939037 | 8975.592933 | 9285.555808 |
|  | 10 | 6391.108422 | 6718.515289 | 7147.87217 | 7490.671813 | 7876.856484 | 8268.51438 | 8625.60496 | 9085.984148 | 9447.345718 |
|  | 15 | 6361.852786 | 6675.249484 | 7099.983418 | 7450.588962 | 7858.124703 | 8285.369144 | 8687.439448 | 9223.911494 | 9659.639009 |
|  | 20 | 6340.27254 | 6644.68058 | 7065.691827 | 7420.821206 | 7842.561019 | 8295.371453 | 8731.813967 | 9330.264724 | 9830.347728 |
|  | 30 | 6136.863285 | 6498.966224 | 6986.25425 | 7385.406115 | 7845.786392 | 8324.250949 | 8770.603935 | 9360.264547 | 9834.275118 |
|  | 40 | 6021.292589 | 6381.457282 | 6867.736085 | 7267.389576 | 7729.789472 | 8211.941251 | 8663.150659 | 9261.258145 | 9743.684799 |
|  | 50 | 5805.914634 | 6126.245416 | 6577.109501 | 6964.553157 | 7433.23252 | 7946.897273 | 8452.35256 | 9162.163761 | 9770.328391 |
|  | 60 | 5446.35923 | 5789.148375 | 6247.702364 | 6621.00501 | 7049.034137 | 7491.060002 | 7900.895012 | 8438.682497 | 8868.084896 |
|  | 70 | 4627.654852 | 5136.649366 | 5752.623353 | 6210.684031 | 6697.29501 | 7163.861391 | 7568.932194 | 8066.427093 | 8439.756051 |
|  | 80 | 4698.928834 | 5046.115403 | 5537.015783 | 5960.255394 | 6473.138027 | 7035.494007 | 7588.299685 | 8362.395068 | 9022.600418 |
|  | 90 | 5441.414268 | 5586.902136 | 5815.14093 | 6040.968242 | 6367.601212 | 6831.567763 | 7479.584354 | 9082.288694 | 11800.6391 |
| **Right**  **Thalamus** | 6 | 6509.958 | 6791.534402 | 7151.572781 | 7439.183443 | 7772.373962 | 8130.873632 | 8487.190574 | 9010.42047 | 9495.371468 |
|  | 10 | 6347.448023 | 6647.418553 | 7026.587227 | 7326.589679 | 7671.798875 | 8041.449482 | 8407.97777 | 8946.103245 | 9445.798945 |
|  | 15 | 6146.883598 | 6473.081774 | 6878.544074 | 7194.896377 | 7555.358263 | 7938.640803 | 8317.349951 | 8873.126502 | 9390.491437 |
|  | 20 | 5976.227025 | 6333.683886 | 6769.140551 | 7103.249149 | 7479.496278 | 7876.211137 | 8266.512375 | 8838.884663 | 9373.058951 |
|  | 30 | 5732.67599 | 6167.975491 | 6671.309901 | 7041.387002 | 7446.190825 | 7864.510521 | 8272.178101 | 8869.858022 | 9432.062362 |
|  | 40 | 5512.65103 | 6009.334054 | 6550.476464 | 6930.498406 | 7334.745729 | 7746.368317 | 8147.283243 | 8743.14427 | 9317.991484 |
|  | 50 | 5212.737096 | 5744.529737 | 6292.002481 | 6660.819822 | 7045.095131 | 7435.113883 | 7820.589053 | 8414.131684 | 9016.954532 |
|  | 60 | 4855.152588 | 5426.316643 | 5985.015629 | 6347.570599 | 6719.921765 | 7100.935119 | 7489.379776 | 8124.815963 | 8826.101761 |
|  | 70 | 4400.391358 | 5032.818345 | 5624.175488 | 5994.333241 | 6370.478589 | 6762.618517 | 7182.522791 | 7935.518966 | 8878.505726 |
|  | 80 | 4011.474412 | 4708.150068 | 5331.486059 | 5705.47945 | 6080.621941 | 6481.851647 | 6941.529247 | 7881.313475 | 9305.376338 |
|  | 90 | 3705.767278 | 4479.370856 | 5137.671191 | 5509.830016 | 5874.113219 | 6274.152207 | 6772.873017 | 7991.453702 | 10460.02019 |
| **Left**  **Caudate** | 6 | 3048.889921 | 3322.503325 | 3670.992412 | 3944.01802 | 4249.46905 | 4560.405414 | 4847.972545 | 5229.148185 | 5540.373947 |
|  | 10 | 2983.108271 | 3248.844477 | 3587.133599 | 3852.408769 | 4149.889567 | 4453.964177 | 4736.753154 | 5114.583721 | 5426.104704 |
|  | 15 | 2902.506223 | 3159.559192 | 3486.179276 | 3742.365698 | 4030.380936 | 4326.301802 | 4603.549629 | 4978.049295 | 5291.0857 |
|  | 20 | 2829.326254 | 3079.480485 | 3396.229352 | 3644.448434 | 3924.060046 | 4212.831872 | 4485.54722 | 4858.469309 | 5175.117707 |
|  | 30 | 2722.24672 | 2965.003575 | 3268.41987 | 3504.718297 | 3771.382624 | 4049.610891 | 4317.159582 | 4693.983684 | 5026.658329 |
|  | 40 | 2622.741767 | 2864.622906 | 3160.008213 | 3386.883938 | 3642.415089 | 3911.747573 | 4176.510881 | 4564.178803 | 4925.062426 |
|  | 50 | 2514.117887 | 2764.665106 | 3059.713074 | 3280.939199 | 3528.406106 | 3792.013781 | 4058.59474 | 4470.236855 | 4883.094535 |
|  | 60 | 2416.251059 | 2693.591932 | 3003.359987 | 3226.958423 | 3473.70785 | 3739.642431 | 4019.047118 | 4484.307029 | 5004.30123 |
|  | 70 | 2296.695157 | 2626.77588 | 2969.583251 | 3203.158421 | 3454.897831 | 3729.839736 | 4034.488641 | 4600.909609 | 5345.925027 |
|  | 80 | 2099.036787 | 2520.918444 | 2920.879612 | 3171.245484 | 3430.73946 | 3718.548565 | 4062.715353 | 4818.335063 | 6106.54265 |
|  | 90 | 1764.782176 | 2339.522214 | 2837.065398 | 3113.713739 | 3382.999033 | 3687.133987 | 4093.841835 | 5248.594515 | 8300.964194 |
| **Right**  **Caudate** | 6 | 3175.320489 | 3437.417866 | 3777.005335 | 4045.877078 | 4347.58805 | 4653.528584 | 4933.513403 | 5297.574965 | 5587.021232 |
|  | 10 | 3091.412525 | 3346.222825 | 3677.646598 | 3941.220491 | 4238.389942 | 4541.437782 | 4820.467893 | 5185.985026 | 5478.979063 |
|  | 15 | 2989.920186 | 3236.442749 | 3558.362716 | 3815.66496 | 4107.440608 | 4407.160442 | 4685.379103 | 5053.580466 | 5352.166369 |
|  | 20 | 2898.290293 | 3137.799237 | 3451.396364 | 3703.085251 | 3990.041194 | 4286.990693 | 4565.045097 | 4937.251336 | 5243.127077 |
|  | 30 | 2780.612752 | 3008.051303 | 3305.392677 | 3544.774423 | 3819.837096 | 4108.362193 | 4383.498338 | 4761.542595 | 5082.378998 |
|  | 40 | 2693.386867 | 2913.125245 | 3196.080931 | 3422.136544 | 3682.156029 | 3957.656102 | 4225.300751 | 4604.641909 | 4940.217368 |
|  | 50 | 2596.598594 | 2821.174378 | 3101.20279 | 3320.29174 | 3570.686421 | 3838.058069 | 4103.62591 | 4496.173862 | 4864.655805 |
|  | 60 | 2494.420854 | 2748.287328 | 3047.565142 | 3272.559256 | 3525.523006 | 3797.387017 | 4075.760454 | 4513.915302 | 4964.521837 |
|  | 70 | 2334.355125 | 2657.718502 | 3005.74021 | 3249.616946 | 3515.063603 | 3801.860957 | 4109.265305 | 4643.840465 | 5280.466764 |
|  | 80 | 2039.907693 | 2511.196159 | 2954.548905 | 3230.410027 | 3513.409804 | 3820.647389 | 4175.406615 | 4904.271632 | 6014.933349 |
|  | 90 | 1419.75829 | 2212.682164 | 2857.241202 | 3189.312943 | 3496.131393 | 3831.354257 | 4271.914063 | 5484.918118 | 8378.61899 |
| **Left**  **Putamen** | 6 | 5275.396377 | 5706.977339 | 6245.755552 | 6658.215804 | 7108.179672 | 7552.255967 | 7949.183128 | 8453.421366 | 8845.846911 |
|  | 10 | 5011.419727 | 5423.291637 | 5938.265296 | 6333.288931 | 6765.244673 | 7192.847549 | 7576.380484 | 8065.769882 | 8448.58675 |
|  | 15 | 4713.727595 | 5103.874291 | 5592.35428 | 5967.855744 | 6379.622592 | 6788.829327 | 7157.565878 | 7630.949654 | 8003.897924 |
|  | 20 | 4499.159818 | 4874.850419 | 5345.520091 | 5707.927311 | 6106.387266 | 6503.953926 | 6863.997468 | 7329.376115 | 7699.012499 |
|  | 30 | 4305.361876 | 4673.884295 | 5134.789093 | 5490.162658 | 5882.669897 | 6277.562703 | 6639.274722 | 7114.502861 | 7499.635861 |
|  | 40 | 4100.871443 | 4465.55589 | 4918.4574 | 5266.871591 | 5652.874306 | 6044.6416 | 6408.476196 | 6896.767908 | 7303.405678 |
|  | 50 | 3808.657095 | 4167.709339 | 4607.08695 | 4942.516756 | 5314.410805 | 5695.379982 | 6055.345599 | 6552.42458 | 6982.285433 |
|  | 60 | 3581.076973 | 3949.933391 | 4389.837607 | 4720.443187 | 5085.945884 | 5464.093641 | 5829.45299 | 6354.232708 | 6832.948979 |
|  | 70 | 3351.401562 | 3746.024466 | 4197.698224 | 4527.919871 | 4889.952255 | 5268.526529 | 5645.371542 | 6217.71058 | 6781.787808 |
|  | 80 | 3085.034018 | 3529.553345 | 4007.903314 | 4342.21334 | 4702.583352 | 5083.844989 | 5479.359443 | 6130.920007 | 6850.984787 |
|  | 90 | 2721.312048 | 3253.078999 | 3778.142897 | 4120.129439 | 4477.824869 | 4861.244695 | 5283.242595 | 6068.490708 | 7101.792189 |
| **Right**  **Putamen** | 6 | 5196.91273 | 5546.775849 | 5992.56834 | 6343.025167 | 6737.567683 | 7143.175059 | 7523.054374 | 8035.443466 | 8462.687848 |
|  | 10 | 4943.07681 | 5286.323314 | 5721.680477 | 6062.822355 | 6446.263476 | 6840.380969 | 7209.967276 | 7709.994022 | 8128.86072 |
|  | 15 | 4654.290237 | 4990.752376 | 5414.772535 | 5745.521977 | 6116.450366 | 6497.603989 | 6855.669733 | 7342.18209 | 7752.369399 |
|  | 20 | 4436.002914 | 4770.088059 | 5188.047874 | 5512.38229 | 5875.185719 | 6247.86944 | 6598.679706 | 7077.637524 | 7484.411878 |
|  | 30 | 4187.374226 | 4528.633134 | 4947.991018 | 5269.274709 | 5626.380981 | 5992.866459 | 6339.470242 | 6818.166924 | 7231.876258 |
|  | 40 | 3934.763393 | 4283.487533 | 4702.282359 | 5017.893893 | 5365.834956 | 5722.497667 | 6061.879692 | 6537.618765 | 6958.074218 |
|  | 50 | 3641.180096 | 4001.447484 | 4421.808569 | 4732.114324 | 5070.769031 | 5417.553096 | 5750.289241 | 6225.930567 | 6658.691716 |
|  | 60 | 3361.824284 | 3750.121462 | 4187.205323 | 4501.602843 | 4840.488603 | 5187.262165 | 5523.772956 | 6017.398575 | 6483.80463 |
|  | 70 | 3087.119525 | 3525.627355 | 3997.531171 | 4325.994411 | 4674.546087 | 5031.040106 | 5382.235383 | 5915.118672 | 6443.75847 |
|  | 80 | 2780.761369 | 3295.513584 | 3819.144328 | 4168.542237 | 4531.912775 | 4903.406492 | 5276.692257 | 5868.568754 | 6493.573835 |
|  | 90 | 2392.371334 | 3012.491844 | 3601.602457 | 3973.935557 | 4351.136625 | 4736.587442 | 5134.069531 | 5801.393408 | 6564.499653 |
| **Left**  **Pallidum** | 6 | 1472.79607 | 1631.634206 | 1810.078638 | 1938.568219 | 2077.856186 | 2222.110164 | 2364.627944 | 2579.567985 | 2790.03578 |
|  | 10 | 1374.763158 | 1534.158141 | 1714.249465 | 1844.463457 | 1985.947358 | 2132.612462 | 2277.438458 | 2495.400407 | 2708.087847 |
|  | 15 | 1263.606525 | 1423.336328 | 1605.023126 | 1737.028237 | 1880.835348 | 2030.047462 | 2177.265482 | 2398.200645 | 2612.811808 |
|  | 20 | 1175.313666 | 1333.494387 | 1514.365726 | 1646.259386 | 1790.192256 | 1939.548793 | 2086.678783 | 2306.687187 | 2519.2621 |
|  | 30 | 1097.222463 | 1249.648499 | 1424.806342 | 1552.913654 | 1692.741151 | 1837.438206 | 1979.175851 | 2189.056735 | 2389.191698 |
|  | 40 | 1041.387435 | 1193.332547 | 1368.803653 | 1497.516451 | 1638.03656 | 1783.075082 | 1924.393346 | 2131.733994 | 2327.012212 |
|  | 50 | 979.1068071 | 1126.616458 | 1297.537479 | 1423.123822 | 1560.143228 | 1701.110492 | 1837.67691 | 2036.165603 | 2220.80328 |
|  | 60 | 966.2705303 | 1111.834259 | 1280.750463 | 1404.895316 | 1540.122609 | 1678.690685 | 1812.100692 | 2004.128622 | 2180.544007 |
|  | 70 | 912.51532 | 1062.449132 | 1237.07327 | 1365.652304 | 1505.656268 | 1648.724057 | 1785.779694 | 1981.435131 | 2159.244409 |
|  | 80 | 785.5010648 | 949.214283 | 1141.155146 | 1283.033196 | 1437.679235 | 1595.483629 | 1746.086947 | 1959.61223 | 2151.850868 |
|  | 90 | 633.1688059 | 821.6618398 | 1044.374355 | 1209.643496 | 1389.938751 | 1573.614271 | 1748.22334 | 1994.080503 | 2213.3772 |
| **Right**  **Pallidum** | 6 | 1364.616213 | 1494.644741 | 1645.158661 | 1756.636713 | 1880.68189 | 2012.996089 | 2147.916945 | 2359.952682 | 2578.17603 |
|  | 10 | 1308.094344 | 1437.180414 | 1585.510402 | 1694.704503 | 1815.717297 | 1944.464329 | 2075.630471 | 2281.893222 | 2494.55656 |
|  | 15 | 1242.609153 | 1370.899924 | 1516.913035 | 1623.555833 | 1741.125314 | 1865.797205 | 1992.672061 | 2192.360947 | 2398.739306 |
|  | 20 | 1186.337481 | 1314.690195 | 1459.317502 | 1564.081131 | 1678.955342 | 1800.358223 | 1923.773211 | 2118.208541 | 2319.67218 |
|  | 30 | 1111.554511 | 1244.343494 | 1390.781378 | 1495.006926 | 1607.992793 | 1726.565803 | 1846.853477 | 2036.805759 | 2234.73734 |
|  | 40 | 1055.228775 | 1195.5763 | 1346.709906 | 1452.230728 | 1565.224535 | 1682.936319 | 1802.116296 | 1990.859032 | 2188.786588 |
|  | 50 | 997.7950771 | 1146.993216 | 1303.465342 | 1410.439136 | 1523.485168 | 1640.34489 | 1758.447096 | 1946.126227 | 2144.347236 |
|  | 60 | 944.9052918 | 1105.601782 | 1269.228735 | 1378.539419 | 1492.422493 | 1609.196175 | 1727.016465 | 1915.007367 | 2115.138491 |
|  | 70 | 873.2944307 | 1044.542584 | 1213.253475 | 1323.132727 | 1435.865219 | 1550.478506 | 1665.950286 | 1851.064695 | 2049.866947 |
|  | 80 | 792.8182698 | 974.9099623 | 1147.896273 | 1257.460165 | 1368.026261 | 1479.436808 | 1591.543286 | 1772.241569 | 1968.175101 |
|  | 90 | 717.9655303 | 914.2941917 | 1093.773407 | 1204.049116 | 1313.375014 | 1422.510405 | 1532.221046 | 1710.158056 | 1905.13088 |
| **Left Hippocampus** | 6 | 3275.564726 | 3575.178737 | 3888.247654 | 4100.889633 | 4321.581305 | 4541.464506 | 4751.804704 | 5058.770806 | 5349.499828 |
|  | 10 | 3255.09555 | 3555.68787 | 3873.196612 | 4090.412794 | 4316.355667 | 4540.912895 | 4754.207424 | 5061.459128 | 5347.395511 |
|  | 15 | 3224.605727 | 3527.159951 | 3850.53913 | 4073.4769 | 4305.857034 | 4536.060026 | 4752.877001 | 5060.489619 | 5340.984065 |
|  | 20 | 3190.309889 | 3494.969314 | 3823.887111 | 4052.084056 | 4290.276473 | 4525.431995 | 4745.141216 | 5052.487711 | 5327.520158 |
|  | 30 | 3132.847076 | 3440.372502 | 3777.613828 | 4013.777272 | 4260.553743 | 4502.462808 | 4725.155291 | 5028.917392 | 5291.855781 |
|  | 40 | 3071.303262 | 3382.332815 | 3726.915326 | 3969.517913 | 4222.795077 | 4469.2403 | 4693.076061 | 4991.802983 | 5243.166451 |
|  | 50 | 2987.857021 | 3307.755293 | 3663.741534 | 3914.694415 | 4175.89041 | 4427.986853 | 4654.076455 | 4950.026338 | 5193.026093 |
|  | 60 | 2796.163192 | 3132.99207 | 3506.318527 | 3768.310611 | 4039.26255 | 4298.193479 | 4527.429581 | 4822.0993 | 5058.76633 |
|  | 70 | 2445.279125 | 2813.926488 | 3214.928053 | 3492.423336 | 3776.011267 | 4043.38702 | 4276.663077 | 4571.065693 | 4802.597937 |
|  | 80 | 2018.752835 | 2437.986427 | 2877.277955 | 3173.629083 | 3471.076046 | 3746.730401 | 3983.33178 | 4276.422235 | 4502.341795 |
|  | 90 | 1573.229194 | 2052.683776 | 2532.75797 | 2845.489292 | 3152.225873 | 3430.87971 | 3665.967016 | 3951.92608 | 4168.269659 |
| **Right**  **Hippocampus** | 6 | 3506.338825 | 3701.389446 | 3948.294353 | 4142.161793 | 4361.410941 | 4589.272383 | 4806.210504 | 5106.229069 | 5364.506632 |
|  | 10 | 3474.41212 | 3676.967057 | 3931.152361 | 4129.269272 | 4352.099631 | 4582.668514 | 4801.551571 | 5103.736175 | 5363.753682 |
|  | 15 | 3427.693575 | 3640.19755 | 3903.732068 | 4107.105598 | 4334.192091 | 4567.828551 | 4788.822635 | 5093.319666 | 5355.263017 |
|  | 20 | 3373.819776 | 3596.939712 | 3870.055597 | 4078.556893 | 4309.567129 | 4545.828465 | 4768.496881 | 5074.766599 | 5338.282748 |
|  | 30 | 3293.149363 | 3542.315902 | 3838.184491 | 4058.553383 | 4298.543093 | 4540.882884 | 4767.637237 | 5078.697027 | 5346.817712 |
|  | 40 | 3201.387835 | 3482.79916 | 3804.750215 | 4037.649541 | 4286.381484 | 4534.145498 | 4764.390647 | 5079.89003 | 5352.993605 |
|  | 50 | 3053.09256 | 3378.53496 | 3733.881712 | 3982.077978 | 4241.274567 | 4495.675293 | 4730.564645 | 5052.640859 | 5333.384325 |
|  | 60 | 2800.869427 | 3188.009777 | 3585.458351 | 3851.136421 | 4121.28523 | 4382.037401 | 4621.240445 | 4949.97173 | 5239.277981 |
|  | 70 | 2392.243555 | 2860.070629 | 3302.873823 | 3582.693192 | 3858.11442 | 4118.826963 | 4356.387388 | 4684.118289 | 4976.11471 |
|  | 80 | 1913.136749 | 2481.645846 | 2978.187975 | 3271.543734 | 3549.333536 | 3806.50039 | 4039.26303 | 4362.300066 | 4654.624389 |
|  | 90 | 1460.939279 | 2098.321523 | 2647.245007 | 2951.479013 | 3227.749486 | 3477.540576 | 3702.365703 | 4017.27896 | 4307.920003 |
| **Left**  **Amygdala** | 6 | 1203.653862 | 1295.532176 | 1415.051776 | 1511.602898 | 1623.933339 | 1744.549341 | 1863.40471 | 2034.862839 | 2189.606029 |
|  | 10 | 1193.615377 | 1288.04844 | 1410.129853 | 1508.164773 | 1621.625341 | 1742.817703 | 1861.670635 | 2032.250654 | 2185.412601 |
|  | 15 | 1179.630125 | 1277.382798 | 1402.718759 | 1502.584453 | 1617.382919 | 1739.186198 | 1857.921008 | 2027.258234 | 2178.359842 |
|  | 20 | 1166.562868 | 1267.947676 | 1396.796037 | 1498.614829 | 1614.833479 | 1737.298009 | 1855.952781 | 2024.11986 | 2173.27242 |
|  | 30 | 1149.768149 | 1260.15 | 1397.694255 | 1504.443629 | 1624.47194 | 1749.156885 | 1868.484038 | 2035.540334 | 2182.014196 |
|  | 40 | 1118.737737 | 1240.11136 | 1388.009423 | 1500.547728 | 1625.086808 | 1752.57861 | 1873.117124 | 2039.923624 | 2184.669442 |
|  | 50 | 1052.940988 | 1187.127537 | 1346.380387 | 1464.874028 | 1593.74896 | 1723.662557 | 1844.982925 | 2010.988279 | 2153.657679 |
|  | 60 | 950.8252161 | 1100.700502 | 1272.682987 | 1397.238545 | 1530.033971 | 1661.634335 | 1782.916846 | 1946.957658 | 2086.606957 |
|  | 70 | 816.1512357 | 989.3575533 | 1178.997714 | 1311.625937 | 1449.644015 | 1583.726252 | 1705.474914 | 1868.093167 | 2005.169485 |
|  | 80 | 648.6814827 | 852.7830708 | 1062.94315 | 1203.332251 | 1345.106605 | 1479.633967 | 1599.7416 | 1757.982395 | 1889.988841 |
|  | 90 | 476.3429138 | 710.0322389 | 942.0023909 | 1089.216385 | 1232.782221 | 1365.424379 | 1481.681781 | 1632.658896 | 1757.294888 |
| **Right Amygdala** | 6 | 1232.614802 | 1328.989782 | 1457.564877 | 1563.029978 | 1686.132621 | 1817.183547 | 1943.714058 | 2119.610347 | 2270.382742 |
|  | 10 | 1226.053035 | 1323.112271 | 1452.184522 | 1557.817584 | 1680.983704 | 1812.1021 | 1938.842035 | 2115.488419 | 2267.499399 |
|  | 15 | 1217.343656 | 1315.335493 | 1445.062482 | 1550.893394 | 1674.102092 | 1805.262126 | 1932.240525 | 2109.857227 | 2263.537041 |
|  | 20 | 1209.626554 | 1308.82654 | 1439.461588 | 1545.62819 | 1668.990302 | 1800.280956 | 1927.582784 | 2106.332409 | 2261.903733 |
|  | 30 | 1204.270514 | 1307.864179 | 1442.371402 | 1550.526698 | 1675.442557 | 1808.102785 | 1937.014621 | 2119.366675 | 2280.018597 |
|  | 40 | 1187.097785 | 1297.972223 | 1438.88052 | 1550.267486 | 1677.517392 | 1811.821811 | 1942.245547 | 2127.725596 | 2292.94528 |
|  | 50 | 1136.757495 | 1259.310281 | 1410.044078 | 1526.04916 | 1656.178225 | 1791.845808 | 1922.931827 | 2109.713673 | 2277.526836 |
|  | 60 | 1050.870234 | 1192.187823 | 1357.753463 | 1480.259626 | 1614.058079 | 1750.99796 | 1882.168478 | 2069.072414 | 2238.358722 |
|  | 70 | 931.4520983 | 1102.658889 | 1289.509287 | 1420.424851 | 1558.453801 | 1696.479037 | 1827.384554 | 2014.213918 | 2185.419912 |
|  | 80 | 772.2583867 | 989.7773052 | 1204.829781 | 1344.853061 | 1486.097411 | 1623.603317 | 1752.85216 | 1938.490638 | 2111.804215 |
|  | 90 | 588.6165746 | 863.1200329 | 1113.699741 | 1263.410433 | 1406.800878 | 1542.445761 | 1669.201214 | 1853.795501 | 2031.064614 |
| **Left Nucleus**  **Accumbens** | 6 | 486.6956193 | 547.2878891 | 624.1840175 | 685.1651656 | 755.5017423 | 831.1077829 | 906.4779478 | 1017.99569 | 1122.472789 |
|  | 10 | 437.6064563 | 496.5835133 | 571.6571726 | 631.2984055 | 700.1239593 | 774.061662 | 847.6467559 | 956.1834356 | 1057.416141 |
|  | 15 | 387.5516377 | 445.0999842 | 518.5654333 | 577.0049019 | 644.4305868 | 716.7498391 | 788.5194157 | 893.8769966 | 991.5190147 |
|  | 20 | 357.6867249 | 415.0543803 | 488.3093264 | 546.5235667 | 613.5486563 | 685.192251 | 755.9725099 | 859.2095245 | 954.1223612 |
|  | 30 | 340.0184981 | 397.3917072 | 469.9457398 | 527.0055847 | 591.9982904 | 660.580999 | 727.3909017 | 823.1129365 | 909.333591 |
|  | 40 | 309.2997641 | 369.6309789 | 445.4598522 | 504.6520842 | 571.5175544 | 641.3528988 | 708.6032027 | 803.5408376 | 887.6084996 |
|  | 50 | 267.7308236 | 329.5416149 | 406.4885994 | 465.9476864 | 532.4376804 | 601.0713832 | 666.347931 | 757.1020681 | 836.1049141 |
|  | 60 | 238.5142156 | 300.6351962 | 376.6945521 | 434.5926932 | 498.4849016 | 563.5159217 | 624.5081264 | 707.9368116 | 779.3018636 |
|  | 70 | 168.5823234 | 241.0455744 | 328.4930159 | 394.0297595 | 465.3714799 | 536.9699832 | 603.2138326 | 692.4336052 | 767.5123102 |
|  | 80 | 89.56975547 | 169.8755095 | 270.1259956 | 343.9651882 | 422.7753079 | 500.3546333 | 570.900328 | 664.1924836 | 741.2749873 |
|  | 90 | 47.64995278 | 116.9955882 | 218.4524612 | 293.8126709 | 372.7321704 | 448.7729285 | 516.6142049 | 604.6046189 | 675.9531536 |
| **Right Nucleus**  **Accumbens** | 6 | 488.5508053 | 538.1377664 | 605.1333146 | 660.1392956 | 723.6335694 | 789.5992987 | 851.0577142 | 932.0565737 | 996.9651694 |
|  | 10 | 470.4475101 | 520.4068166 | 587.8583991 | 643.191329 | 707.0045774 | 773.232097 | 834.869009 | 916.0059564 | 980.9457408 |
|  | 15 | 448.2825117 | 498.7357776 | 566.7851109 | 622.540557 | 686.7593295 | 753.3118738 | 815.1624036 | 896.4501189 | 961.4049781 |
|  | 20 | 426.6189568 | 477.602188 | 546.2842196 | 602.4798491 | 667.1134647 | 733.9897177 | 796.0440036 | 877.4589124 | 942.4028443 |
|  | 30 | 384.6991347 | 436.8681558 | 506.9357157 | 564.0715871 | 629.5664113 | 697.0866688 | 759.5177791 | 841.1140666 | 905.9565142 |
|  | 40 | 344.3973255 | 397.9461965 | 469.5756444 | 527.733216 | 594.1247704 | 662.2720804 | 725.0250134 | 806.6859385 | 871.3060736 |
|  | 50 | 305.4301035 | 360.5997144 | 433.9979325 | 493.265449 | 560.585 | 629.3305593 | 692.3364058 | 773.926647 | 838.1905252 |
|  | 60 | 267.5511584 | 324.660084 | 400.0885321 | 460.5736512 | 528.8596678 | 598.1737137 | 661.359935 | 742.7399009 | 806.5125959 |
|  | 70 | 230.4777941 | 289.961723 | 367.7595843 | 429.5975312 | 498.9008481 | 568.7567533 | 632.051127 | 713.0828986 | 776.2338397 |
|  | 80 | 193.8429645 | 256.2953034 | 336.8966956 | 400.2500057 | 470.6285479 | 540.9962812 | 604.3212784 | 684.8623107 | 747.2609808 |
|  | 90 | 157.3210752 | 223.4422339 | 307.3791189 | 372.4369116 | 443.9525079 | 514.795597 | 578.0666132 | 657.9688328 | 719.4848101 |

| **Table S8. Centile Values for Subcortical Volumes in Females** | | | | | | | | | | | |
| --- | --- | --- | --- | --- | --- | --- | --- | --- | --- | --- | --- |
| **Region** |  | **Age** | **C0.4** | **C2** | **C10** | **C25** | **C50** | **C75** | **C90** | **C98** | **C99.6** |
| **Left**  **Lateral**  **Ventricle** | 12 | 6 | 545.1216169 | 1078.58624 | 2007.807114 | 2938.843923 | 4255.556466 | 6051.462899 | 8400.47564 | 13467.50553 | 20999.53335 |
|  | 13 | 10 | 723.4633652 | 1305.727128 | 2286.170501 | 3255.762634 | 4619.511333 | 6470.778291 | 8878.045583 | 14022.46007 | 21581.52107 |
|  | 14 | 15 | 989.7525114 | 1633.372379 | 2681.873864 | 3704.494128 | 5133.476758 | 7061.612927 | 9550.421669 | 14808.67226 | 22427.9925 |
|  | 15 | 20 | 1282.275244 | 1979.293903 | 3088.407675 | 4159.281278 | 5648.398736 | 7648.143487 | 10213.97226 | 15584.54542 | 23279.26416 |
|  | 16 | 30 | 1802.001931 | 2542.786093 | 3696.501898 | 4803.650601 | 6339.856433 | 8392.338734 | 10998.03942 | 16339.91599 | 23773.22836 |
|  | 17 | 40 | 2329.797276 | 3131.975958 | 4383.08966 | 5592.685408 | 7281.714966 | 9542.504783 | 12397.85207 | 18158.57313 | 25963.6407 |
|  | 18 | 50 | 2803.320408 | 3692.591795 | 5076.962634 | 6415.135918 | 8281.708237 | 10770.60964 | 13890.87072 | 20097.57239 | 28333.95115 |
|  | 19 | 60 | 3258.519329 | 4366.650239 | 6051.363505 | 7637.714444 | 9797.317162 | 12609.46698 | 16063.60688 | 22791.1779 | 31536.53951 |
|  | 20 | 70 | 4505.581063 | 6257.315748 | 8697.909526 | 10802.39707 | 13465.086 | 16720.56045 | 20541.40981 | 27727.61583 | 36862.02039 |
|  | 21 | 80 | 6656.04142 | 9563.034851 | 13010.14887 | 15577.68418 | 18500.48666 | 21814.07907 | 25557.79611 | 32550.14677 | 41620.41174 |
|  | 22 | 90 | 8811.943032 | 13317.9431 | 17753.71275 | 20564.37308 | 23456.64194 | 26567.35108 | 30079.33722 | 36967.68368 | 46678.45074 |
| **Right**  **Lateral**  **Ventricle** | 23 | 6 | 697.9018839 | 1184.928734 | 2002.552002 | 2818.427738 | 3975.878316 | 5552.244199 | 7591.517182 | 11872.17132 | 17972.4952 |
|  | 24 | 10 | 780.0281619 | 1313.725711 | 2200.032904 | 3075.93753 | 4307.292218 | 5966.824963 | 8088.869458 | 12471.59496 | 18592.425 |
|  | 25 | 15 | 899.2502846 | 1497.179323 | 2476.140437 | 3431.697873 | 4759.703868 | 6526.234296 | 8752.809001 | 13260.88226 | 19404.03226 |
|  | 26 | 20 | 1039.527543 | 1699.720371 | 2763.262323 | 3788.024732 | 5196.180568 | 7045.99998 | 9346.155154 | 13918.18209 | 20010.42963 |
|  | 27 | 30 | 1464.724322 | 2208.377019 | 3363.704053 | 4454.305478 | 5933.929054 | 7855.822406 | 10220.23411 | 14859.6413 | 20958.50131 |
|  | 28 | 40 | 2102.556549 | 2905.14381 | 4127.663102 | 5279.568013 | 6855.308246 | 8934.686693 | 11549.4154 | 16866.65043 | 24228.98988 |
|  | 29 | 50 | 2581.833494 | 3439.734066 | 4740.415873 | 5971.93022 | 7677.6894 | 9977.957081 | 12961.61535 | 19363.50964 | 28978.75645 |
|  | 30 | 60 | 2995.602464 | 4035.607576 | 5581.183267 | 7013.391798 | 8959.643389 | 11538.42341 | 14836.06236 | 21813.47455 | 32149.50445 |
|  | 31 | 70 | 4150.613955 | 5743.559961 | 7943.197156 | 9828.491109 | 12212.396 | 15145.18806 | 18635.39433 | 25380.4701 | 34311.93624 |
|  | 32 | 80 | 6121.033068 | 8719.865176 | 11902.65243 | 14340.66493 | 17154.63069 | 20339.2266 | 23869.21302 | 30183.79003 | 37860.71812 |
|  | 33 | 90 | 8203.891382 | 12140.78803 | 16399.08875 | 19334.3978 | 22478.99882 | 25837.62077 | 29417.8494 | 35625.57743 | 42967.85086 |
| **Left**  **Thalamus** | 34 | 6 | 6369.352705 | 6672.525908 | 7060.969906 | 7367.387838 | 7712.635762 | 8066.965738 | 8397.442038 | 8839.86761 | 9204.986596 |
|  | 35 | 10 | 6371.789847 | 6677.664262 | 7071.026355 | 7382.818878 | 7736.111445 | 8101.338231 | 8444.798302 | 8909.493388 | 9297.68345 |
|  | 36 | 15 | 6367.345442 | 6676.278251 | 7075.273964 | 7393.402747 | 7756.496955 | 8135.478 | 8495.874183 | 8990.661596 | 9411.106065 |
|  | 37 | 20 | 6349.753831 | 6660.998524 | 7064.587055 | 7388.285436 | 7760.556652 | 8153.171817 | 8531.174718 | 9058.778879 | 9515.993037 |
|  | 38 | 30 | 6292.453086 | 6606.209692 | 7015.924239 | 7348.452979 | 7737.219534 | 8157.002716 | 8573.082744 | 9177.714927 | 9728.166911 |
|  | 39 | 40 | 6149.315827 | 6467.234558 | 6881.159739 | 7218.152371 | 7616.060446 | 8053.794022 | 8499.367545 | 9173.922503 | 9822.004086 |
|  | 40 | 50 | 5894.439993 | 6222.765506 | 6641.500133 | 6977.968782 | 7373.983904 | 7812.720113 | 8267.349077 | 8979.489403 | 9698.578502 |
|  | 41 | 60 | 5551.666122 | 5909.348918 | 6344.220791 | 6680.888602 | 7068.754198 | 7495.263146 | 7940.864767 | 8658.425124 | 9416.985174 |
|  | 42 | 70 | 5096.697108 | 5533.537466 | 6017.116783 | 6364.223645 | 6746.280983 | 7157.905657 | 7591.265572 | 8316.921687 | 9136.036307 |
|  | 43 | 80 | 4428.860662 | 5078.803402 | 5685.774385 | 6065.04908 | 6451.087962 | 6856.445404 | 7296.274965 | 8103.625673 | 9148.468587 |
|  | 44 | 90 | 3190.304655 | 4449.183301 | 5358.774316 | 5808.385914 | 6211.356819 | 6622.986424 | 7103.925291 | 8159.193108 | 9881.654224 |
| **Right**  **Thalamus** | 45 | 6 | 6338.998475 | 6616.160682 | 6959.346937 | 7223.894687 | 7519.067284 | 7822.572016 | 8109.530104 | 8504.789585 | 8844.760628 |
|  | 46 | 10 | 6267.373802 | 6549.840058 | 6899.918879 | 7170.41001 | 7473.356509 | 7786.674462 | 8085.129629 | 8500.556037 | 8862.483534 |
|  | 47 | 15 | 6176.807047 | 6465.845935 | 6824.447561 | 7102.346227 | 7415.142371 | 7741.185635 | 8054.926672 | 8497.965278 | 8890.873841 |
|  | 48 | 20 | 6095.48745 | 6391.132691 | 6758.339524 | 7043.803218 | 7366.831073 | 7706.390947 | 8036.760109 | 8510.720821 | 8939.423954 |
|  | 49 | 30 | 5996.89494 | 6302.99646 | 6684.342166 | 6982.880608 | 7324.560825 | 7690.171475 | 8054.260052 | 8594.545694 | 9104.565413 |
|  | 50 | 40 | 5834.82364 | 6146.999562 | 6534.129279 | 6837.078229 | 7185.474338 | 7562.464708 | 7944.425825 | 8527.126006 | 9097.948745 |
|  | 51 | 50 | 5580.664721 | 5904.473325 | 6297.037731 | 6599.272034 | 6944.298233 | 7318.064809 | 7700.645941 | 8297.557314 | 8902.364784 |
|  | 52 | 60 | 5285.628159 | 5635.240678 | 6039.277294 | 6339.084374 | 6674.493007 | 7035.850839 | 7409.676676 | 8010.935153 | 8650.127677 |
|  | 53 | 70 | 4949.161955 | 5359.391395 | 5793.729217 | 6094.537235 | 6418.452787 | 6763.549439 | 7127.047517 | 7743.254413 | 8453.874495 |
|  | 54 | 80 | 4453.324053 | 5011.395749 | 5519.857851 | 5831.562782 | 6145.943924 | 6476.156955 | 6838.123013 | 7517.928026 | 8427.942311 |
|  | 55 | 90 | 3640.623925 | 4558.728984 | 5230.013402 | 5569.146239 | 5877.853682 | 6197.315422 | 6574.974774 | 7419.548052 | 8848.770322 |
| **Left**  **Caudate** | 56 | 6 | 3086.018579 | 3345.35412 | 3675.46552 | 3932.634552 | 4217.292372 | 4502.157772 | 4759.873281 | 5091.193698 | 5351.889427 |
|  | 57 | 10 | 3029.179945 | 3274.790158 | 3590.480312 | 3838.825151 | 4116.263142 | 4396.701132 | 4652.943302 | 4986.101441 | 5251.354251 |
|  | 58 | 15 | 2960.214583 | 3190.956274 | 3490.633539 | 3729.009354 | 3998.279902 | 4273.916331 | 4529.062579 | 4865.896268 | 5138.515327 |
|  | 59 | 20 | 2899.771146 | 3118.841356 | 3405.752002 | 3636.201865 | 3899.259134 | 4171.951577 | 4427.829294 | 4771.304416 | 5054.489837 |
|  | 60 | 30 | 2803.939825 | 3009.459932 | 3280.821677 | 3501.55643 | 3757.741291 | 4029.427961 | 4291.35277 | 4655.790141 | 4969.207745 |
|  | 61 | 40 | 2710.556764 | 2914.642774 | 3181.473735 | 3397.838057 | 3650.073204 | 3921.051926 | 4187.768019 | 4571.289819 | 4915.653691 |
|  | 62 | 50 | 2612.778202 | 2826.616706 | 3096.991755 | 3311.020101 | 3557.675817 | 3822.650687 | 4086.666388 | 4477.144977 | 4842.992194 |
|  | 63 | 60 | 2525.21406 | 2767.325637 | 3053.683028 | 3268.837875 | 3509.185638 | 3763.811818 | 4018.867707 | 4406.491355 | 4787.018659 |
|  | 64 | 70 | 2405.504903 | 2717.620089 | 3045.32649 | 3269.600484 | 3506.486654 | 3750.699409 | 3996.281252 | 4383.280082 | 4787.537383 |
|  | 65 | 80 | 2136.597197 | 2619.43241 | 3033.377768 | 3276.201666 | 3511.562351 | 3745.777704 | 3985.059543 | 4387.830853 | 4851.956342 |
|  | 66 | 90 | 1649.994031 | 2443.93574 | 3023.081379 | 3298.640279 | 3535.947792 | 3763.903572 | 4007.844426 | 4471.296121 | 5091.826211 |
| **Right**  **Caudate** | 67 | 6 | 3128.821569 | 3413.767022 | 3767.164882 | 4037.553196 | 4334.21322 | 4630.687773 | 4900.582875 | 5252.863919 | 5536.510082 |
|  | 68 | 10 | 3059.710774 | 3327.379391 | 3662.765672 | 3922.033432 | 4209.339917 | 4499.696445 | 4767.078136 | 5120.851171 | 5409.92627 |
|  | 69 | 15 | 2976.937913 | 3225.811075 | 3541.033238 | 3787.563608 | 4064.033319 | 4347.384881 | 4612.249012 | 4969.149 | 5266.752933 |
|  | 70 | 20 | 2910.289834 | 3143.826268 | 3442.273082 | 3678.134591 | 3945.700859 | 4223.865198 | 4488.032001 | 4851.195705 | 5161.015719 |
|  | 71 | 30 | 2826.278697 | 3039.235968 | 3314.391592 | 3535.318763 | 3791.075377 | 4064.445392 | 4332.818812 | 4718.627241 | 5065.793174 |
|  | 72 | 40 | 2736.807133 | 2944.802539 | 3211.308917 | 3424.834436 | 3673.323786 | 3942.626929 | 4212.859537 | 4615.158423 | 4994.281783 |
|  | 73 | 50 | 2641.054411 | 2859.671481 | 3131.93163 | 3345.480461 | 3591.249077 | 3857.07973 | 4125.979269 | 4534.691774 | 4932.525021 |
|  | 74 | 60 | 2541.902622 | 2791.539833 | 3086.743672 | 3308.70151 | 3557.159716 | 3821.431541 | 4087.694385 | 4496.018522 | 4901.611498 |
|  | 75 | 70 | 2404.540368 | 2724.461462 | 3070.036253 | 3311.440678 | 3568.622285 | 3833.075178 | 4095.319572 | 4496.964096 | 4900.054942 |
|  | 76 | 80 | 2135.709499 | 2619.028969 | 3063.248216 | 3338.154484 | 3609.686636 | 3875.410877 | 4133.042687 | 4526.592414 | 4925.922624 |
|  | 77 | 90 | 1718.366693 | 2473.302784 | 3088.287632 | 3414.568729 | 3707.336038 | 3977.975883 | 4235.430509 | 4632.193982 | 5044.75085 |
| **Left**  **Putamen** | 78 | 6 | 5115.166628 | 5467.16292 | 5911.579918 | 6256.124807 | 6637.054346 | 7019.087513 | 7366.547671 | 7817.464285 | 8176.853246 |
|  | 79 | 10 | 4921.592817 | 5270.645301 | 5710.129402 | 6050.253943 | 6426.069755 | 6803.12811 | 7146.539338 | 7593.374153 | 7950.825121 |
|  | 80 | 15 | 4687.429032 | 5032.811771 | 5465.916674 | 5800.235691 | 6169.292914 | 6539.76554 | 6877.835824 | 7319.368414 | 7674.442736 |
|  | 81 | 20 | 4495.104462 | 4838.220579 | 5266.415955 | 5595.91596 | 5959.22866 | 6324.129409 | 6657.8478 | 7095.569954 | 7449.72406 |
|  | 82 | 30 | 4304.947992 | 4652.300096 | 5080.380963 | 5407.084462 | 5766.134773 | 6127.127649 | 6459.0206 | 6898.975478 | 7260.345234 |
|  | 83 | 40 | 4102.78668 | 4457.439365 | 4886.990044 | 5211.030516 | 5565.449449 | 5922.173067 | 6252.418359 | 6696.422736 | 7068.581137 |
|  | 84 | 50 | 3884.370731 | 4248.297244 | 4678.627993 | 4998.015529 | 5344.92857 | 5694.484462 | 6021.063296 | 6468.561712 | 6854.021942 |
|  | 85 | 60 | 3643.158684 | 4026.577918 | 4465.338737 | 4783.721957 | 5126.172006 | 5471.657601 | 5798.422624 | 6257.882655 | 6668.547906 |
|  | 86 | 70 | 3301.211562 | 3735.574233 | 4211.429994 | 4546.359105 | 4901.855444 | 5261.171203 | 5606.864328 | 6110.592119 | 6584.199639 |
|  | 87 | 80 | 2885.501477 | 3428.49468 | 3990.560948 | 4370.259556 | 4766.089416 | 5167.306419 | 5562.629005 | 6167.955306 | 6778.024632 |
|  | 88 | 90 | 2390.384451 | 3125.852579 | 3837.00925 | 4291.758245 | 4754.19648 | 5224.749869 | 5704.029626 | 6490.230663 | 7361.531851 |
| **Right**  **Putamen** | 89 | 6 | 5158.08374 | 5417.335582 | 5764.484922 | 6046.741386 | 6369.446125 | 6706.170308 | 7025.629484 | 7454.953082 | 7805.386052 |
|  | 90 | 10 | 4946.077424 | 5210.732821 | 5563.48034 | 5848.47974 | 6171.954949 | 6509.183797 | 6830.998469 | 7265.819236 | 7622.205812 |
|  | 91 | 15 | 4685.885047 | 4957.140681 | 5316.516203 | 5604.514061 | 5928.339944 | 6265.493176 | 6589.558767 | 7030.283119 | 7393.295496 |
|  | 92 | 20 | 4456.962601 | 4734.209974 | 5099.101929 | 5388.978065 | 5711.688987 | 6047.080265 | 6371.656715 | 6815.73988 | 7183.141549 |
|  | 93 | 30 | 4184.554265 | 4476.896681 | 4855.786403 | 5150.983838 | 5472.563 | 5805.087296 | 6130.924062 | 6581.435665 | 6956.882468 |
|  | 94 | 40 | 3914.538332 | 4227.816641 | 4627.378742 | 4932.51562 | 5257.60852 | 5592.029882 | 5923.897609 | 6387.510919 | 6776.606519 |
|  | 95 | 50 | 3675.742075 | 4005.71465 | 4418.868265 | 4727.50897 | 5048.570344 | 5376.771703 | 5706.370595 | 6171.048368 | 6563.312956 |
|  | 96 | 60 | 3470.241765 | 3814.795079 | 4237.423018 | 4545.77495 | 4858.601376 | 5176.192918 | 5498.883164 | 5957.712787 | 6347.030443 |
|  | 97 | 70 | 3186.251451 | 3568.71887 | 4028.008398 | 4355.195791 | 4678.87946 | 5005.362934 | 5341.194992 | 5822.939924 | 6233.885576 |
|  | 98 | 80 | 2810.202653 | 3278.119213 | 3826.24627 | 4206.549326 | 4572.835478 | 4939.557995 | 5321.245922 | 5873.088933 | 6345.873198 |
|  | 99 | 90 | 2342.490471 | 2967.266576 | 3673.532058 | 4147.472526 | 4590.049794 | 5028.640168 | 5489.536454 | 6159.384652 | 6734.312771 |
| **Left**  **Pallidum** | 100 | 6 | 1454.531934 | 1599.160086 | 1770.478342 | 1898.37257 | 2038.832313 | 2183.010491 | 2321.111062 | 2517.281308 | 2693.947064 |
|  | 101 | 10 | 1360.753253 | 1500.292004 | 1665.264928 | 1788.169456 | 1922.854684 | 2060.729266 | 2192.38412 | 2378.643344 | 2545.58911 |
|  | 102 | 15 | 1252.607627 | 1386.270535 | 1543.86962 | 1660.952004 | 1788.886282 | 1919.388167 | 2043.511805 | 2218.222756 | 2373.891244 |
|  | 103 | 20 | 1174.959709 | 1305.625188 | 1459.217038 | 1572.971536 | 1696.887027 | 1822.825453 | 1942.125521 | 2109.184444 | 2257.154407 |
|  | 104 | 30 | 1119.458401 | 1255.364382 | 1413.937959 | 1530.554856 | 1656.732072 | 1783.971832 | 1903.500402 | 2069.140326 | 2214.124206 |
|  | 105 | 40 | 1057.085588 | 1198.28504 | 1361.498322 | 1480.519058 | 1608.317909 | 1736.109572 | 1855.110614 | 2018.27613 | 2159.414436 |
|  | 106 | 50 | 991.0108789 | 1138.035511 | 1305.988993 | 1427.244334 | 1556.328188 | 1684.233295 | 1802.255833 | 1962.341043 | 2099.191805 |
|  | 107 | 60 | 940.6015351 | 1097.389943 | 1273.862641 | 1399.761354 | 1532.498981 | 1662.737061 | 1781.770832 | 1941.466884 | 2076.392354 |
|  | 108 | 70 | 877.5156353 | 1044.215391 | 1228.399668 | 1357.959968 | 1493.08281 | 1624.262985 | 1742.970793 | 1900.468913 | 2031.994339 |
|  | 109 | 80 | 809.6330663 | 987.9690969 | 1180.575998 | 1313.824725 | 1451.112692 | 1582.882571 | 1700.895301 | 1855.719135 | 1983.523134 |
|  | 110 | 90 | 770.6948892 | 971.4569893 | 1182.58688 | 1325.843285 | 1471.450292 | 1609.503492 | 1731.820359 | 1890.476736 | 2019.950549 |
| **Right**  **Pallidum** | 111 | 6 | 1414.889417 | 1542.07547 | 1684.389797 | 1786.021142 | 1894.462225 | 2003.694389 | 2107.527633 | 2255.38053 | 2390.047039 |
|  | 112 | 10 | 1342.313387 | 1460.157532 | 1593.849646 | 1690.406198 | 1794.297294 | 1899.699331 | 2000.452731 | 2144.649152 | 2276.575162 |
|  | 113 | 15 | 1261.429599 | 1368.861463 | 1492.880721 | 1583.764045 | 1682.638051 | 1783.93064 | 1881.51596 | 2022.215543 | 2151.823758 |
|  | 114 | 20 | 1211.500509 | 1311.319357 | 1428.753517 | 1516.222983 | 1612.604683 | 1712.496105 | 1809.670098 | 1951.135747 | 2082.67308 |
|  | 115 | 30 | 1196.63002 | 1290.722693 | 1405.411355 | 1493.555335 | 1593.16722 | 1698.865149 | 1803.783146 | 1959.710571 | 2107.707862 |
|  | 116 | 40 | 1151.472934 | 1242.978579 | 1356.449188 | 1445.029752 | 1546.446621 | 1655.416266 | 1764.794726 | 1929.302608 | 2087.383548 |
|  | 117 | 50 | 1106.652381 | 1194.879394 | 1304.584033 | 1390.519167 | 1489.317221 | 1596.075821 | 1703.975281 | 1867.83548 | 2027.20796 |
|  | 118 | 60 | 1085.312439 | 1177.268616 | 1288.748696 | 1374.253656 | 1471.137023 | 1574.784841 | 1679.073207 | 1837.540986 | 1992.466087 |
|  | 119 | 70 | 1014.260372 | 1126.5381 | 1252.388558 | 1342.888597 | 1441.020782 | 1542.866754 | 1643.883352 | 1797.248192 | 1948.75733 |
|  | 120 | 80 | 830.1017252 | 1002.945774 | 1165.033735 | 1266.839673 | 1368.61167 | 1469.609205 | 1569.109587 | 1724.440996 | 1886.220144 |
|  | 121 | 90 | 554.4449578 | 862.5301734 | 1113.794211 | 1242.18147 | 1355.612099 | 1462.53852 | 1570.039328 | 1752.708392 | 1966.840124 |
| **Left**  **Hippocampus** | 122 | 6 | 3192.03296 | 3523.246922 | 3843.655417 | 4049.261466 | 4254.585701 | 4452.894293 | 4638.280817 | 4903.402312 | 5149.993729 |
|  | 123 | 10 | 3195.066412 | 3524.541564 | 3846.236393 | 4053.948472 | 4261.844444 | 4462.397901 | 4649.039754 | 4913.693068 | 5157.056855 |
|  | 124 | 15 | 3193.845292 | 3521.723766 | 3845.289594 | 4055.687459 | 4266.795252 | 4470.132789 | 4658.344678 | 4922.539135 | 5162.214817 |
|  | 125 | 20 | 3174.012053 | 3500.841472 | 3826.430472 | 4039.455039 | 4253.630522 | 4459.570871 | 4649.189692 | 4912.799299 | 5148.882483 |
|  | 126 | 30 | 3113.930317 | 3445.829336 | 3781.392327 | 4003.030087 | 4226.412294 | 4440.321677 | 4635.246053 | 4901.322052 | 5133.931507 |
|  | 127 | 40 | 3051.000549 | 3389.710865 | 3736.097105 | 3966.512472 | 4199.040907 | 4420.746109 | 4620.837787 | 4889.499426 | 5119.344416 |
|  | 128 | 50 | 2971.14628 | 3316.954188 | 3673.525898 | 3911.880012 | 4152.469906 | 4380.817909 | 4585.054946 | 4855.213331 | 5081.892022 |
|  | 129 | 60 | 2828.15887 | 3190.821153 | 3565.28118 | 3815.672369 | 4067.80261 | 4305.646388 | 4516.360566 | 4791.058815 | 5017.359496 |
|  | 130 | 70 | 2553.909684 | 2944.985185 | 3344.919466 | 3610.542536 | 3876.275059 | 4124.752637 | 4342.497594 | 4622.134645 | 4848.394625 |
|  | 131 | 80 | 2153.178662 | 2574.494779 | 2996.60652 | 3273.070995 | 3546.773121 | 3799.841028 | 4018.951874 | 4296.101381 | 4516.473992 |
|  | 132 | 90 | 1745.107121 | 2200.461005 | 2645.327395 | 2931.069803 | 3210.177682 | 3464.942235 | 3682.769454 | 3954.239124 | 4166.579161 |
| **Right**  **Hippocampus** | 133 | 6 | 3405.323534 | 3637.86611 | 3897.406012 | 4082.053405 | 4277.935393 | 4473.441756 | 4657.048316 | 4913.997021 | 5143.090079 |
|  | 134 | 10 | 3394.986242 | 3635.252412 | 3901.986403 | 4090.919837 | 4290.671501 | 4489.421528 | 4675.577639 | 4935.385904 | 5166.407873 |
|  | 135 | 15 | 3376.40102 | 3626.557723 | 3902.326049 | 4096.541867 | 4300.979756 | 4503.577071 | 4692.684794 | 4955.686582 | 5188.74108 |
|  | 136 | 20 | 3342.999534 | 3602.510042 | 3886.530105 | 4085.391042 | 4293.781006 | 4499.435986 | 4690.699983 | 4955.690909 | 5189.615336 |
|  | 137 | 30 | 3272.044372 | 3549.773102 | 3849.978472 | 4058.031795 | 4274.21508 | 4485.693709 | 4680.671046 | 4948.059983 | 5181.482321 |
|  | 138 | 40 | 3197.288693 | 3496.685662 | 3815.340835 | 4033.513466 | 4258.058737 | 4475.653343 | 4674.48159 | 4944.402121 | 5177.511804 |
|  | 139 | 50 | 3096.529262 | 3428.821596 | 3771.978517 | 4001.9106 | 4235.29125 | 4459.152331 | 4662.424808 | 4937.360485 | 5174.480115 |
|  | 140 | 60 | 2902.659269 | 3286.058092 | 3659.875052 | 3900.80089 | 4140.119808 | 4367.135424 | 4573.081303 | 4854.012298 | 5100.211718 |
|  | 141 | 70 | 2575.023149 | 3044.253206 | 3459.92501 | 3710.80199 | 3951.830938 | 4177.688215 | 4384.189147 | 4673.778372 | 4938.523429 |
|  | 142 | 80 | 2087.166858 | 2675.695914 | 3144.502228 | 3401.823964 | 3637.479239 | 3855.450608 | 4058.796326 | 4359.574064 | 4655.936804 |
|  | 143 | 90 | 1581.063982 | 2263.414933 | 2797.546553 | 3061.661659 | 3289.163986 | 3497.176888 | 3698.728833 | 4023.841164 | 4381.957269 |
| **Left**  **Amygdala** | 144 | 6 | 1083.018601 | 1204.015933 | 1345.06942 | 1448.431722 | 1559.458385 | 1670.026836 | 1772.141141 | 1910.207422 | 2027.439367 |
|  | 145 | 10 | 1106.628627 | 1225.185701 | 1364.425161 | 1467.178074 | 1578.297347 | 1689.828213 | 1793.702751 | 1935.629554 | 2057.576465 |
|  | 146 | 15 | 1130.226513 | 1245.520175 | 1382.054195 | 1483.643302 | 1594.414034 | 1706.698989 | 1812.419183 | 1958.87454 | 2086.717232 |
|  | 147 | 20 | 1136.995051 | 1248.2629 | 1380.966358 | 1480.444671 | 1589.772943 | 1701.688968 | 1808.239066 | 1957.989491 | 2090.923794 |
|  | 148 | 30 | 1123.398385 | 1230.265302 | 1358.349871 | 1455.044785 | 1562.34084 | 1673.732588 | 1781.675902 | 1937.184307 | 2079.459724 |
|  | 149 | 40 | 1101.182081 | 1212.717444 | 1344.842478 | 1443.779468 | 1553.206569 | 1666.961045 | 1777.85639 | 1939.554071 | 2090.047124 |
|  | 150 | 50 | 1058.369951 | 1181.783469 | 1323.635565 | 1427.284456 | 1540.01259 | 1655.804564 | 1767.983465 | 1931.298574 | 2083.695521 |
|  | 151 | 60 | 965.4344304 | 1113.26344 | 1273.074461 | 1384.28783 | 1501.115657 | 1617.78521 | 1728.552967 | 1887.21265 | 2033.428493 |
|  | 152 | 70 | 773.2579178 | 986.5564213 | 1190.508894 | 1320.03628 | 1448.101748 | 1570.002742 | 1681.784884 | 1837.284581 | 1977.190939 |
|  | 153 | 80 | 486.0997604 | 762.9372632 | 1025.483443 | 1175.77776 | 1313.10323 | 1436.141835 | 1544.405425 | 1690.261567 | 1818.378555 |
|  | 154 | 90 | 350.717871 | 585.8798481 | 864.9347528 | 1027.211404 | 1167.685522 | 1287.017596 | 1388.185224 | 1520.847511 | 1635.291622 |
| **Right**  **Amygdala** | 155 | 6 | 1210.941248 | 1300.118439 | 1413.907702 | 1503.550521 | 1604.75091 | 1709.195175 | 1807.47598 | 1940.880605 | 2052.955364 |
|  | 156 | 10 | 1215.112932 | 1308.367886 | 1426.229212 | 1518.532331 | 1622.572598 | 1730.221908 | 1832.187109 | 1972.261408 | 2091.905222 |
|  | 157 | 15 | 1214.295507 | 1313.542018 | 1436.963403 | 1532.617762 | 1640.07444 | 1751.626863 | 1858.342735 | 2007.711376 | 2138.677548 |
|  | 158 | 20 | 1198.963967 | 1303.737551 | 1431.787563 | 1529.902978 | 1639.706702 | 1754.088322 | 1864.686288 | 2022.66874 | 2165.193979 |
|  | 159 | 30 | 1176.333036 | 1279.577909 | 1407.475906 | 1506.666382 | 1618.780869 | 1736.671904 | 1851.595501 | 2017.123523 | 2167.658439 |
|  | 160 | 40 | 1174.842448 | 1271.06085 | 1395.462237 | 1495.110125 | 1609.833876 | 1731.299198 | 1849.019639 | 2015.098663 | 2161.053145 |
|  | 161 | 50 | 1151.369553 | 1250.574738 | 1379.629124 | 1483.190043 | 1602.053016 | 1726.844573 | 1846.194666 | 2011.114763 | 2152.205336 |
|  | 162 | 60 | 1076.977241 | 1193.143272 | 1337.477981 | 1448.640872 | 1572.045076 | 1697.70501 | 1814.915201 | 1973.198952 | 2106.022288 |
|  | 163 | 70 | 912.7378544 | 1083.548063 | 1265.488694 | 1390.335736 | 1519.067559 | 1643.940748 | 1758.026289 | 1912.682842 | 2045.926155 |
|  | 164 | 80 | 557.2242869 | 866.5992107 | 1137.260692 | 1284.413213 | 1417.415506 | 1540.289733 | 1656.337818 | 1832.714197 | 2013.019998 |
|  | 165 | 90 | 375.6423394 | 665.1482319 | 1003.786131 | 1174.298522 | 1308.234264 | 1432.599264 | 1574.438394 | 1881.880392 | 2341.098861 |
| **Left**  **Nucleus**  **Accumbens** | 166 | 6 | 497.9355322 | 543.8842851 | 608.8664247 | 664.8825546 | 732.7345011 | 807.0868525 | 880.1303222 | 982.3468014 | 1069.479548 |
|  | 167 | 10 | 460.6164344 | 505.5428009 | 568.7646679 | 623.0028975 | 688.4257116 | 759.8271641 | 829.7344596 | 927.2631165 | 1010.203527 |
|  | 168 | 15 | 419.3481465 | 463.6740272 | 525.4411668 | 577.9737559 | 640.9162187 | 709.2422683 | 775.9166889 | 868.8097429 | 947.8855279 |
|  | 169 | 20 | 388.8887849 | 434.3118218 | 496.5309261 | 548.7089293 | 610.63514 | 677.4649896 | 742.5957466 | 833.6887 | 911.9388769 |
|  | 170 | 30 | 352.456032 | 405.760535 | 474.5615799 | 529.6142131 | 593.0932573 | 660.6653376 | 726.770418 | 821.4876372 | 906.4556302 |
|  | 171 | 40 | 313.1964837 | 371.6514849 | 444.3167328 | 500.6521369 | 564.1905699 | 630.7914336 | 695.4871894 | 788.2582942 | 872.1845604 |
|  | 172 | 50 | 285.9677343 | 342.7363299 | 414.5903765 | 470.7726703 | 533.9056577 | 598.9661817 | 660.3299599 | 744.21395 | 815.4920325 |
|  | 173 | 60 | 255.6383801 | 314.5821178 | 389.9797523 | 449.0770298 | 515.0094027 | 581.7332646 | 642.9353055 | 723.1023961 | 787.6112271 |
|  | 174 | 70 | 194.1408832 | 261.5937435 | 346.0183587 | 410.8751122 | 481.8811823 | 552.2088639 | 615.2569457 | 695.5562277 | 758.1659176 |
|  | 175 | 80 | 110.6678365 | 189.7155307 | 286.6122592 | 358.2094607 | 434.1925762 | 507.3519122 | 571.3628401 | 650.89285 | 711.4172584 |
|  | 176 | 90 | 54.2953935 | 127.2445293 | 234.5591702 | 313.688571 | 395.0490775 | 470.9554487 | 535.6830354 | 614.2030371 | 672.6932235 |
| **Right**  **Nucleus**  **Accumbens** | 177 | 6 | 529.2730042 | 576.3561206 | 639.835888 | 691.6159882 | 750.8096493 | 813.0210155 | 872.8943697 | 954.448558 | 1021.754033 |
|  | 178 | 10 | 502.6840424 | 549.6651341 | 612.8583539 | 664.2359621 | 722.7458771 | 784.1711427 | 843.3977911 | 924.1924992 | 990.931091 |
|  | 179 | 15 | 471.0453143 | 517.9313115 | 580.8046173 | 631.7072561 | 689.3941621 | 749.8668213 | 808.3063186 | 888.1698658 | 954.2072736 |
|  | 180 | 20 | 443.0801108 | 490.0911868 | 552.942645 | 603.6152519 | 660.7633654 | 720.5866732 | 778.5328969 | 857.8689612 | 923.542 |
|  | 181 | 30 | 402.5961083 | 451.1755019 | 515.7729052 | 567.4488927 | 625.1942064 | 685.5075826 | 744.2363907 | 824.993747 | 892.0330717 |
|  | 182 | 40 | 367.3395952 | 418.3243721 | 485.6617408 | 539.0376519 | 598.0617102 | 659.500224 | 719.5761037 | 802.4431534 | 871.3445087 |
|  | 183 | 50 | 329.8004061 | 384.4735758 | 455.6815173 | 511.2362441 | 571.6865059 | 634.052609 | 694.9839535 | 778.826174 | 848.2704222 |
|  | 184 | 60 | 288.0760998 | 348.6706669 | 425.5528619 | 483.9897918 | 546.0753837 | 609.122697 | 670.2949528 | 753.7285825 | 822.1546626 |
|  | 185 | 70 | 236.9919819 | 306.4633455 | 390.6522703 | 452.1442806 | 515.3792782 | 578.1554368 | 638.3199268 | 719.2369259 | 784.6511621 |
|  | 186 | 80 | 173.8973594 | 257.0604975 | 351.1176132 | 415.8745757 | 479.6304671 | 541.0437975 | 598.8716922 | 675.2086037 | 735.8183122 |
|  | 187 | 90 | 116.3025037 | 209.7914938 | 316.1223775 | 385.156562 | 449.8743212 | 510.1520979 | 565.7635253 | 637.6802508 | 693.7059895 |
